# Brain PDGFRβ^+^ cells exhibit diverse reactive phenotypes after stroke without requiring KLF4

**DOI:** 10.64898/2026.04.09.712632

**Authors:** Daniel Manrique-Castano, Ayman ElAli

**Author notes:** **Correspondence**, Dr Ayman ElAli, Department of Psychiatry and Neuroscience, Faculty of Medicine, Université Laval, Neuroscience axis (Office T2-39), Research center of CHU de Québec-Université Laval (CHUL), 2705 Boulevard Laurier, Quebec City, G1V 4G2, QC, Canada. **Data availability statement** All the data associated with the publication are made fully available via https://zenodo.org/records/10553084, which includes original microscopy images stored in Zenodo (10.5281/zenodo.10553084) as well as access to complementary data stored in Open Science Framework (OSF) repository (10.17605/OSF.IO/74MQN), and reproducible workflows for image processing and analysis are available on GitHub (10.5281/zenodo.13948420). **Author contributions** Ayman ElAli and Daniel Manrique-Castano designed the study. Daniel Manrique-Castano performed the experiments and prepared the figures. Ayman ElAli and Daniel Manrique-Castano validated the analysis and interpreted the results. Ayman ElAli and Daniel Manrique-Castano drafted and edited the manuscript. **Ethics approval statement** Animal studies were performed according to the Canadian Council on Animal Care guidelines, as implemented by the institutional animal ethics committee (*Comité de Protection des Animaux de l’Université Laval-3* (CPAUL-3); Protocol # # 20-470) and were reported according to ARRIVE 2.0 guidelines.

## Abstract

Ischemic stroke triggers a cascade of molecular and cellular processes leading to fibrotic scar formation, entailing activation of brain platelet-derived growth factor receptor (PDGFR)β^+^ cells. Krüppel-like factor (KLF)4 plays an important role in regulating the activation of peripheral PDGFRβ^+^ perivascular cells in response to hypoxia/ischemia. Herein, we aimed to characterize the spatiotemporal responses of brain PDGFRβ^+^ cells while assessing the contribution of KLF4. This was achieved using transgenic mice that enable tracking or conditionally depleting KLF4 in PDGFRβ^+^ cells, which were subjected to experimental ischemic stroke. Next, we employed point pattern analysis (PPA) and topological data analysis (TDA) to quantitatively characterize cell phenotypic changes and spatial distribution over injury progression after ischemic stroke. We show that brain PDGFRβ^+^ cells rapidly become reactive and early localize to regions prone to irreversible damage. We report the emergence of parenchymal PDGFRβ^+^ cells, which cannot be causally linked to proliferation or vascular detachment. Moreover, our analysis reveals that KLF4 is barely expressed in brain PDGFRβ^+^ cells under normal conditions, and that its expression is slightly induced in reactive cells in the injured brain. Notably, specific attenuation of KLF4 induced expression in PDGFRβ^+^ cells does not affect cell reactivity and spatiotemporal distribution, nor scar formation and injury severity. These observations suggest that in contrast with the periphery, KLF4 is not implicated in regulating the responses of brain PDGFRβ^+^ cells. Our results indicate that the reactivity of brain PDGFRβ^+^ cells after stroke is spatiotemporally diverse, evolve over injury progression, and is distinct from peripheral perivascular cells.

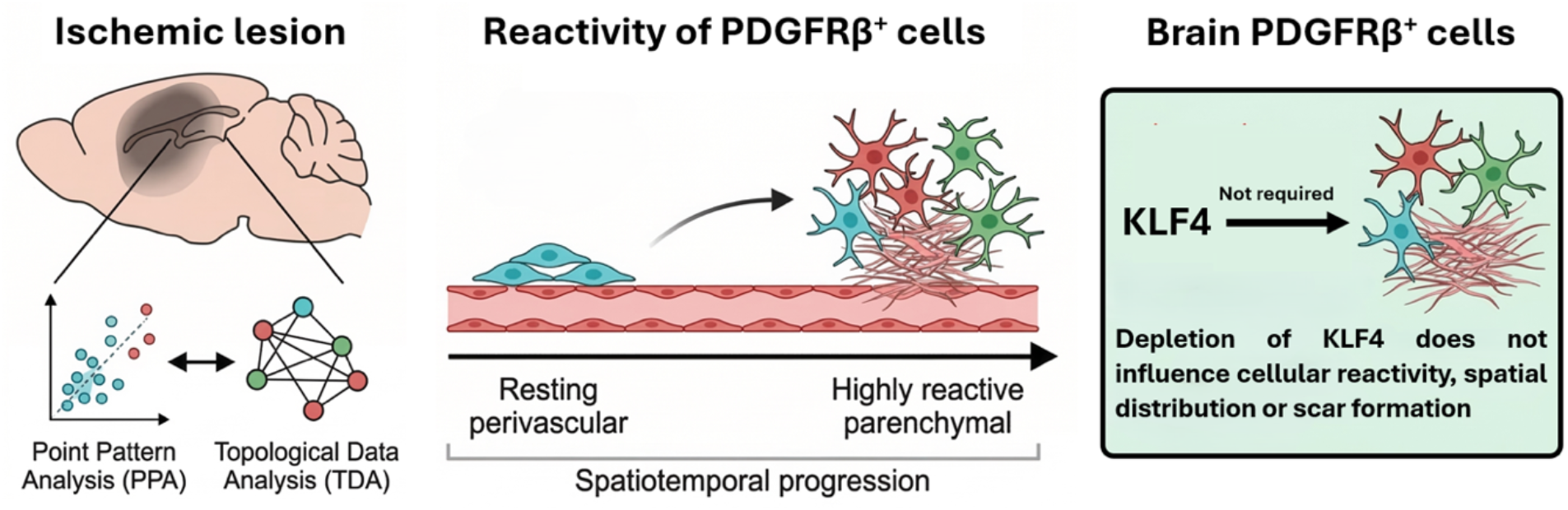

## 1. Introduction

Brain injury upon ischemic stroke includes an infarct core comprising irreversibly lost tissue, surrounded by a peri-lesional region comprising salvageable tissue (Lo, 2008). Upon initial insult associated with cell death, subacute and chronic molecular and cellular responses are initiated as an attempt to promote repair, including neuroinflammation and scar formation, which decisively affect definitive lesion maturation and neurological outcomes (Sofroniew, 2009). Tissue scar is a finely organized multicellular structure that is conserved across different central nervous system (CNS) injuries (Goritz et al., 2011; Sofroniew, 2009; R. Zhang et al., 2018). It comprises a neatly demarcated glial scar formed essentially by reactive astrocytes and activated microglia (Zamanian et al., 2012; R. Zhang et al., 2018), surrounding a fibrotic scar that is mainly formed by fibrotic-like cells and macrophages, including platelet-derived growth factor receptor (PDGFR)β^+^ cells (Fernandez-Klett et al., 2013). Although scar formation represents an important step in neuroinflammation resolution, whether it helps or impedes brain regeneration remains debatable (Adams & Gallo, 2018; Anderson et al., 2016; Bradbury & Burnside, 2019; H. Wang et al., 2018). Glial and fibrotic scar formation is a complex progressive process that implicates narrowly regulated spatiotemporal reorganization of reactive cells. The current findings are indicating that the cells involved in fibrotic scar formation provide cues to enable an adequate organization of glial scar (Shibahara et al., 2020, 2023). Yet, a comprehensive spatiotemporal analysis of cellular reorganization underlying scar formation relative to injury progression after stroke is still lacking.

Resident brain perivascular cells ubiquitously express PDGFRβ, as well as divergent markers under normal conditions such as CD13 in pericytes and α-smooth muscle actin (SMA) in vascular smooth cells (VSMCs) (He et al., 2016). PDGFRβ^+^ cells are among the first responders to injury (Duan et al., 2018), undergo profound morphological changes, and acquire new cellular functions presumably through reprogramming (Dias et al., 2021). Ischemic stroke induces a rapid loss of CD13^+^ pericytes and the emergence of PDGFRβ^+^ cells exhibiting stromal characteristics that partake in fibrotic scar formation via the release of extracellular matrix (ECM) proteins (Fernandez-Klett et al., 2013; Shen et al., 2012; Shibahara et al., 2023). It is proposed that upon injury, reactive PDGFRβ^+^ cells detach from adjacent vasculature, proliferate and migrate towards the infarct core to contribute to the fibrotic reaction (Dias et al., 2021; Fernandez-Klett et al., 2013). Reactive PDGFRβ^+^ cells have been reported to actively regulate the inflammatory responses (Rustenhoven et al., 2017) and to orchestrate the spatial organization of glial scar (Dias et al., 2021; Shibahara et al., 2020, 2023). Interestingly, extent of PDGFRβ^+^ cell reactivity is narrowly dependent upon severity of brain injury upon stroke (Dias et al., 2021). Yet, the temporal phenotypic changes of reactive PDGFRβ^+^ cells and spatial reorganization over injury progression, as well as the mechanisms underlying activation remain unexplored. A deeper understanding of these aspects will enable providing novel insights into the dynamics underpinning scar organization after ischemic stroke.

Krüppel-like factor (KLF4) is a zinc-finger transcription factor that regulates key biological processes, including reprogramming, dedifferentiation, differentiation, proliferation, and migration (Ghaleb & Yang, 2017). In lung fibrosis, KLF4 expression is induced in reactive PDGFRβ^+^ cells that are transitioning towards fibrotic-like cells, leading to the activation of transforming growth factor (TGF)β pathway and the release of ECM proteins (Chandran et al., 2021). Hypoxia associated with pulmonary hypertension promotes PDGFRβ^+^ cell dedifferentiation, migration, and expansion via KLF4 action (Sheikh et al., 2015). In atherosclerotic lesions, KLF4 regulates PDGFRβ^+^ perivascular cell reprogramming and subsequent transition into mesenchymal stem cell-like cells (Shankman et al., 2015a). Moreover, KLF4 is reported to maintain the functions of PDGFRβ^+^ perivascular cells within the microvascular bed upon cardiac ischemia (Haskins et al., 2018). Finally, KLF4 induces a dedifferentiated state in pericytes within PDGFRβ^+^ perivascular metastatic niches (Murgai et al., 2017). These observations outline major roles of KLF4 in shaping the responses of PDGFRβ^+^ cells upon hypoxic/ischemic insults in peripheral organs. These roles were proposed to be conserved in all PDGFRβ^+^ cells. However, in contrast to peripheral organs, PDGFRβ^+^ cells in the brain originate essentially from the neuroectoderm (Korn et al., 2002a). Therefore, whether KLF4 fulfills similar roles in regulating the reactivity of brain PDGFRβ^+^ cells and subsequent contribution to fibrotic scar organization after ischemic stroke remain unknown.

Herein, we aimed to analyze the spatiotemporal phenotypic changes and reorganization of reactive PDGFRβ^+^ cells in experimental ischemic stroke while elucidating the contribution of the regulatory functions of KLF4. For this purpose, transgenic mice in which PDGFRβ^+^ cells are tracked via expression of tdTomato, PDGFRβ^tdTomato^ mice, and mice in which KLF4 is conditionally depleted in PDGFRβ^+^ cells, PDGFRβ^KLF4-KO^ mice, were subjected to transient middle cerebral artery occlusion (MCAo). We employed novel unbiased and automated workflows incorporating point pattern analysis (PPA) and topological data analysis (TDA) (Manrique-Castano et al., 2024) to analyze the morphological changes, spatial distribution, and interaction of reactive PDGFRβ^+^ cells with neighboring cells over injury progression. Our results provide a model-based quantitative characterization of the spatiotemporal reorganization of reactive PDGFRβ^+^ cells within the neurovascular interface after ischemic stroke. Using PDGFRβ^tdTomato^ reporter mice, we show that reactive brain PDGFRβ^+^ cells exhibit predictable spatial distribution within the injured brain and are early localized to the tissue that is prone to be irreversibly lost over time. Notably, reactive PDGFRβ^+^ cells do not show a significant proliferative activity in our reporter mice. Using PDGFRβ^KLF4-KO^ mice, our results indicate that KLF4 is not required to regulate the response of brain PDGFRβ^+^ cells to ischemic stroke. The conditional specific depletion of KLF4 in PDGFRβ^+^ cells does not affect neither the morphology nor the spatiotemporal reorganization of reactive cells and does not affect injury progression after stroke. Furthermore, the contribution of reactive PDGFRβ^+^ cells to the fibrotic reaction translated by the release of ECM proteins as well as glial scar organization after stroke remain unchanged in PDGFRβ^KLF4-KO^ mice. Our study provides novel insights into the spatiotemporal dynamics of brain PDGFRβ^+^ cells over injury progression after stroke and challenges the current proposed paradigms for the mechanisms governing cell reactivity upon hypoxic/ischemic CNS injuries.

## 2. Materials and Methods

All raw data and analysis pipelines are publicly available for validation and reuse. This article comprises a Zenodo repository hosting raw widefield and confocal microscopy images (Manrique-Castano & ElAli, 2024c) and an Open Science Framework (OSF) repository (Manrique-Castano & ElAli, 2024a) hosting a wide variety of resources, including low-resolution images (.png), raw and processed datasets (.csv), point patterns (.RDS), point clouds (.npy), statistical models (.RDS), and statistical model summary tables (.tex/html). The OSF repository also comprises the pipelines used for image pre-processing and analysis derived from FIJI (.ijm) (Schindelin et al., 2012a), CellProfiler (.cpproj) (Stirling et al., 2021a), Ilastik (.ilp) (Berg et al., 2019a), and QuPath (.groovy) (Bankhead et al., 2017a). To ensure full reproducibility and reuse of data/results, we host fully annotated [Quarto](https://quarto.org/) (QN) (Allaire et al., 2022) and [Jupyter](https://jupyter.org/) (JN) notebooks encompassing data processing and statistical modeling in a versioned GitHub repository (Manrique-Castano & ElAli, 2024b). Additionally, we provide supplementary methods detailing the implemented analysis pipelines available in a repository in [GitHub](https://github.com/elalilab/Stroke_PDGFR-B_Reactivity/blob/main/PDGFR-B_Reactivity_SupplementaryTables.qmd). Note that the main article and the supplementary methods have direct links to specific Zenodo, OSF, or GitHub resources. While some resources are rendered in their respective platform, others need to be processed in the corresponding software. In all cases, our approach uses open-source/free software. Our experimental design and analysis workflow are summarized in **Figure 1B**.

**Figure 1.**
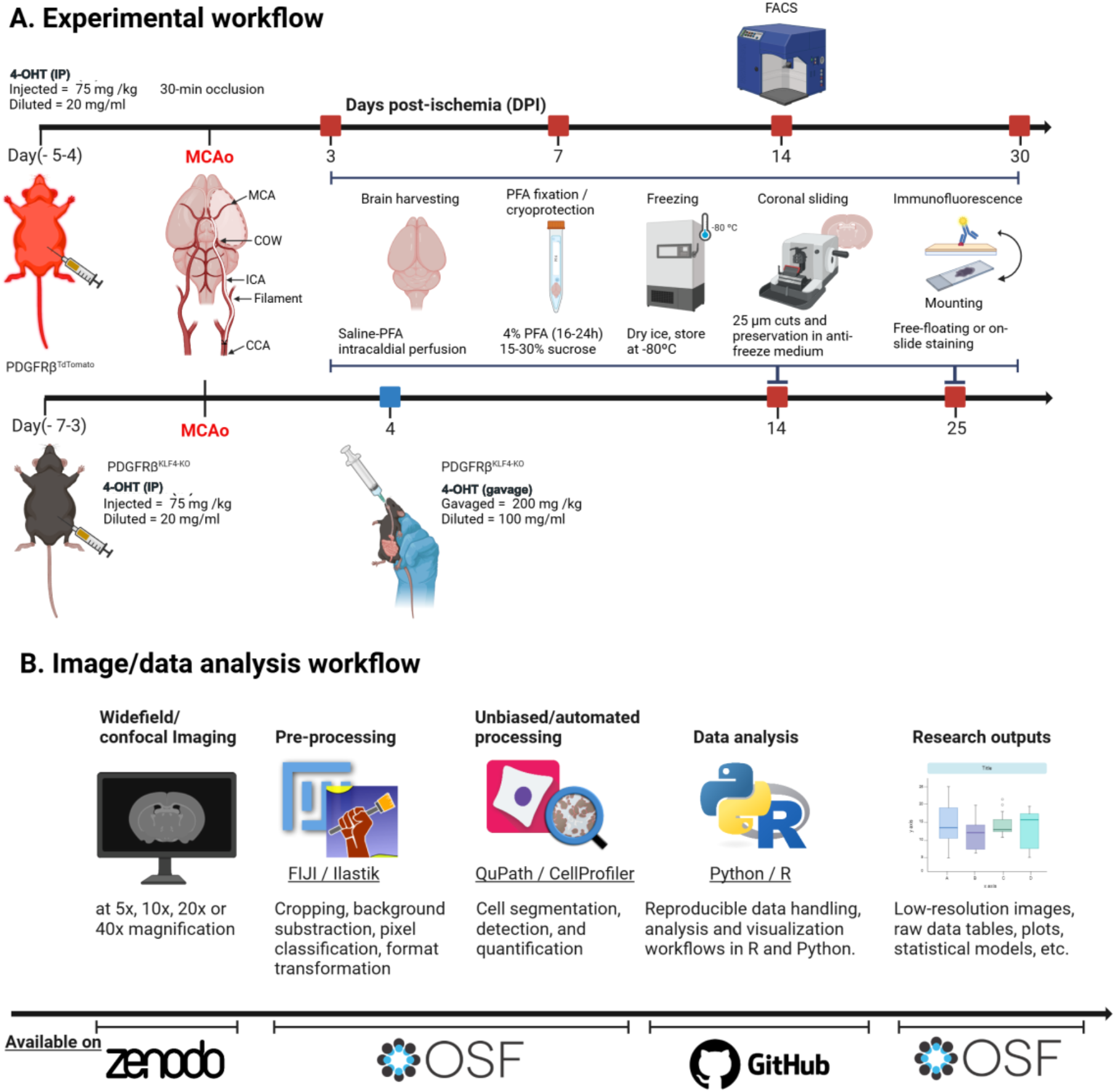
Experimental design and analysis workflows. **A)** tdTomato was induced in PDGFRβ^+^ cells via 4-hydroxytamoxifen (4-OHT) intraperitoneal (IP) administration in B6.Cg-*Gt(ROSA)26Sor*^tm14(CAGtdTomato)Hze^/J;B6.Cg-Pdgfrb^tm1.1(cre/ERT2)Csln^/J mice [PDGFRβ^tdTomato^ reporter mice]. KLF4 was specifically depleted in PDGFRβ^+^ cells either via tamoxifen IP (initiated at day 7 prior to ischemic stroke onset) or gavage (initiated at day 4 after ischemic stroke onset) administration in B6.129S6-Klf4^tm1khk^/Mmmh;B6.Cg-Pdgfrb^tm1.1(cre/ERT2)Csln^/J mice [PDGFRβ^KLF4-KO^ mice]. Both genotypes, along with corresponding control or sham animals, underwent a 30-minute middle cerebral artery occlusion (MCAo) using a nylon-coated monofilament (CCA: Common carotid artery; ICA: Internal carotid artery; COW: Circle of Willis; MCA: Middle cerebral artery). Brains were harvested from PDGFRβ^tdTomato^ mice at 3, 7, 14, and 30 days post-ischemic stroke (DPI) and processed for immunofluorescence (on-slide or free-floating) or for fluorescence-activated cell sorting (FACS) at 14 DPI. For the PDGFRβ^KLF4-KO^ animals, brains were harvested at 14 DPI for post-ischemic stroke onset KLF4 depletion and at 25 DPI for pre-ischemic stroke onset KLF4 depletion. Consistent protocols were applied for immunofluorescence across genotypes. **B)** A schematic illustration of the image and data analysis workflow using open-source software. Upon brain section imaging, preprocessing was performed using FIJI and/or Ilastik, followed by unbiased and automated cell detection with CellProfiler or QuPath. The results were analyzed using R or Python code. All raw images, data, and research outputs are available in our Zenodo (Manrique-Castano & ElAli, 2024c), OSF (ElAli & Manrique-Castano, 2024), or GitHub (Manrique-Castaño & Interactions, 2024) repositories, as depicted. The figure was created using BioRender https://BioRender.com/v11z645. High-resolution figure: https://osf.io/jxb4z/files/y7csx.

### 2.1. Animals

Three to five months old C57BL6/J male mice were obtained from The Jackson Laboratory (Bar Harbor, ME, USA). Age-matching transgenic mice heterozygous for B6.Cg-PDGFRβ^tm1.1(cre/ERT2)Csln^/J (PDGFRβ^CreERT2^; #030201, The Jackson Laboratory) were crossbred with either homozygous mice B6.Cg-Gt(Rosa)26Sor^tm9(CAG-tdTomato)Hze^ (Ai9; #007909, The Jackson Laboratory) to generate PDGFRβ^tdTomato^ reporter mice, or homozygous for B6.129S6-Klf4^tm1khk^/Mmmh (KLF4^Flox^; RRID:MMRRC_029877-MU, Mutant Mouse Resource & Research Centers (MMRRC), MO, USA) (PMID: 12015290; PMID: 15825076) to generate PDGFRβ^KLF4-KO^ mice. In PDGFRβ^CreERT2^:Ai9 mice a *loxP*-flanked STOP cassette, which prevents the transcription of a CAG promoter-driven tdTomato, is inserted into Gt(ROSA)26Sor locus. Cre-recombinase was induced via administration of 4-hydroxytamoxifen (4-OHT) (80 mg/kg in corn oil; Intraperitoneal; 1x daily for 2 days) prior to experimental ischemic stroke to excise the flanked *loxP* sites, allowing tdTomato specific expression in PDGFRβ^+^ cells. In PDGFRβ^CreERT2^:KLF4^Flox^ mice, exons 2 and 3 of the *Klf4* gene was flanked with *loxP* sites in introns 1 and 3. In some experiments, Cre-recombinase was induced via administration of 4-hydroxytamoxifen (4-OHT) (200 mg/kg in corn oil; Oral gavage; 1x daily for 3 consecutive days) after experimental ischemic stroke (post-onset depletion), and in other experiments via administration of 4-OHT (80 mg/kg in corn oil; Intraperitoneal; 1x daily for 5 consecutive days) prior to experimental ischemic stroke (pre-onset depletion), to excise the flanked *loxP* sites, leading to KLF4 specific depletion in PDGFRβ^+^ cells **(Figure 1)**.

All mouse genotypes were confirmed by PCR using DNA isolated from ear punches. The following probes were used; For PDGFRβ^CreERT2^: Common: 5’ – CAC AAC TGA AGT AAG TTC CAC C – 3’; Wild type reverse: 5’ – GTC GAT GTG ACG GTGT TTC GA – 3’; Mutant reverse: 5’ – CGT TCT TGG ACT ACC TGT ACA – 3’. For Ai9: oIMR9020: Wild type forward: 5’ – AAG GGA GCT GCA GTG GAG TA – 3’; oIMR9021: Wild type reverse: 5’ – CCG AAA ATC TGT GGG AAG TC – 3’; oIMR9103: Mutant reverse: 5’ – GGC ATT AAA GCA GCG TAT CC – 3’; oIMR9105: Mutant forward: 5’ – CTG TTC CTG TAC GGC ATG G – 3’. For KLF4^Flox^: Exon 1: 5’-CTG GGC CCC CAC ATT AAT GAG-3’; Exon 2: 5’-CGC TGA CAG CCA TGT CAG ACT-3’ (PMID: 12015290; PMID: 15825076). All animal procedures and handling were performed according to the Canadian Council on Animal Care guidelines, as implemented by *Comité de Protection des Animaux de l’Université Laval-3* (CPAUL-3; Protocol # 20-470). Animal studies were reported according to ARRIVE 2.0 guidelines.

### 2.2. Experimental ischemic stroke

Mice were subjected to focal ischemic stroke via the transient middle cerebral artery occlusion (MCAo) using an intraluminal filament technique (Manrique-Castano et al., 2024). Mice were anesthetized with 3 % isoflurane for induction and 1.5 % for maintenance (95 % O2, 2 l/min) and body temperature was maintained between 36 and 37 °C using a feedback-controlled heating system (Harvard Apparatus®, QC, Canada). A midline neck incision was performed, and the left common artery (CCA) was exposed under a surgical microscope and ligated. A 7-0 silicon-coated nylon monofilament (MCAO suture, Doccol Corporation, Cat# 7022910PK5Re) was inserted through the internal carotid artery (ICA) until the origin of MCA. The monofilament was left in place for 30 minutes and then withdrawn to mediate reperfusion **(Figure 1)**. This model generates a focal injury in the striatum and overlaying cortex as is associated with the formation of neatly demarcated astroglial and fibrotic scar. Brains were harvested at 3, 7, 14, and 30 days post-ischemic stroke (DPI) for further analysis **(Figure 1)**. C57BL6/J, PDGFRβ^CreERT2^, KLF4^Flox^, and Ai9 sham-operated animals with similar 4-OHT regimen were generated and used whenever required as controls.

### 2.3. Brain sectioning and immunolabeling

Mice were euthanized mice via transcardial perfusion with ice-cold phosphate-buffered saline (PBS) followed by 4% paraformaldehyde (PFA). Brains were post-fixed at 4 °C by immersion in 4% paraformaldehyde (PFA) for 16– to 24 hours. Cryoprotection was achieved through sequential immersion in 15% and then 30% sucrose in PBS, and were subsequently frozen using dry ice and stored at -80 °C until sectioning using a freezing microtome. We performed 25 μm-thick coronal sectioning and conducted staining either on-slide or free-floating, depending on experimental requirements. We share the detailed staining protocols and corresponding antibodies/reagents in the “Protocols” component of the OSF repository.

### 2.4. Fluorescent in situ hybridization (FISH)

Fluorescence *in situ* hybridization (FISH) was performed to assess PDGFRβ mRNA expressing using the RNAscope^®^ Fluorescent Multiplex Reagent Kit (ACDBIO, Cat #323100), strictly following the manufacturer’s protocol. Free-floating brain sections were mounted onto the Superfrost® Plus slide and kept at -20 °C to dry for 1 hour before being baked for 30 minutes at 60 °C. The sections were fixed for 15 minutes in 4% PFA and washed with 0.1M PBS. Sections were next dehydrated using ethanol and incubated in hydrogen peroxidase (H_2_O_2_) for 10 minutes. Following retrieval, sections were immersed in a 100% ethanol bath, and a hydrophobic barrier was drawn, and 5 drops of Protease III cocktail were added to completely cover the sections for 40 minutes at room temperature (RT). Following series of washes with 0.1M PBS, 4 drops of mouse RNAscope^®^ Probe-*PDGFRβ*-C1 (411381) were applied to entirely cover sections, which were immediately placed in the humidity control tray of the HybEZ™ Oven for 2 hours at 40 °C. Sections were gently removed, liquid excess was removed and were next rinsed with 1X wash buffer. For hybridization, 4 drops of Amp-1 were added to entirely cover the entire sections, and were incubated in the HybEZ™ Oven for 30 minutes at 40 °C. This process was repeated with Amp-2 for 30 minutes and Amp-3 for 15 minutes. PDGFRβ mRNA transcripts were detected using Opal™ 690 (1/700) and 4 drops of DAPI (provided in the kit) were added to sections and kept for 30 seconds, followed by a series of washes with 1X wash buffer. Slides were next mounted with 110 μL of Fluoromount-G^®^ anti-fade medium, carefully cover-slipped, and placed at 4 °^°^C in the dark until analysis. Epifluorescence images were acquired using Axio Observer microscope equipped with a module for optical sectioning (Apotome.2) and Axiocam 503 monochrome camera and processed using ZEN Imaging Software (Carl Zeiss Canada, ON, Canada).

### 2.5. Brain section imaging

Brain sections were imaged using either widefield Zeiss Axio Observer or laser scan confocal Zeiss (LSM800) microscope with adequate objectives. Due to strong recombination activity, tdTomato signal leaked into the green channel (488 nm) when using widefield microscopy. Consequently, we exclusively used laser scan confocal microscopy to image the green channel, employing appropriate excitation wavelengths and detector ranges (see corresponding *readme* files). We share widefield images in .tif or .czi formats to encourage reuse and validation of our image sets. Technical metadata, including imaging parameters and *readme* files, are incorporated in the .zip files. The “[Images_Low-Resolution](https://osf.io/mb257/)” component of the OSF repository contains complete sets of low-resolution images used for illustration. Brain sections were consistently imaged at the level of the MCA territory (bregma 0.44 to -0.06 mm) **(Figure 1A)** to acquire images of whole-brain, ipsilateral, or region of interests (ROIs) at different magnifications, as specified in supplementary methods and results sections. To facilitate cell detection, we used tailored batch-processing FIJI (ImageJ v1.54f) scripts (.ijm) to handle and enhance images (Schindelin et al., 2012b) The scripts are available in each .zip file at the Zenodo repositories and in the “[Images_Processing](https://osf.io/9zdc7/)” component of the OSF repository.

### 2.6. Unbiased cell detection, quantification, and colocalization

Unbiased and automated cell detection, quantification, and colocalization analysis was applied using [QuPath](https://qupath.github.io/) (v 0.4.4) (Bankhead et al., 2017b) and [CellProfiler](https://cellprofiler.org/) (v.4.2.4) (Schindelin et al., 2012a; Stirling et al., 2021b). When necessary, machine learning-based pixel classification was employed to obtain segmentation maps using [Ilastik](https://www.ilastik.org/) (Berg et al., 2019b). Herein, we provide detailed descriptions for each image set in the supplementary methods and share the pipeline files/scripts in the “[Images_Processing](https://osf.io/9zdc7/)” component of the OSF repository to encourage validation and ensure full reproducibility of our results.

### 2.7. Morphological analysis of PDGFRβ^+^ cells and principal component analysis

Z-stack [images](https://osf.io/mfyge) of PDGFRβ^tdTtomato^/CD31- and CD13-immunolabeled sections were acquired using laser scan confocal microscope at 40x (see immunolabeling [protocol](https://osf.io/d2un3/files/9mgyh)) to analyze the morphology and spatial location of PDGFRβ^tdTomato^ cells in defined ROIs, specifically, the injured cortex, the injured striatum, and the healthy cortical peri-lesional regions **(see** [QN](https://github.com/elalilab/Stroke_PDGFR-B_Reactivity/blob/main/Confocal_40x_ROIs_Pdgfrb_Morpho_BatchScript.qmd); **Suppl. Method 10)**. Using [FIJI](https://osf.io/tns7r), z-projection images were generated and manually segmented into one or more well-identified cells per ROI. Next, Python libraries skimage (Walt et al., 2014) and scipy (Virtanen et al., 2020) were used to enhance, threshold, and generate [masks](https://osf.io/h2uxq) of individual cells. The various objects were imported to CellProfiler to perform a user-defined classification into i) amoeboid, ii) perivascular, iii) reticuloparenchymal, iv) reticulite morphological categories. Subsequently, the classification results were imported into R to conduct principal component analysis (PCA) using the FactoMineR and factoextra packages **(see Suppl. Method 10)**.

### 2.8. Fluorescence-activated cell sorting (FACS)

Brain PDGFRβ^tdTomato^ cells from PDGFRβ^tdTomato^ reporter mice were isolated via FACS at 14 DPI using a modified protocol of the adult brain dissociation kit (Miltenyi Biotec, cat # 130-107-677), as detailed in ([Protocol](https://osf.io/mdnzy); **see Suppl. Method 8)**. Data was visualized and analyzed using open-source Bioconductor packages including flowCore (Hahne et al., 2009) and ggcyto (Van et al., 2018). Additionally, quality checks were performed using the flowAI package (Monaco et al., 2016). The original “.fcs files”, along with compensation controls and quality checks files, are available in the “[Datasets/FACS](https://osf.io/zvnxa/)” component of the OSF repository. FACS files were processed using quarto [notebook](https://github.com/elalilab/Stroke_PDGFR-B_Reactivity/blob/main/FlowCytometry_Pdgfrb_Processing.qmd) running R version 4.4, while the [modeling](https://github.com/elalilab/Stroke_PDGFR-B_Reactivity/blob/main/FlowCytometry_Pdgfrb_Analysis.qmd) with the brms package was conducted in R 4.1.2.

### 2.9. Data analysis

We analyzed the data using open-source notebooks and scripts in [Quarto](https://quarto.org/) (QN) (R language), and [Jupyter](https://jupyter.org/) (Python language) both within the [R-studio](https://posit.co/) IDE environment. The analysis strategy involved i) generating Bayesian posterior distributions via Markov chain Monte Carlo (MCMC) methods (Kruschke & Liddell, 2018) using the [brms](https://paul-buerkner.github.io/brms/) package (Bürkner, 2017); ii) performing point pattern analysis (PPA) with the [spatstat](https://github.com/spatstat/spatstat) package (Baddeley & Turner, 2005); iii) conducting topological data analysis (Bhaskar et al., 2021a) using [Ghudi](https://gudhi.inria.fr/python/latest/) (The GUDHI Project, 2023) and [Ripser](https://ripser.scikit-tda.org/en/latest/) (Bauer, 2021); and iv) applying bootstrapping techniques (Chernik, 2007a) using [SciPy](https://scipy.org/) (Virtanen et al., 2020). The versions of the packages and libraries are indicated in the *sessionInfo* of each notebook. We performed scientific inference based on parameter and coefficient values and their uncertainty. To encourage validation and reuse of our analysis pipelines and results, we share fully annotated QNs and JNs in our GitHub repository. These resources contain tailored approaches, including raw data handling, exploratory data analysis, statistical modeling, model comparison and diagnostics, and analysis and visualization of results. Below, we specify common aspects of our data analysis approaches.

The primary approach for data analysis involves MCMC-based Bayesian modeling using the brms package (Bürkner, 2017). We explored the parameter space and derived stable posterior distributions using the following standard arguments: *seed = 8807, chains = 4, cores = 4, warmup = 2500, iter = 5000, control = list(adapt_delta = 0*.*99, max_treedepth = 15)*. In each QN, we specify priors, model formulas, family distributions, and handling of research outputs. We share the fitted statistical models as .rds (R) objects in the “[Datasets/StatisticalModels_brms](https://osf.io/zvnxa/)” OSF repository component, and result tables as .tex or .html files in the “[Tables](https://osf.io/e7gwm/)” component. Model summaries are available in supplementary methods in [GitHub](https://github.com/elalilab/Stroke_PDGFR-B_Reactivity/blob/main/PDGFR-B_Reactivity_SupplementaryTables.qmd). We report the mean parameter or coefficient estimates with their 95% credible intervals (CI) (Hespanhol et al., 2019). Note that the scale of the coefficients may vary (link function ≠ identity) depending on the [family distribution](https://rdrr.io/cran/brms/man/brmsfamily.html) used to fit the model. For model diagnostics, we plotted the model predictions against the data using the *pp_check* function from brms, and performed model evaluation/selection based on the Watanabe-Akaike Information Criterion (WAIC) or Leave-one-out (Loo) cross-validation using the loo package (Vehtari et al., 2017). For visualization of Bayesian models, we use [ggplot](https://ggplot2.tidyverse.org/) (Wickham, 2016) and [ggdist](https://mjskay.github.io/ggdist/) (Kay, 2024) packages, or the *conditional_effects* function from brms. We share the source code in each QN and the individual plots in the “[Communication/Plots](https://osf.io/jxb4z/)” component of the OSF repository. Broadly, the results are presented using *stat_halfeye* from ggdist to depict probability densities or as point estimates from *conditional_effects* with 95% CI represented as whiskers. When pertinent, we calculated pairwise contrasts of interest using the [emmeans](https://github.com/rvlenth/emmeans) package (Searle et al., 1980). These comparisons are presented as half-eye probability densities and point interval ranges, together with a Region of Practical Equivalence (ROPE) depicting the posterior density that falls within the error variance (Makowski et al., 2019).

### 2.10. Point pattern analysis (PPA)

We employed the spatstat package (Baddeley & Turner, 2005) and its associated dependencies to analyze the spatial arrangement of cells through point pattern analysis (PPA). Using functions within spatstat, we created point patterns (*ppp*), calculated density kernels (*density*), estimated relative distributions (*rhohat)*, and fitted multiple point process models (*mppm*). The point patterns as “.rds” hyperframes files are shared in the “[Datasets/PointPatterns](https://osf.io/zvnxa/)” component of the OSF repository. We used the base plotting system in R to visualize the graphical results and tables from PPA. For density kernels, the *topo*.*colors(256)* color scale was applied tailored to each data set (refer to QNs and figure legends). We plotted the relative distributions as the pooled (per group) mean (displayed in magenta) with two-sigma confidence intervals represented by a gray-shaded region. Additionally, we depicted the pooled mean intensity as a *cyan-dashed line*, along with the corresponding lower (*black-*solid line) and upper (*green-dotted line*) limits of the two-sigma confidence intervals. We note that the scales of the plots were adjusted or constrained to facilitate visualization (see QNs and figure legends). Since the mppm spatial models utilize a Poisson distribution, the summary tables in the QNs are presented on the log scale.

### 2.11. Topological data analysis (TDA)

We used the Ghudi and Ripser Python libraries to extract topological features from cell-derived point clouds, which we share as .npy files in the “[Datasets/PointClouds](https://osf.io/zvnxa/)” component of the OSF repository. To create Vietoris-Rips complexes, we utilized gd.RipsComplex with specific filtration values **(see JNs; Suppl. Method 13)**. We share sets of [static](https://osf.io/f59bt) and [animated](https://osf.io/49aqu) Vietoris-Rips complexes in the “[Communication/Plots](https://osf.io/jxb4z/)” OSF component. The dimension-0 and dimension-1 persistent homology (Otter et al., 2017) were then calculated using *ripser*, for which the corresponding persistence diagrams are shared as .png [plots](https://osf.io/exf7s) or .npy [objects](https://osf.io/fyasg). Using these persistent diagrams, we computed Betti curves and compared the topological features of PDGFRβ^+^ cells among different time points within the specific ROIs. To obtain point estimates and 95% confidence intervals, we bootstrapped (1000 iterations) (Chernik, 2007b) the median bottleneck (*bottleneck_distance*) and Wasserstein (*wasserstein_distance*) distances. These results are presented as heat maps displaying median distances along with their corresponding 95% CI.

## 3. Results

### 3.1. Astrocytes and PDGFRβ^+^ cells respond distinctively to ischemic stroke injury

Ischemic stroke pathophysiology involves neuroinflammation associated with brain edema and subsequent atrophy of the injured tissue (Gu et al., 2022; Krueger et al., 2019). We imaged the complete ischemic hemisphere to analyze the ratio of brain shrinkage (atrophy) in mice subjected to experimental stroke **(Figure 2A; see Suppl. Method 1)**. Our analysis estimates that from 7 to ∼20 DPI, shrinkage proceeds at a rate of 0.34 mm^2^ per day [7 DPI = 14 mm^2^ ± 0.4; 30 DPI = 7.6 mm^2^ ± 0.4] ([QN](https://github.com/elalilab/Stroke_PDGFR-B_Reactivity/blob/main/Widefield_5x_Ipsilateral_Pdgfrb_Shrinkage.qmd); **Figure 2B; Suppl. Figure 1D-E; Suppl. Table 1A-B)**. This represents a ∼46% reduction in hemispheric volume, which accounts for a substantial loss of the cortical mass and the gradual impairment of the cytoarchitectural division between the cortex and striatum at the level of corpus callosum. Glial scar is formed essentially by reactive astrocytes expressing high levels of GFAP surrounding a fibrotic scar formed mainly by reactive PDGFRβ^+^ cells exhibiting fibroblast-like characteristics (Fernandez-Klett et al., 2013; Goritz et al., 2011; Zamanian et al., 2012; R. Zhang et al., 2018). Using PDGFRβ^tdTomato^ reporter mice, we assessed the reactivity of PDGFRβ^+^ cells that endogenously express tdTomato (PDGFRβ^tdTomato^ cells) in conjunction with GFAP^+^ astrocytes by quantifying the changes in integrated density (IntDen) **(see Suppl. Method 2)** over the course of injury progression **(Figure 2C; Suppl. Figure 1A)**. Analysis for PDGFRβ^+^ cells ([QN](https://github.com/elalilab/Stroke_PDGFR-B_Reactivity/blob/main/Widefield_5x_Ipsilateral_Pdgfrb_IntDen.qmd); **Suppl. Table 2A-B)** yields a non-linear increase in the IntDen until the 3^rd^ week after ischemic stroke [0 to 23 DPI = 841, β1 = 36], followed by a nearly-plateau phase [23 to 30 DPI = -29, β1 = -4.4] **(Figure 2D; Suppl. Figure 1D-E)**. We found that reactive PDGFRβ^+^ cells were essentially located outside the envelope formed by GFAP^+^ cells, a region prone to shrinkage and liquefaction. In PDGFRβ^tdTomato^ cells, tdTomato intensity is proportional to the basal activity of PDGFRβ promoter that drives initial Cre-recombinase expression. Notably, by conditioning IntDen to the hemispheric area **(Figure 2E; Suppl. Table 3)**, our model predicts that tdTomato IntDen increases in small hemispheric volumes, suggesting that the reactivity of PDGFRβ^+^ cells implicates PDGFRβ activation. An analogous analysis for GFAP^+^ cells ([QN](https://github.com/elalilab/Stroke_PDGFR-B_Reactivity/blob/main/Widefield_5x_Ipsilateral_Gfap_IntDen.qmd)) reveals a similar increase the IntDen until the 2^nd^ week after ischemic stroke [0 to 16 DPI = +835, β1 = 50.1], followed by a resolution phase [16 to 30 DPI = -480, β1 = -36] **(Suppl. Figure 1A-E: Suppl. Table 4A-B)**. Notably, when conditioning to the hemispheric area, our model predicts an increased reactivity of GFAP^+^ cells at 30 DPI under non shrinking conditions **(QN, Suppl. Figure 1C; Suppl. Table 5)**. This implies that brain shrinkage upon ischemic stroke favors emergence of reactive PDGFRβ^+^ cells and does not substantially affect the distribution of reactive GFAP^+^ cells.

**Figure 2.**
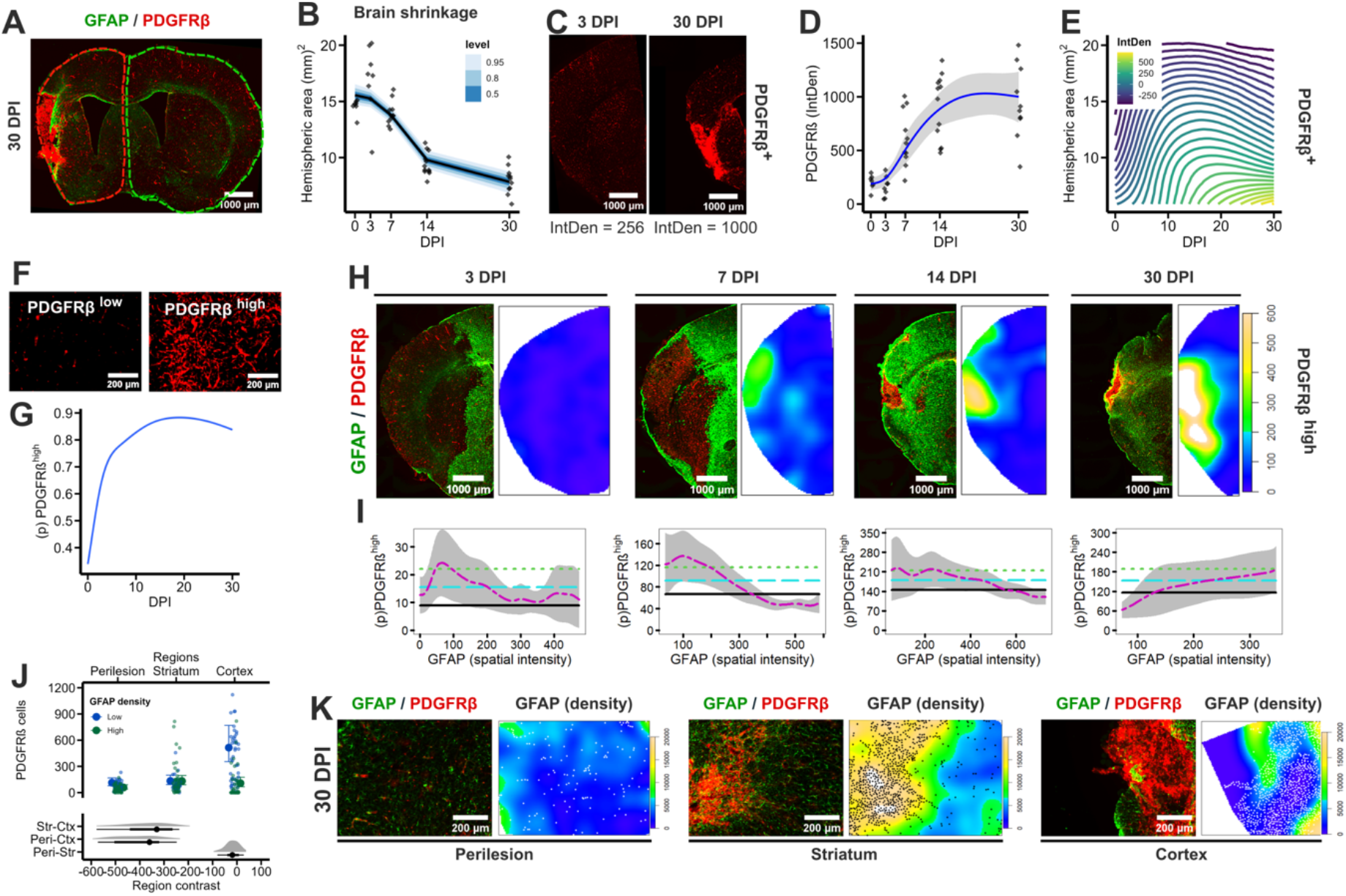
PDGFRβ^+^ cells actively respond to injury after ischemic stroke to partake in the formation of an inner scar compartment. **A)** Representative brain section (stitched, 5x magnification) at 30 DPI showing the measured ipsilateral (red outline) and contralateral (green outline) hemispheres. **B)** Fitted splines [model](https://osf.io/z6qus) for hemispheric area ∼ s(DPI). The stat_lineribbon() is displayed with the mean hemispheric area (black) and different credible intervals: 0.05, 0.8, 0.95 (blue brewer scale) and observations as diamonds (see Suppl. Figure 1 D-E (left); Suppl. Table 1; source [data](https://osf.io/3t8b2), [code](https://github.com/elalilab/Stroke_PDGFR-B_Reactivity/blob/main/Widefield_5x_Ipsilateral_Pdgfrb_Shrinkage.qmd), and [plot](https://osf.io/fr29g)). **C)** Representative brain sections (stitched, 5x magnification) at 3 and 30 DPI showing PDGFRβ^tdTomato^ cells (see [image set](https://osf.io/jrzq5)). Integrated density (IntDen) values at the button are shown as a reference. **D)** Fitted splines [model](https://osf.io/ag87j) for PDGFRβ_IntDen_ ∼ s(DPI). The conditional_effects are displayed with the mean (blue) and 95% CI (gray-shadowed region). The observations are displayed as black diamonds (see Suppl. Figure 1D-E (middle); Suppl. Table 2; source [data](https://osf.io/3t8b2), [code](https://github.com/elalilab/Stroke_PDGFR-B_Reactivity/blob/main/Widefield_5x_Ipsilateral_Pdgfrb_IntDen.qmd), and [plot](https://osf.io/fr29g)). **E)** Fitted splines [model](https://osf.io/kyqnc) for PDGFRβ_IntDen_ ∼ t2(DPI, Area). The conditional_effects are displayed with a Viridis scale (IntDen) (see Suppl. Table 3; source [data](https://osf.io/3t8b2), [code](https://github.com/elalilab/Stroke_PDGFR-B_Reactivity/blob/main/Widefield_5x_Ipsilateral_Pdgfrb_IntDen.qmd), and [plot](https://osf.io/56q9f)). **F)** Sample crops of PDGFRβ^low^ (non-reactive) and PDGFRβ^high^ (reactive) cells in the ischemic hemisphere. **G)** Fitted binomial [model](https://osf.io/nkgsm) for PDGFRβ^high^ | PDGFRβ^total^ ∼ s(DPI). The conditional_effects are displayed with mean (blue) and 95% CI (gray-shadowed region) (see Suppl. Table 6; source [data](https://osf.io/3t8b2), [code](https://github.com/elalilab/Stroke_PDGFR-B_Reactivity/blob/main/Widefield_5x_Ipsilateral_Pdgfrb_LowHigh.qmd), and [plot](https://osf.io/rq2ad)). **H)** Representative GFAP^+^/PDGFRβ^tdTomato^ immunolabeling in brain sections (left) (stitched, 5x magnification) with corresponding GFAP density kernels (right) at 3, 7, 14, and 30 DPI. The density kernels are represented using the *topo*.*colors(256)* scale (see [image set](https://osf.io/jrzq5); [point patterns](https://osf.io/ab4y5), and density kernels available in the [QN](https://github.com/elalilab/Stroke_PDGFR-B_Reactivity/blob/main/Widefield_5x_Ipsilateral_Gfap-Pdgfrb_Covariance.qmd)). **I)** Relative distribution diagrams (*rhohat*) depicting the probability of PDGFRβ^high^ cells allocation (y-axis) relative to GFAP spatial intensity (x-axis). The pooled mean intensity (*cyan-dashed line*) is shown with their corresponding lower (*black-*solid line) and upper (*green-dotted line*) limits of the two-sigma confidence intervals. Note the dissimilar *xy* axes limited by the mapping of the rhohat function for each DPI (see source [point patterns](https://osf.io/ab4y5), [code](https://github.com/elalilab/Stroke_PDGFR-B_Reactivity/blob/main/Widefield_5x_Ipsilateral_Pdgfrb_LowHigh.qmd)). The raw plots are available in the “[Communication/Plots](https://osf.io/jxb4z/)” component of the OSF repository. **J)** Fitted [model](https://osf.io/2kfye) for PDGFRβ^high^ ∼ Region:GFAP. The conditional_effects are displayed with mean cell counts and 95% CI as whiskers. At the bottom, entire posterior densities for the contrast between regions are shown with stat_pointinterval (see Suppl. Figure 2F; Suppl. Table 8; source [data](https://osf.io/cbgur), [code](https://github.com/elalilab/Stroke_PDGFR-B_Reactivity/blob/main/Widefield_10x_ROIs_Gfap-Pdgfrb_Covariance.qmd), and [plot](https://osf.io/rdbgt)). **K)** Representative GFAP^+^/PDGFRβ^tdTomato^ immunolabeling in brain sections (left) (10x magnification) with corresponding GFAP density kernels (right) at 30 DPI. The density kernels are represented using the *topo*.*colors(256)* scale and PDGFRβ^tdTomato^ cells as black or white dots (see [image set](https://osf.io/qrgws); [point patterns](https://osf.io/3fymw), and density kernels available in the [QN](https://github.com/elalilab/Stroke_PDGFR-B_Reactivity/blob/main/Widefield_10x_ROIs_Gfap-Pdgfrb_Covariance.qmd)). High-resolution figure: https://osf.io/jxb4z/files/m9g8v.

### 3.2. Highly reactive PDGFRβ^+^ cells aggregate in the inner compartment of the scar

The glial and fibrotic scars are composed of different cells that dynamically interact to fulfill various functions, mainly limiting the expansion of inflammatory signals towards intact tissue and preserving tissue integrity (Faulkner et al., 2004). Herein, we employed PPA to analyze the spatial localization of reactive PDGFRβ^+^ cells that form the inner scar compartment relative to reactive GFAP^+^ astrocytes that form the outer compartment ([QN](https://github.com/elalilab/Stroke_PDGFR-B_Reactivity/blob/main/Widefield_5x_Ipsilateral_Gfap-Pdgfrb_Covariance.qmd), **Suppl. Method 4)**. Ahead, we used machine learning-based classifiers in QuPath to identify PDGFRβ^high^ (high tdTomato intensity = highly reactive) and PDGFRβ^low^ (low tdTomato intensity = quiescent or non-reactive) cells ([QN](https://github.com/elalilab/Stroke_PDGFR-B_Reactivity/blob/main/Widefield_5x_Ipsilateral_Pdgfrb_LowHigh.qmd); **Suppl. Method 3)**. We observed an increasing likelihood of PDGFRβ^high^ cells [β1 = 25 ± 0.85] in lesioned areas up to the 2^nd^ week after ischemic stroke [∼80 - 90 %] **(Figure 2F-G; Suppl. Table 6)**. This highlights that most of the PDGFRβ^+^ cells actively respond to brain injury. We then examined the distribution of PDGFRβ^high^ cells relative to reactive GFAP^+^ cells using PPA ([QN](https://github.com/elalilab/Stroke_PDGFR-B_Reactivity/blob/main/Widefield_5x_Ipsilateral_Gfap-Pdgfrb_Covariance.qmd); **Suppl. Method 4)**. Our graphical results (*rhohat*) and *mppm* show that PDGFRβ^high^ cells cluster outside the envelope formed by GFAP^+^ cells (convex hull), particularly during the 1^st^ 2 weeks after ischemic stroke [β_1_ = -0.0015 (3D); -0.0018 (7D); -0.00067 (14D)]. At 30 DPI, we observe a moderate overlap between both cell populations [β_1_ = 0.0017] **(see Figure 2H-I;** [QN](https://github.com/elalilab/Stroke_PDGFR-B_Reactivity/blob/main/Widefield_5x_Ipsilateral_Gfap-Pdgfrb_Covariance.qmd)). We interpret this observation as the result of brain atrophy and subsequent loss of the defined boundaries between the cortex and the striatum in the injured tissue. In contrast, PDGFRβ^low^ cells, which are essentially located at the perivascular space, remain predominantly localized with the structure comprising GFAP^+^ cells, as shown in **Suppl. Figure 1F** and [QN](https://github.com/elalilab/Stroke_PDGFR-B_Reactivity/blob/main/Widefield_5x_Ipsilateral_Gfap-Pdgfrb_Covariance.qmd). Collectively, this suggests that highly reactive PDGFRβ^+^ cells form an inner and well-delineated compartment at the boundaries of GFAP^+^ glial scar. To further validate this preceding notion, we imaged whole brain sections to assess the establishment of GFAP convex hull after brain injury ([QN](https://github.com/elalilab/Stroke_PDGFR-B_Reactivity/blob/main/Widefield_5x_Ipsilateral_Gfap_Convex.qmd); **Suppl. Method 5; Suppl. Figure 1G)**. The results denote that GFAP^+^ glial scar borders are formed within the 1^st^ week after ischemia and remain stable over time [β1 = -0.10, 95% CI = -0.38 – 0.18] **(Suppl. Figure 1G-H; Suppl. Table 7)**. Based on these results, we hypothesize that reactive PDGFRβ^+^ cells are early destined to localize to the injured regions prone to irreversible degeneration and possible liquefaction (see next section), as delimited by the envelope formed by GFAP^+^ cells during the 1^st^ week of ischemic stroke. This concept is further supported by imaging of ROIs at the injured cortex, the injured striatum, and the healthy peri-lesion region, and analyses using cell counts in GFAP^low^ and GFAP^high^ density-defined tessellations ([QN](https://github.com/elalilab/Stroke_PDGFR-B_Reactivity/blob/main/Widefield_10x_ROIs_Gfap-Pdgfrb_Covariance.qmd), **Suppl. Method 6)**. Indeed, we observe a ∼ 3-fold change [β_cortex_ = exp(1.06) = 2.89, 95% CI = 1.9 – 4.3) in the value for PDGFRβ^high^ cells in GFAP^low^ areas in the injured cortex compared to the healthy peri-lesional region. For GFAP^high^ areas in the same regions, this value decreases by a factor of exp(−0.64) = -1.89, 95% CI -3.32 – -1.06 **(Figure 2J-K)**. The [QN](https://github.com/elalilab/Stroke_PDGFR-B_Reactivity/blob/main/Widefield_10x_ROIs_Gfap-Pdgfrb_Covariance.qmd), **Suppl. Table 8** and **Suppl. Figure 2E-F** portrait estimates by region/DPI.

Severity of brain injury after ischemic stroke has been proposed to affect the response of PDGFRβ^+^ cells (Dias et al., 2021). We next examined the brain of animals in which mild injury was induced only in the striatum at 14 and 30 DPI ([QN](https://github.com/elalilab/Stroke_PDGFR-B_Reactivity/blob/main/Widefield_5x_Ipsilateral_Gfap-Pdgfrb_Covariance_Str.qmd), **Suppl. Method 6)** to evaluate the reactivity of PDGFRβ^+^ cells in conditions that do not involve massive brain atrophy and substantial cortical damage. Our results show that the probability of the emergence of PDGFRβ^high^ cells is approximately half of that of larger cortico-striatal injury (∼0.90 vs 0.45) **(Suppl. Figure 2A-B; Suppl. Table 9)**. Therefore, we conclude that PDGFRβ^high^ and PDGFRβ^low^ cells are not segregated in mild striatal-specific injury as it is the case in more severe cortico-striatal injury, involving important brain atrophy **(Suppl. Figure 2C-D)**. This is consistent with the notion that aggregation of reactive PDGFRβ^high^ cells is characteristic of irreversible tissue damage upon brain injuries. Nonetheless, reactivity of PDGFRβ^+^ cells specific to the different brain structures cannot be excluded.

### 3.3. Phenotypically distinct populations of PDGFRβ^+^ cells are present at the injured tissue

Ischemic stroke induces a rapid loss of CD13^+^ pericytes followed by emergence of reactive PDGFRβ^+^ cells that are proposed to ‘detach’ from the vasculature to migrate towards the injured parenchyma **(Figure 3A)** (Dore-Duffy et al., 2000; Fernandez-Klett et al., 2013). It is noteworthy to mention that these investigations were performed using antibody-based approaches to track brain PDGFRβ^+^ cells. Herein, using PDGFRβ^tdTomato^ reporter mice, we analyzed the dynamics of perivascular and parenchymal PDGFRβ^+^ cell populations and crosstalk with CD31^+^ vasculature in specific ROIs in the injured cortex, the injured striatum and the healthy peri-lesional region ([QN](https://github.com/elalilab/Stroke_PDGFR-B_Reactivity/blob/main/Widefield_10x_ROIs_CD31-Pdgfrb_Coloc.qmd), **Suppl. Method 7)**. Our analysis shows that in the healthy peri-lesional region, the probability of parenchymal PDGFRβ^+^ cells (non-vascular associated) ranges only from 3-9%. This value peaks at ∼25%, in the injured striatum and to ∼50% in the injured cortex during the 2^nd^ week after ischemic stroke **(Figure 3B-C; Suppl. Table 10C)**. We display cell counts and the contrast between the injured cortex and injured striatum in **(Suppl. Figure 3A-B)**. We interpret the decreased probability of parenchymal PDGFRβ^+^ cells in the injured regions as the result of progressive cell loss over time, presumably due to degeneration and tissue liquefaction.

**Figure 3.**
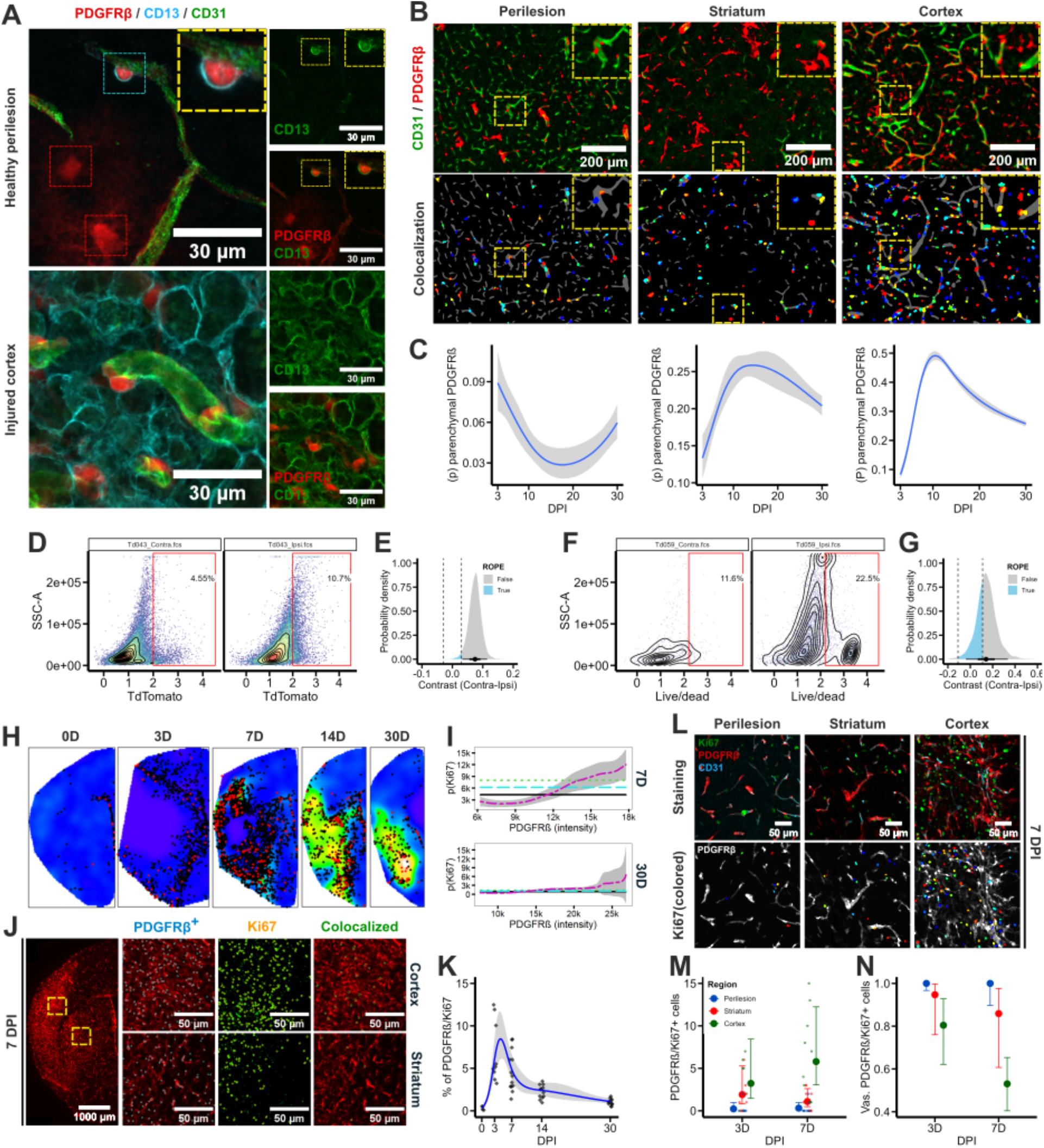
Reactive PDGFRβ^+^ cells comprise perivascular and parenchymal cells. **A)** Representative images (40x magnification) of PDGFRβ^+^ cells in the healthy peri-lesional and injured cortex. PDGFRβ^tdTomato^ cells are associated with CD31^+^ vasculature, and their nuclei co-localize with CD13, a common pericyte marker. Parenchymal PDGFRβ^tdTomato^/CD13^-^ cells are visible interacting with the CD31^+^ vasculature through fine ramifications. In the injured cortex, nuclei-based colocalization with CD13 is lost and parenchymal PDGFRβ^tdTomato^ cells are not in place (see the entire [image set](https://osf.io/mfyge)). **B)** Representative images (10x magnification) of PDGFRβ^tdTomato^/CD31^+^ immunolabeling (upper row), and co-localization analysis (CellProfiler) of PDGFRβ^tdTomato^ nuclei (colored) and CD31^+^ vasculature (gray) (bottom row) for peri-lesional region, striatum, and cortex. The lined-yellow box is zoomed in the upper right corner of each image (see the entire [image set](https://osf.io/csvrx)). **C)** Fitted binomial models per ROIs for PDGFRβ^colocalized^ | PDGFRβ^total^ ∼ s(DPI). The conditional_effects are displayed with a mean (blue) and 95% CI (gray-shadowed region) (see Suppl. Figure 3A-B; Suppl. Table 10; source [data](https://osf.io/zfcgh), [code](https://github.com/elalilab/Stroke_PDGFR-B_Reactivity/blob/main/Widefield_10x_ROIs_CD31-Pdgfrb_Coloc.qmd), and plots in the “Communication/Plots” component of the OSF repository. **D, F)** Representative images for contralateral (left, contra) and ipsilateral (right, ipsi) hemispheres showing total PDGFRβ^tdTomato^ cells (D) and PDGFRβ^tdTomato^/Live Dead^+^ cells (F). In both cases, ggcyto is displayed with geom_density2d, geom_gate, and geom_stats and additional ggplot aesthetics (see Suppl. Figure 4; source data in the “[Datasets/FACS](https://osf.io/zvnxa/)” component of the OSF, [code](https://github.com/elalilab/Stroke_PDGFR-B_Reactivity/blob/main/FlowCytometry_Pdgfrb_Processing.qmd), and plots in the “[Communication/Plots/FACS_Pdgfrb](https://osf.io/jxb4z/)” component of the OSF repository and). **E, G)** Contrast between the percentage of PDGFRβ^tdTomato^ cells (E) and the percentage of PDGFRβ^tdTomato^ dead cells (G) in the contralateral (contra) and ipsilateral (ipsi) hemispheres for the linear model Cells ∼ Hemisphere. Posterior draws (spread_draws) are shown with stat_halfeye() and ROPE (see Suppl. Table 11; source [data](https://osf.io/7h6mg), [code](https://github.com/elalilab/Stroke_PDGFR-B_Reactivity/blob/main/FlowCytometry_Pdgfrb_Analysis.qmd), and plots for [E](https://osf.io/ac2g4) and [G](https://osf.io/49k2a). **H)** Representative density kernels of PDGFRβ^tdTomato+^ cells at 0 (control), 3, 7, 14 and 30 DPI. The density kernels are represented using the *topo*.*colors(256)* scale, and KI67^+^ cells are represented as black dots and co-localized PDGFRβ^tdTomato^/Ki67^+^ immunolabeling is red colored (see [point patterns](https://osf.io/fxgr6) and density kernels in the [QN](https://github.com/elalilab/Stroke_PDGFR-B_Reactivity/blob/main/Widefield_10x_Ipsilateral_Ki67_PPA.qmd)). **I)** Relative distribution diagrams (*rhohat*) depicting the probability of Ki67^+^ cell allocation (y-axis) relative to PDGFRβ^tdTomato^ cell spatial intensity (x-axis). The pooled mean intensity (*cyan-dashed line*) is shown with their corresponding lower (*black-*solid line) and upper (*green-dotted line*) limits of the two-sigma confidence intervals. Note the dissimilar *xy* axis limits defined by the point arguments mapped by the rhohat function for each DPI (see source [point patterns](https://osf.io/fxgr6), [code](https://github.com/elalilab/Stroke_PDGFR-B_Reactivity/blob/main/Widefield_10x_Ipsilateral_Ki67_PPA.qmd), and plots for 7 and 30 DPI). **J)** (left) Representative image (stitched 10x magnification) of PDGFRβ^tdTomato^/Ki67^+^ staining in brain section at 7 DPI. The upper (cortex) and bottom (striatum) panels show the boxed yellow regions, with overlays of detected (CellProfiler) PDGFRβ^tdTomato^ cells (cyan), Ki67 (orange), and co-localized PDGFRβ^tdTomato^/Ki67^+^ cells (green) (see the entire [image set](https://osf.io/52h64)). **K)** Fitted splines [model](https://osf.io/cnrbg) for Percentage ∼ s(DPI, k = 5), family = lognormal. The conditional_effects are displayed with mean (blue) and 95% CI (gray-shadowed region) and observations as diamonds (see Suppl. Table 15 with complementary model derivatives; source [data](https://osf.io/km6hv), [code](https://github.com/elalilab/Stroke_PDGFR-B_Reactivity/blob/main/Widefield_10x_Ipsilateral_Ki67-Pdgfrb_Coloc.qmd), and [plot](https://osf.io/sndb3)). **L)** Representative images (20x magnification) of PDGFRβ^tdTomato^/CD31^+^/Ki67^+^ immunolabeling in brain sections (upper row) in the peri-lesional region, the injured striatum and the injured cortex at 7 DPI. The bottom row depicts Ki67^+^ cells (colored) and PDGFRβ^tdTomato^ cells (gray) (see the entire [image set](https://osf.io/ut7h5)). **M)** Fitted [model](https://osf.io/hnuca) for the total number of colocalized PDGFRβ^tdTomato^/Ki67^+^ cells (co-localized ∼ DPI * Region, family = negbinomial). Posterior estimates are displayed with conditional_effects (mean + 95% CI). Dots are observations (see Suppl. Table 16; source [data](https://osf.io/6a95w), [code](https://github.com/elalilab/Stroke_PDGFR-B_Reactivity/blob/main/Confocal_20x_ROIs_Ki67-Pdgfrb_Coloc.qmd), and [plot](https://osf.io/sndb3)). **N)** Fitted [model](https://osf.io/zd5x3) for the proportion of vascular PDGFRβ^tdTomato^/Ki67^+^ cells (Vascular | trials(total) ∼ DPI * Region, family = binomial). Posterior estimates are displayed with conditional_effects (mean + 95% CI) (see Suppl. Table 16; source [data](https://osf.io/6a95w), [code](https://github.com/elalilab/Stroke_PDGFR-B_Reactivity/blob/main/Confocal_20x_ROIs_Ki67-Pdgfrb_Coloc.qmd), and [plot](https://osf.io/mnjhz)). High-resolution figure: https://osf.io/jxb4z/files/s65pg.

To substantiate this perspective, we sorted PDGFRβ^+^ cells using FACS to assess the extent of cell death **(Figure 3D-G; Suppl. Figure 4; Suppl. Method 8)**. We found that the percentage of total PDGFRβ^+^ cells [β_1_ = 0.074, 95% CI = 0.033 – 0.114] as well as dead PDGFRβ^+^ cells [β_1_ = 0.137, 95% CI = -0.006 – 0.299] are likely to increase at 14 DPI **(see Suppl. Table 11A-B)**. We note that although this indicates that reactive PDGFRβ^+^ cells populating the injured tissue are prone to death, further research using specific methods and causal inference is needed to further elucidate this specific aspect. Furthermore, we analyzed the proportion of parenchymal PDGFRβ^+^ cells upon striatal-specific injury at 14 and 30 DPI **(see Suppl. Method 7)**. Interestingly, we report an increasing trend in the probability of parenchymal PDGFRβ^+^ cells **(Suppl. Figure 3E; Suppl. Table 12)**, even though the total cell count rises in both, striatal-specific and cortico-striatal lesions at 14 and 30 DPI **(Suppl. Figure 3C-D; Suppl. Table 13)**. We hypothesize that these divergent trends between mild striatal and cortico-striatal injuries could stem from accelerated cell loss associated with injury severity in the latter. Finally, we ruled out the possibility that prolonged Cre-recombination via 4-OHT **(see Suppl. Method 7)** induces stronger tdTomato expression in parenchymal PDGFRβ^low^ cells, which could account for the increased density of overall PDGFRβ^+^ parenchymal cells **(Suppl. Figure 3F-G; Suppl. Table 14)**.

### 3.4. Reactive PDGFRβ^+^ cells do not massively proliferate within the injured brain

Previous studies using antibodies targeting PDGFRβ have reported important proliferation of PDGFRβ^+^ cells at the injured site during the 1^st^ week of ischemic stroke as well as after traumatic brain injury (Özen et al., 2014; Zehendner et al., 2015). Based on these observations, we hypothesized that the appearance of PDGFRβ^+^ parenchymal cells may be due to localized proliferation of resident cells rather than cell detachment from the vasculature, as previously suggested (Dore-Duffy et al., 2000; Fernandez-Klett et al., 2013; Menezes et al., 2020). Accordingly, we immunolabelled the proliferation marker Ki67 to analyze its distribution relative to the spatial intensity of PDGFRβ^tdTomato^ cells as well as its co-localization with individual PDGFRβ^tdTomato^ cells **(see Suppl. Method 9-10)**. First, we observed consistent expression of Ki67 in regions of elevated tdTomato spatial intensity [β_0_ = 6.85, β_1_ =7.26^e-0^], particularly at 3 [β_1_ = 1.07^e-04^] and 7 DPI [β1 = 7.05^e-06^] ([QN](https://github.com/elalilab/Stroke_PDGFR-B_Reactivity/blob/main/Widefield_10x_Ipsilateral_Ki67_PPA.qmd), **Figure 3H-I)**. These findings are consistent with the critical periods of cell proliferation associated with glial scar formation (Wanner et al., 2013; Xie et al., 2020). However, in contrast to what is reported in previous studies, our analysis revealed that the proportion of PDGFRβ^+^/Ki67^+^ cells (relative to the total number of PDGFRβ^+^ cells) does not exceed 10% at its peak at 3 DPI [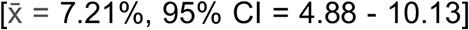, 95% CI = 4.88 - 10.13] **(see** [QN](https://github.com/elalilab/Stroke_PDGFR-B_Reactivity/blob/main/Widefield_10x_Ipsilateral_Ki67-Pdgfrb_Coloc.qmd), **Figure 3J-K; Suppl. Table 15)**. We acknowledge that these results occurred despite potential systemic overestimation of co-localized PDGFRβ^+^/Ki67^+^ cells due to partial object overlap inherent in our automated/unbiased analysis method **(see Suppl. Method 9)**. We further analyzed PDGFRβ^+^/Ki67^+^ cells by laser scan confocal microscopy (20x magnification) along with the endothelial marker CD31 in defined ROIs **(Figure 3L;** [QN](https://github.com/elalilab/Stroke_PDGFR-B_Reactivity/blob/main/Confocal_20x_ROIs_Ki67-Pdgfrb_Coloc.qmd)**; Suppl. Method 10)**. Our results reveal that only a small fraction of co-localizing PDGFRβ^+^/Ki67^+^ cells **(Figure 3M; Suppl. Table 16A)** is mainly derived from non-vascular associated cells in the injured cortex and even to a lesser extent in the injured striatum **(Figure 3N; Suppl. Table 16B)**. Overall, our findings with the PDGFRβ^tdTomato^ reporter mouse model suggest that reactive PDGFRβ^+^ cells do not seem to massively locally proliferate after injury, nor could the small fraction of proliferative cells account for the substantial increase in the density of PDGFRβ^+^ cells observed in the injured regions. In our view, the origin of the parenchymal proliferative PDGFRβ^+^ cells reported in the previous studies using antibody-based approaches remains ambiguous. On the one hand, these cells could be cells detached from the brain vasculature, as previously shown, or due to the recruitment at the injury core of an undefined subpopulation of infiltrating PDGFRβ^+^ peripheral cells with macrophage properties as previously reported by Sakuma et al. (2016). This implies new avenues in this research field.

### 3.5. Reactive brain PDGFRβ^+^ cells comprise morphologically diverse populations

Using a semiautomated computational pipeline in Python (skimage and scipy), we analyzed the morphological characteristics and changes of reactive parenchymal and vascular-associated PDGFRβ^+^ cells across injury progression after ischemic stroke ([QN](https://github.com/elalilab/Stroke_PDGFR-B_Reactivity/blob/main/Confocal_40x_ROIs_Pdgfrb_Morpho_BatchScript.qmd)**-batch processing**, [QN](https://github.com/elalilab/Stroke_PDGFR-B_Reactivity/blob/main/Confocal_40x_ROIs_Pdgfrb_Morpho_Analysis.qmd)**-analysis, Suppl. Method 11; Suppl. Figure 5)**. We observed that the healthy peri-lesional regions are associated with perivascular PDGFRβ^+^ cells that are abundantly co-labeled with CD13, a marker that is expressed in pericytes under normal conditions and some peripheral myeloid immune cells (Nguyen et al., 2023a) **(Figure 3A)**. In addition, our PDGFRβ^tdTomato^ reporter mouse line revealed the presence of a subpopulation of quiescent PDGFRβ^low^ cells in the same region. These cells, hereafter referred to as reticuloparenchymal cells, exhibit a highly ramified morphology and maintain substantial narrow contacts with the vasculature via fine processes. Unlike perivascular cells, these PDGFRβ^low^ cells do not express CD13, indicating that they are neither pericytes nor infiltrating immune cells. On the other hand, the injured brain contains PDGFRβ^high^ amoeboid cells, which may be either parenchymal or perivascular, as well as small ramified parenchymal cells, hereafter referred to as reticulite cells **(Figure 4A)**. These reticulite cells are immunoreactive for CD13, suggesting a pericytic or myeloid immune identity. However, we could not confirm this aspect in amoeboid cells due to the strong immunoreactivity, cell density, and background noise at the injury core **(Suppl. Figure 5C)**. Based on PCA ([QN](https://github.com/elalilab/Stroke_PDGFR-B_Reactivity/blob/main/Confocal_40x_ROIs_Pdgfrb_Morpho_Analysis.qmd)**-analysis, Figure 4B; Suppl Methods 11; Suppl. Figure 5B)**, we portray PDGFRβ^+^ cells along with three distinct morphological features **(Figure 4C; Suppl. Figure 5D)**. Note that because of the co-linearity between the area and convex hull, the (data variable) area in **Figure 4C** is shown only as a reference; (1) Perivascular PDGFRβ^+^ cells are anchored to CD31^+^ vasculature and exhibit a consistent branch length (0.73), convex hull (0.39), and intensity (0.31) **(Figure 4C, red)**. In contrast, (2) reticuloparenchymal cells are distinguished by a larger convex hull (0.82) while their nuclei are located within the brain parenchyma and interact with CD31^+^ vasculature through fine ramifications **(Figure 4C, green)**. (3) Reticulites, located primarily in the ischemic striatum, are characterized by moderate intensity (0.006) relative to their branch length (−0.58), and convex hull (−0.66) (**Figure 4C, cyan)**. (4) Amoeboid PDGFRβ^+^ cells, prevalent in the ischemic cortex, show high PDGFRβ activity (tdTomato relative intensity) (1.88) and are found both anchored to CD31^+^ vasculature as well as within the liquefied parenchyma **(Figure 4C, magenta)**. We argue that this latter feature may indicate high metabolic stress and susceptibility to cell death. **Suppl. Figure 5D** portrays the probability of each cell subtype given the referred morphological traits. Collectively, our results underscore a strong phenotypic diversity of PDGFRβ^+^ cells that is amplified in the injured brain after ischemic stroke. In our Zenodo and OSF repositories, we provide the raw images, segmented cells, and raw morphological measurements that can be re-used in further research aiming to gain more insights into the morphological changes of reactive PDGFRβ^+^ cells in various conditions.

**Figure 4.**
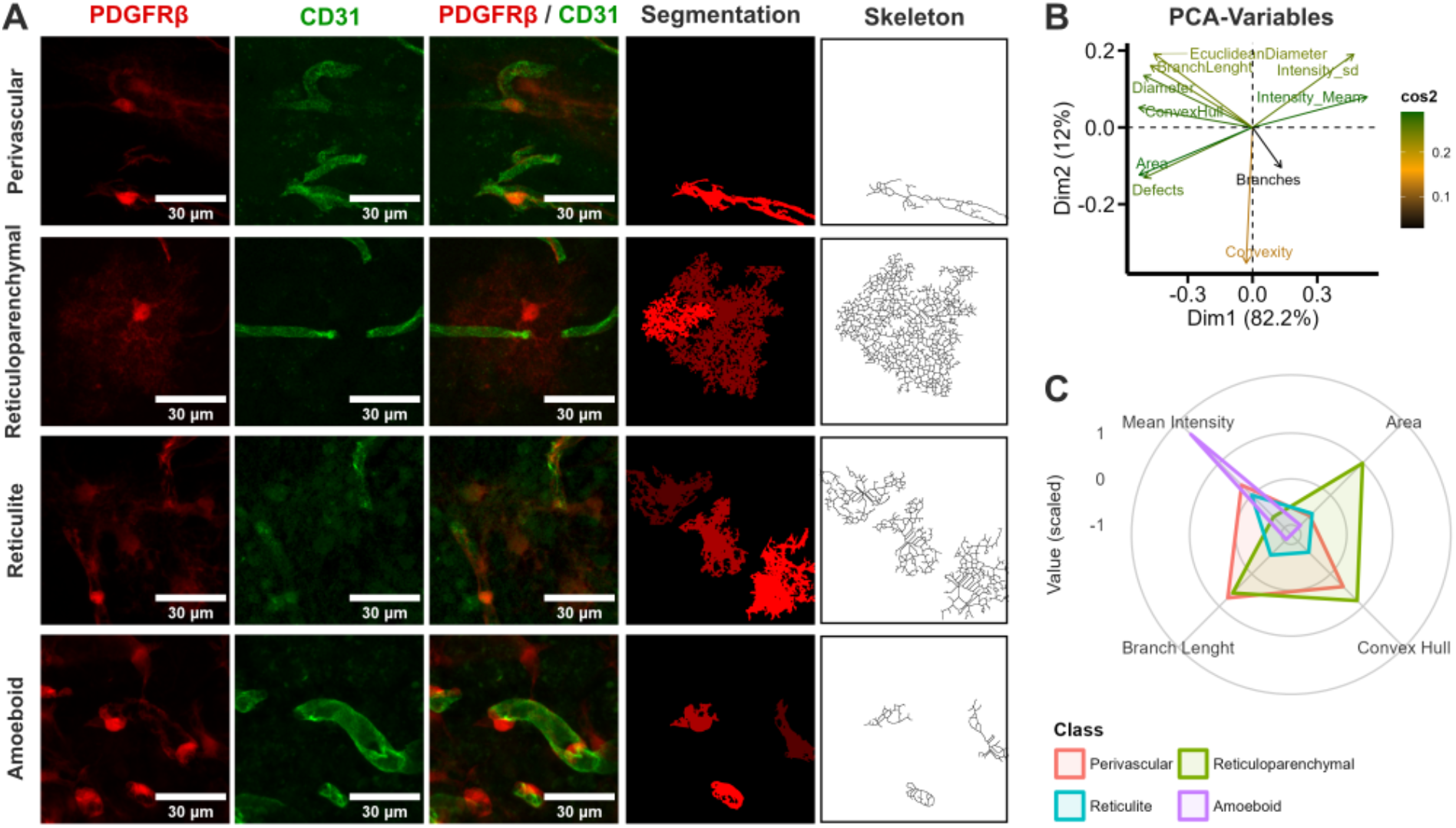
Morphological diversity of brain PDGFRβ^+^ cells. **A)** Representative images (40x magnification) showing the morphological diversity of PDGFRβ^+^ cells. The healthy brain hosts perivascular PDGFRβ^high^ and PDGFRβ^low^ cells, whereas the injured areas showcase predominantly reticulite and amoeboid PDGFRβ^high^ cells (see Suppl. Figure 5A; entire [image set](https://osf.io/mfyge), [cropped cells](https://osf.io/szpj6), and [segmentations](https://osf.io/h2uxq)). **B)** Biplot from the PCA analysis of morphological features of PDGFRβ^+^ cells. Variables negatively correlated are displayed on opposite sides of the Cartesian plane and its weight is shown as distance from the point of origin. Cos2 bar depicts the representation of each variable, with green as higher values (see Suppl. Figure 5B; source [data](https://osf.io/z3hyx), [code](https://github.com/daniel-manrique/Stroke_PDGFR-B_Reactivity/blob/main/Confocal_40x_Pdgfrb_Morphology.qmd), and [plot](https://osf.io/y3nfu)). **C)** Spyder plot showing the three selected variables (mean intensity, convex hull, branch length; areas as reference) via PCA to characterize the morphology of PDGFRβ^+^ cells (see Suppl. Figure 5C; source [data](https://osf.io/z3hyx), [code](https://github.com/daniel-manrique/Stroke_PDGFR-B_Reactivity/blob/main/Confocal_40x_Pdgfrb_Morphology.qmd), and [plot](https://osf.io/n5ysk)). High-resolution figure: https://osf.io/jxb4z/files/uv5cw.

During our investigations, we attempted to further characterize the morphologically distinct PDGFRβ^+^ cells, by co-immunolabeling for CD13 **(Suppl. Figure 6A)**, GFAP **(Suppl. Figure 6B)**, IBA1 **(Suppl. Figure 6C)**, and PDGFRα **(Suppl. Figure 6C)**. However, the high cell density and the lack of well nuclei-defined staining made the automated and unbiased co-localization analysis impracticable. Nonetheless, we observed that perivascular PDGFRβ^+^ cells are immunoreactive to CD13 (R&D systems, cat # AF2335, RRID:AB_2227288 antibody), whereas reticuloparenchymal PDGFRβ^+^ cells in the healthy brain tissue are not **(Suppl. Figure 6A)**. In the injured regions, some GFAP-stained cytoskeletons (Invitrogen, cat # 13-0300, RRID:AB_86543 antibody) appeared to encircle the nuclei of PDGFRβ^+^ cells **(Suppl. Figure 6B)**. However, under these conditions, asserting co-localization is ambiguous because GFAP^+^ cell branches cannot be unbiasedly associated with specific nuclei. Similarly, we could not confirm co-localization between PDGFRβ^+^ and IBA1^+^ cells (Wako, cat # 019-19741, RRID:AB_839504 antibody) as described by (Özen et al., 2014) due to increased cell density and staining artifacts introduced by the cell debris in the injured tissue. Moreover, PDGFRα staining (Abcam, cat # AB203491, RRID:AB_2892065) revealed that in the healthy brain, PDGFRα^+^ cells and PDGFRβ^+^ cells constitute two distinct populations. PDGFRα^+^ cells appeared as ramified parenchymal cells, whereas PDGFRβ^+^ cells are typical perivascular. This observation contrasts with previous reports using an antibody-based approach to immunolabel PDGFRβ with the Abcam antibody ab203491 (RRID:AB_2892065) (Menezes et al., 2020), which has claimed co-localization between these two markers in perivascular PDGFRβ^+^ cells. Notably, Abcam recently reclassified this antibody as recognizing both PDGFRα as well as PDGFRβ. Overall, we argue that further unbiased and reproducible investigations using simultaneous lineage tracking are needed.

### 3.6. Spatial and topological organization of reactive PDGFRβ^+^ cells upon injury

Next, we investigated the topological changes of reactive PDGFRβ^+^ cells in defined ROIs (peri-lesion, striatum, and cortex) after injury using Haralick features **(Suppl. Method 12)** and TDA **(Suppl. Method 13)**. The Haralick features ([QN](https://github.com/elalilab/Stroke_PDGFR-B_Reactivity/blob/main/Widefield_20x_ROIs_Pdgfrb_Haralick_BatchScript.qmd)**-batch processing;** [QN](https://github.com/elalilab/Stroke_PDGFR-B_Reactivity/blob/main/Widefield_20x_ROIs_Pdgfrb_Haralick.qmd)**-analysis)** were applied to understand image texture changes, as described by (Löfstedt et al., 2019). Our results by PCA ([QN](https://github.com/elalilab/Stroke_PDGFR-B_Reactivity/blob/main/Widefield_20x_ROIs_Pdgfrb_Haralick.qmd)**-analysis; Suppl. Figure 7A-B)** revealed that no Haralick features have a strong weight for PDGFRβ^+^ cell reactivity. Given these results, we focused on evaluating the variations in contrast, entropy, and the inverse difference moment (IDM) ([QN](https://github.com/elalilab/Stroke_PDGFR-B_Reactivity/blob/main/Widefield_20x_ROIs_Pdgfrb_Haralick.qmd)**-analysis and Suppl. Method 12)**. We found that increased IDM (homogeneity) is predominant in the different injured regions, especially the cortex [β_Ctx_ = 2.92, 95% CI = 1.13 – 4.83] **(Figure 5A; Suppl. Table 17)**, consistent with increased PDGFRβ^+^ cell-occupied area [β_Ctx_ = 4.5, 95% CI = 2.4 – 6.4] **(Suppl. Figure 7C; Suppl. Table 18)**. In contrast, the peri-lesional region showed high local variation and more complex textures (entropy) **(Figure 5A; Suppl. Table 17)**. In **Suppl. Figure 7D** and **Suppl. Table 17**, we show the influence of entropy, contrast, and IDM on the classification of brain regions. Entropy appears to be the most discriminative attribute, forming a gradient from the peri-lesional region (low entropy) towards the cortex (high entropy). Subsequently, we used TDA and persistence homology (Bhaskar et al., 2021b) to elucidate the reorganization patterns of PDGFRβ^+^ cells in the same brain regions ([JN](https://github.com/elalilab/Stroke_PDGFR-B_Reactivity/blob/main/Widefield_20x_ROIs_Pdgfrb_TDA.ipynb)**; Suppl. Method 13)**. In **Figure 5B**, we illustrate representative Vietoris-Rips complexes showing cell connectivity at a specific filtration value (200) (see OSF repository for the entire [image set](https://osf.io/f59bt) and [animations](https://osf.io/49aqu) for different filtration values, also shown in [Suppl. Video 1](https://osf.io/zteg3). We computed persistent homology **(Suppl. Figure 7E; see the full** [image set](https://osf.io/fyasg)) and derived 0 and 1-dimensional Betti curves to quantify the topological complexity of PDGFRβ^+^ cells over time **(Figure 5C)**. We then calculated Wasserstein **(Figure 6D)** and bottleneck **(Suppl. Figure 7F)** distances to represent the changes between DPI **(Suppl. Table 19)**. Our analysis of the number of connected components (0-dimensional homology) revealed a consistent merging pattern in the peri-lesional region at most time points **(Figure 5C, upper row)**. At 30 DPI (left-yellow line), however, we observed a higher merging at lower filtration values (∼120 filtration value). This suggests that the healthy peri-lesional region experiences small-scale clustering at this specific timepoint. It is consistent with the features of 2-dimensional loops (1-dimensional homology), where components tended to form discrete loops rather than a continuous monolayer of cells **(Figure 5C, left-yellow line)**. This suggests a lack of homogeneity (see Haralick features) of PDGFRβ^+^ cells in the healthy brain tissue.

**Figure 5.**
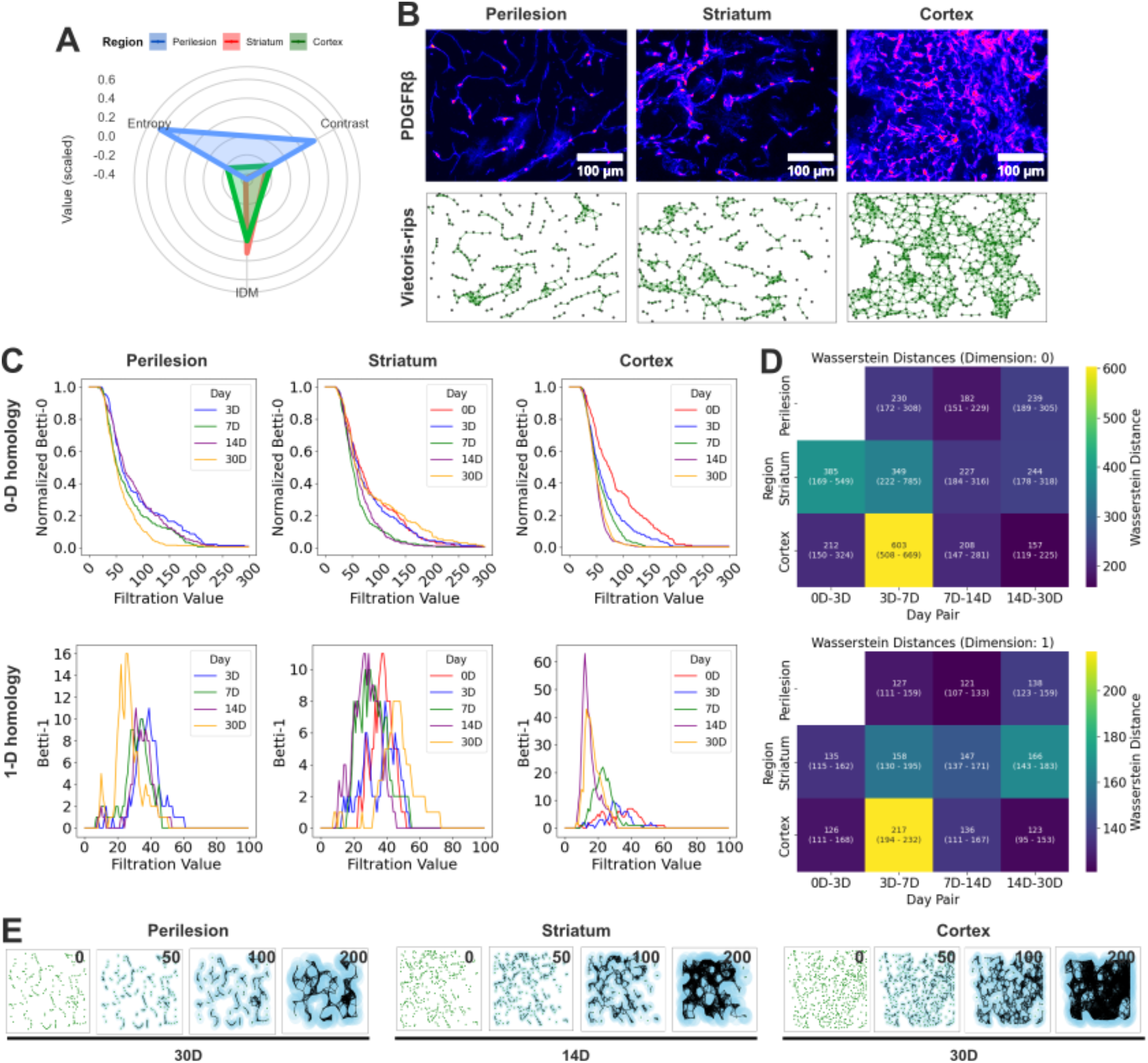
Haralick features and topological data analysis (TDA) of PDGFRβ^+^ cells. **A)** Spyder plot showing the three selected variables (Entropy, contrast, IDM) by PCA to characterize the Haralick variables of PDGFRβ^+^ cells. The represented regions (peri-lesional region, striatum, cortex) comprise animals from all the time points (see Suppl. Figure 7A-C; Suppl. Table 17; source [data](https://osf.io/hzqrm), [code](https://github.com/elalilab/Stroke_PDGFR-B_Reactivity/blob/main/Widefield_20x_ROIs_Pdgfrb_Haralick.qmd), and [plot](https://osf.io/t8wzk)). **B)** (upper row) Representative images (‘fire’ LUT) of the healthy peri-lesional region, the injured striatum and the injured cortex. Magenta represents higher PDGFRβ (tdTomato) promoter activity (see entire [image set](https://osf.io/m76pn)). (Low row) corresponding vietoris-rips complexes, showing the topological complexity of cells (see source [data](https://osf.io/32a8e), entire [image set](https://osf.io/f59bt), different [filtration values](https://osf.io/zxbas), and [animated](https://osf.io/49aqu) vietoris-rips complexes). **C-D)** Mean Betti curves (C) per time point and corresponding Wasserstein distances (D) between pairs of time points calculated from 0 and 1-dimensional homology persistent diagrams (Suppl. Figure 7E). The uncertainty in Wassertains distances was calculated using bootstrapping with 1000 iterations (see Suppl. Table 19C-D). Betti curves for 0-dimensional homology are normalized to 1 (1-dimensional graphs are not normalized). Line plots (C) and heatmaps (D) are shown using matplotlib (see source [point couds](https://osf.io/rykgn), [persistent diagrams](https://osf.io/fyasg), and [code](https://github.com/elalilab/Stroke_PDGFR-B_Reactivity/blob/main/Widefield_20x_ROIs_Pdgfrb_TDA.ipynb)). Note that bottleneck distances are portrayed in Suppl. Figure 7F. The raw plots are available in the “[Communication/Plots/Widefield_20x_ROIs_Pdgfrb_TDA](https://osf.io/jxb4z/)” component of the OSF repository. **E)** Representative images of point clouds derived from PDGFRβ^+^ cells at different filtration values (0, 50, 100, 200) for the healthy peri-lesional region, the injured striatum, and the injured cortex (see entire [image set](https://osf.io/zxbas), source [data](https://osf.io/32a8e), and [code](https://github.com/elalilab/Stroke_PDGFR-B_Reactivity/blob/main/Widefield_20x_ROIs_Pdgfrb_TDA.ipynb)). High-resolution figure: https://osf.io/jxb4z/files/mdre6.

**Figure 6.**
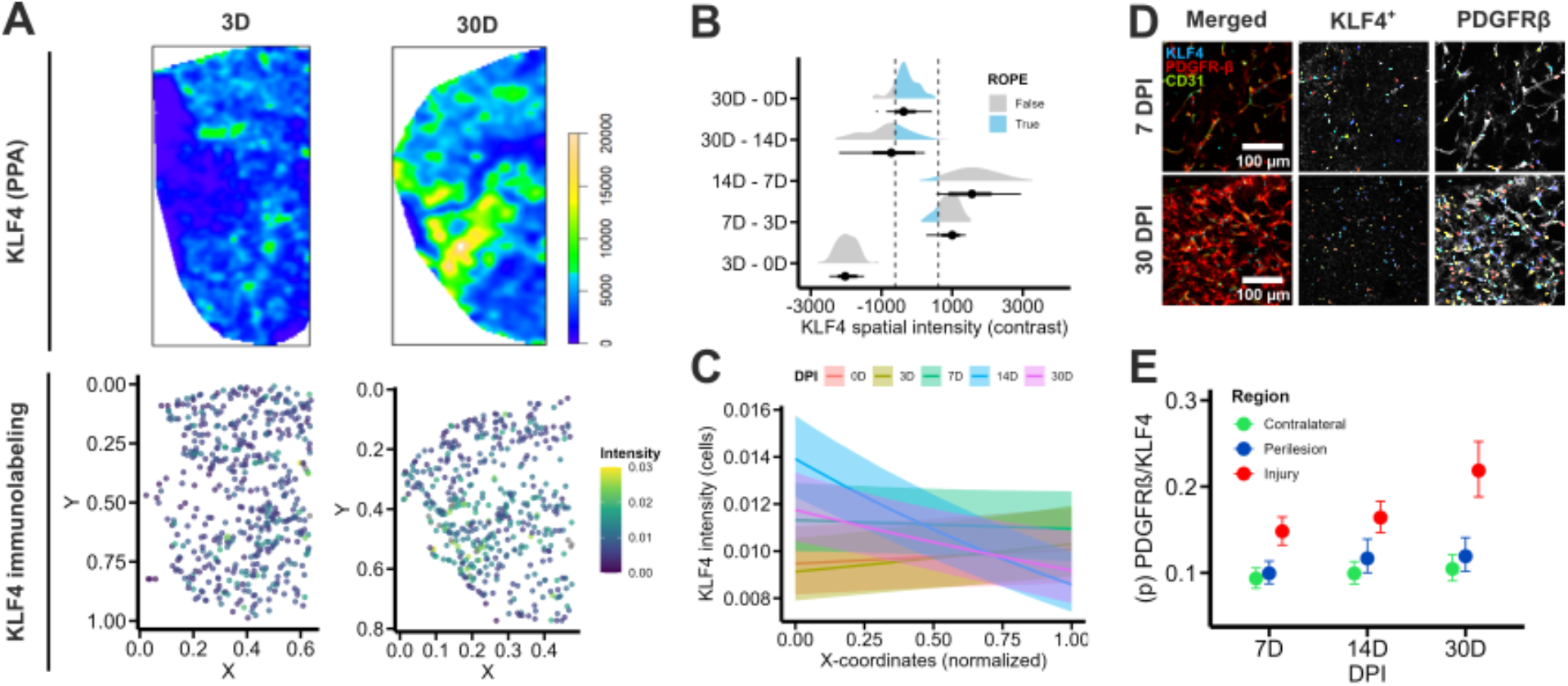
KLF4 overall spatiotemporal regulation after ischemic stroke. **A)** (Upper row) Representative density kernels for KLF4^+^ cells (stitched, 10x magnification) in the ipsilateral hemisphere. The plots are represented using the *topo.colors(256)*scale (see [QN](https://github.com/elalilab/Stroke_PDGFR-B_Reactivity/blob/main/Widefield_10x_Ipsilateral_Klf4_Exp.qmd) and [point patterns](https://osf.io/jdk59)). (Bottom row) Representative plots of single cell KLF4 immunolabeling intensity. Coordinates at 0.0 are the dorsolateral cortex and 0.4 close to the ventricle. The plot is shown using geom_point and scale_color_viridis_c (see entire [image set](https://osf.io/9ngdq)). **B)** Fitted [model](https://osf.io/pq9ck) for Intensity ∼ 0 + DPI, sigma ∼ 0 + DPI, family = student. The posterior probability of contrasts (emmeans) between time points of interest is displayed using add_predicted_draws with stat_halfeye. The ROPE is represented with the shadowed blue region (see Suppl. Figure 8B; Suppl. Table 20; source [data](https://osf.io/gkybe), [code](https://github.com/elalilab/Stroke_PDGFR-B_Reactivity/blob/main/Widefield_10x_Ipsilateral_Klf4_Exp.qmd), and [plot](https://osf.io/mygbz)). **C)** Fitted [model](https://osf.io/q67h4) for Intensity ∼ 0 + DPI * CenterX + (1 | MouseID), family = lognormal. Colored lines and 95% CI per DPI are displayed using conditional_effects. Scaled coordinates at 0.0 are the dorsolateral cortex and 1 close to the ventricle (see Suppl. Figure 8C; Suppl. Table 21; source [data](https://osf.io/vd425), [code](https://github.com/elalilab/Stroke_PDGFR-B_Reactivity/blob/main/Widefield_10x_Ipsilateral_Klf4_Exp.qmd), and [plot](https://osf.io/6ngu5)). **D)** (left) Representative image (20x magnification) of PDGFRβ^tdTomato^/KLF4^+^/CD31^+^ immunolabeling in brain sections at 7 and 30 DPI. (Middle) KLF4^+^ cells detected in CellProfiler with corresponding overlay over PDGFRβ^+^ cells (right) (see the entire [image set](https://osf.io/69m32)). **E)** Fitted [model](https://osf.io/xnzjm) for Colocalized | trials(PDGFRβ) ∼ DPI * Region, family = binomial (7-30 DPI). Posterior estimates are displayed with conditional_effects (mean + 95% CI) (see Suppl. Figure 8D-F; Suppl. Table 23; source [data](https://osf.io/425ed), [code](https://github.com/elalilab/Stroke_PDGFR-B_Reactivity/blob/main/Confocal_20x_ROIs_Klf4-Pdgfrb_Coloc.qmd), and [plot](https://osf.io/mygbz)). High-resolution figure: https://osf.io/jxb4z/files/42tur.

In the striatum, we observed that the connected components merged more rapidly at 7 and 14 DPI (∼120 filtration value) [Wasserstein 0-D: 227, 95% CI = 184 – 316]. Together with the early formation of loops at these time points **(Figure 5C, middle-green/magenta)**, we argue that topological changes in the striatum are evident during the 1^st^ and 2^nd^ week after ischemic stroke. Note, however, that the median distances for the pairs 0-3 DPI [Wasserstein 0-D: 385, 95% CI = 169 – 549] and 3–7 DPI [Wasserstein 0-D: 349, 95% CI = 222 – 785] are larger with considerable uncertainty. We interpret these changes as small-scale interactions, aggregations of few cells that create some subtle non-homogeneities in the tissue. Otherwise, although the merging of connected components at 30 DPI (yellow) is similar to 0 (red) and 3 DPI (blue) **(Figure 5C, middle)**, the late appearance of loops at 30 DPI (1-dimensional homology, yellow) [Wasserstein 1-D: 166, 95% CI = 143 – 183] indicates fewer, but more extensive, interactions between cells. Taken together, the striatum becomes a uniform, large-scale inter-connected monolayer of reactive PDGFRβ^+^ cells between the 1^st^ and 2^nd^ week after ischemic stroke. Conversely, the cortex shows a distinct, decreasing gradient of merging components (0-dimensional homology) over the course of injury progression after stroke **(Figure 5C, right)**. Transitions are larger between 3 and 7 DPI [Wasserstein 0-D: 603, 95% CI = 508 – 669], while the smallest are between 14 and 30 DPI [Wasserstein 0-D: 157, 95% CI = 119 – 225]. This suggests that, in the cortex, a mature fibrotic scar formed by reactive PDGFRβ^+^ cells is steadily established from the 2^nd^ week after injury and undergoes little change thereafter. Compared to the striatum, the maximum merging of connected components (14 and 30 DPI) occurs at ∼75 filtration, indicating a denser arrangement of cells. Together with the analysis of 1-dimensional homology, our results show that the mature scar formed by reactive PDGFRβ^+^ cells (14-30DPI) in the cortex consists of small cell clusters that rapidly form a monolayer at small scales. In **Figure 5E**, we show examples of different filtration values for representative time points per region and bottleneck distances are shown in **Suppl. Figure 7F**.

Overall, our analysis reveals that the topological organization of reactive PDGFRβ^+^ cells evolves more distinctly in the cortex compared to the striatum and the peri-lesional regions. Reactive PDGFRβ^+^ cells in the injured cortex progressively form small-scale cell aggregations that generate a large-scale homogeneous monolayer in the 2^nd^ week after ischemic stroke. At this stage, the fibrotic scar reaches structural maturity beyond which reactive PDGFRβ^+^ cells do not further reorganize.

### 3.7. KLF4 expression is induced at the injured tissue upon ischemic stroke

Previous investigations in different pathological contexts have shown that KLF4 is a critical regulator of the response of peripheral PDGFRβ^+^ cells to hypoxic/ischemic insults (Chandran et al., 2021; Murgai et al., 2017). Therefore, we investigated the expression of KLF4 in our experimental stroke model with emphasis on its regulation in PDGFRβ^+^ cells using immunofluorescence, First, we report that, in our experimental conditions, KLF4 is barely expressed in brain perivascular PDGFRβ^+^ cells but is abundantly expressed in CD31^+^ brain endothelial cells (see immunolabeling protocol and [raw images](https://zenodo.org/api/records/10553084/draft/files/Widefield_10x_Ipsilateral_KLF4-Pdgfrb-CD31.zip/content)). Then, we used PPA and co-localization analysis to evaluate the changes in KLF4 immunoreactivity and its expression in brain PDGFRβ^+^ cells over the course of injury progression. Our PPA analysis in the ipsilateral hemisphere **(see** [QN](https://github.com/elalilab/Stroke_PDGFR-B_Reactivity/blob/main/Widefield_10x_Ipsilateral_Klf4_Exp.qmd)**; Suppl. Method 14; Figure 6A-upper row)** shows that the overall spatial intensity of KLF4 decreases (β_1_ = -0.62) the closer to the intact regions. Nevertheless, modeling of the mean spatial intensity shows that KLF4 immunoreactivity decreases early after injury (3 DPI), and it is progressively restored until the chronic phase (30 DPI) **(Figure 6B; Suppl. Figure 8B; Suppl. Table 20)**.

This suggests that there is no net increase in overall KLF4 expression in the injured brain, but rather expression recovery. Additionally, we modeled the staining intensity per KLF4^+^ cell nuclei using a multilevel model **(see** [QN](https://github.com/elalilab/Stroke_PDGFR-B_Reactivity/blob/main/Widefield_10x_Ipsilateral_Klf4_Exp.qmd)**; Figure 6A-bottom row; Suppl. Figure 8A)**. Our results indicate that the staining intensity of KLF4 increases from 3 to 14 DPI **(Suppl. Figure 8C; Suppl. Table 21)**. We observed that this intensity pattern is also influenced by the position of KLF4^+^ cell nuclei in the x-coordinates (dorsolateral cortex to ventricles) **(Figure 6C; Suppl. Table 22)**, with more intense nuclei near the dorsolateral cortex at 14 [β_1_ = -0.54, 95% CI = -0.67 – -0.41] and 30 DPI [β_1_ = -0.31, 95% CI = -0.47 – -0.15]. Taken together, our data indicate that the number of KLF4^+^ cells does not exceed baseline levels after injury but exhibits a redistribution and marker intensity near the injured region. Next, we evaluated KLF4 expression specifically in PDGFRβ^+^ cells in defined ROIs (contralateral hemisphere, healthy peri-lesional region, and injured cortex) at the time points associated with its upregulation (7, 14, and 30 DPI) ([QN](https://github.com/elalilab/Stroke_PDGFR-B_Reactivity/blob/main/Confocal_20x_ROIs_Klf4-Pdgfrb_Coloc.qmd); **Suppl. Method 15; Figure 6D)**. Quantification of PDGFRβ/KLF4 co-localization is challenging given the anatomical closeness/overlapping of PDGFRβ^+^ cells with the CD31^+^ brain endothelial cells that abundantly express KLF4. For this reason, we used the contralateral hemisphere as a reference, accounting for false-positive co-localization rates. We estimated that this error oscillates between 5 and 10% **(see** [QN](https://github.com/elalilab/Stroke_PDGFR-B_Reactivity/blob/main/Confocal_20x_ROIs_Klf4-Pdgfrb_Coloc.qmd) **and Suppl. Figure 8D-E)**. Our co-localization analysis shows that a maximum of ∼20% of PDGFRβ^+^ cells display detectable KLF4 expression in the injured cortex at 30 DPI [β_1_ = 0.34, 95%CI = 0.15 - 0.53] **(Figure 6E; Suppl. Table 23)**. Considering the estimated co-localization error rate and the false positives that may arise from increased cell density, we conclude that KLF4 expression is barely induced in reactive PDGFRβ^+^ cells and is very subtle compared to that of CD31^+^ brain endothelial cells. **Suppl. Figure 8F** shows the estimated probability of PDGFRβ^+^/KLF4^+^ co-localization relative to the total number of PDGFRβ^+^ cells which is also close to 10% at its maximum when resting the false-positive rate (contralateral hemisphere).

### 3.8. KLF4 induction in PDGFRβ^+^ cells minimally contributes to its overall post-stroke expression

To further assess whether the subtle induced expression of KLF4 in reactive PDGFRβ^+^ cells is biologically significant, we downregulated its expression specifically in PDGFRβ^+^ cells after ischemic stroke onset in PDGFRβ^KLF4-KO^ mice. We chose this strategy to target KLF4 induced expression at its peak, 14 days post-injury (DPI) (see Supplementary Methods 16). First, we obtained the mean spatial intensity using PPA, analogous to our analysis of PDGFRβ^tdTomato^ cells **(Figure 7A)**. The comparison between genotypes [β_1_ = 258, 95%CI = -805 – 1392] **(Figure 7B-C; Suppl. Table 24)** revealed no discernible differences in KLF4 expression in the ipsilateral hemisphere. This indicates that attenuation to KLF4 induced expression after ischemic stroke specifically in PDGFRβ^+^ cells do not largely contribute to its overall expression in the ipsilateral hemisphere. This finding is consistent with our previous results indicating that KLF4 expression is minimal in reactive PDGFRβ^+^ cells and, therefore, not appreciable at the larger scale of the ipsilateral hemisphere. Given the close interrelation between PDGFRβ^+^ cells and CD31^+^ brain vasculature, co-localization studies using anti-PDGFRβ antibodies are non-conclusive (data not shown). We also attempted to sort reactive PDGFRβ^+^ cells from the ipsilateral hemisphere using FACS and MACS approaches; however, we did not achieve a pure suspension that would allow drawing unequivocal conclusions as to the specific subtle changes in KLF4 expression (data not shown), which represents a fundamental limitation of our study. Nonetheless, our results clearly indicate that KLF4 is minimally regulated in PDGFRβ^+^ cells after ischemic stroke.

**Figure 7.**
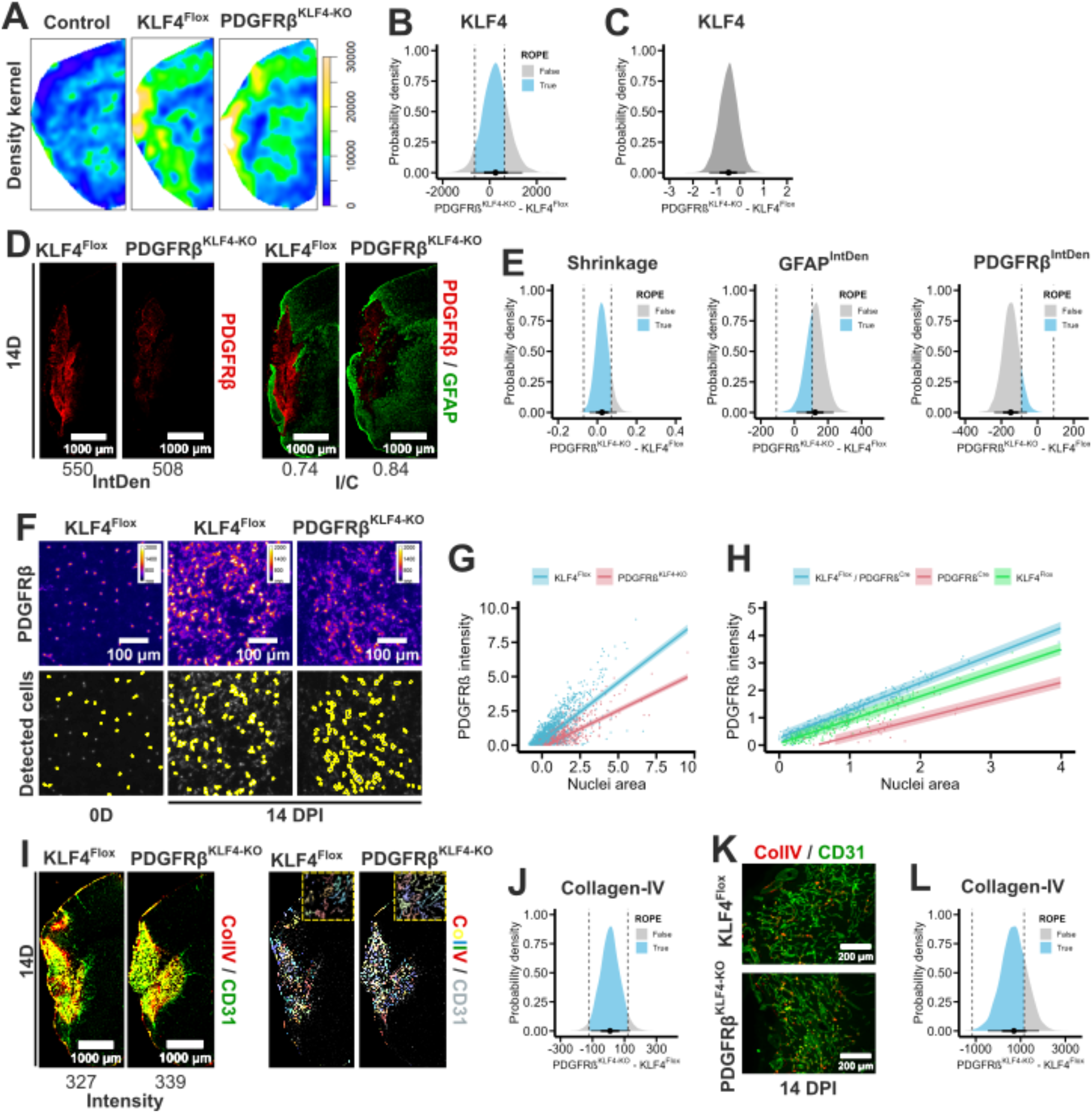
Regulation of KLF4 expression in PDGFRβ^KLF4-KO^ mice. **A)** Representative density kernels for KLF4^+^ cells (stitched, 10x magnification) in the ipsilateral hemisphere of control and PDGFRβ^KLF4-KO^ mice. The plots are represented using the *topo.colors(256)* scale (see [QN](https://github.com/elalilab/Stroke_PDGFR-B_Reactivity/blob/main/Widefield_10x_Ipsilateral_KO_Klf4_Exp.qmd) and [point patterns](https://osf.io/fxgr6)). **B)** Contrast between the spatial intensity of KLF4^+^ cells in the ipsilateral hemisphere of PDGFRβ^KLF4-KO^ mice and corresponding controls for the linear [model](https://osf.io/24ms7) Intensity ∼ Genotype, sigma ∼ Genotype, family = student. Posterior draws (spread_draws) are shown with stat_halfeye() and ROPE (see Suppl. Table 24; source [data](https://osf.io/zhk39), [code](https://github.com/elalilab/Stroke_PDGFR-B_Reactivity/blob/main/Widefield_10x_Ipsilateral_KO_Klf4_Exp.qmd), and [plot](https://osf.io/7b3rf). **C)** Probability density for sigma (unexplained variance) for the model in (B) (see [plot](https://osf.io/k5y7n)). **D)** Representative PDGFRβ (left) and PDGFRβ^+^/GFAP^+^ immunolabeling in brain sections (right) (stitched, 10x magnification) at 14 DPI. Reference values for PDGFRβ IntDen (left) and ratio ipsilateral/contralateral (I/C) (right) are shown for both genotypes (see the entire [image set](https://osf.io/r294m)). **E)** Contrast in hemispheric shrinkage (ipsilateral/contralateral) (left), GFAP IntDen (center) and PDGFRβ IntDen (right) in the ipsilateral hemisphere of PDGFRβ^KLF4-KO^ mice and corresponding controls for the linear models Shrinkage ∼ Genotype, family = student, and IntDen ∼ Genotype, family = student, respectively. Posterior draws (spread_draws) are shown with stat_halfeye() and ROPE (see Suppl. Table 25-27; source [data](https://osf.io/vpxqw), [code](https://github.com/elalilab/Stroke_PDGFR-B_Reactivity/blob/main/Widefield_5x_Ipsilateral_KO_Structure.qmd), and [plots](https://osf.io/vf3n2)). **F)** (upper row) Representative images (10x magnification, ‘fire’ LUT) of ROIs in the ipsilateral cortex labeled for PDGFRβ mRNA (FISH) at 0 and 14 DPI. Yellow represents higher PDGFRβ expression. (bottom row) corresponding cell detections in CellProfiler showing individual PDGFRβ^+^ nuclei (see entire [image set](https://osf.io/ath4m)). **G)** Fitted [model](https://osf.io/vcxez) for Intensity ∼ Area * Genotype + (1 | MouseID), family = student. Colored lines and 95% CI per DPI are displayed using conditional_effects (see Suppl. Table 28; source [data](https://osf.io/jp5bz), [code](https://github.com/elalilab/Stroke_PDGFR-B_Reactivity/blob/main/Widefield_10x_Ipsilateral_KO_CD31-ColIV_Inten.qmd), and [plot](https://osf.io/64scq)). **H)** Fitted [model](https://osf.io/32jy4) for Intensity ∼ Area * Genotype + (1 | MouseID), family = student. Colored lines and 95% CI per DPI are displayed using conditional_effects (see Suppl. Table 32; source [data](https://osf.io/vpxqw), [code](https://github.com/elalilab/Stroke_PDGFR-B_Reactivity/blob/main/Widefield_10x_Ipsilateral_Flox-Cre_Fish_Pdgfrb.qmd), and [plot](https://osf.io/grw8y)). **I)** Representative ColIV^+^/CD31^+^ (left) immunolabeling in brain sections (stitched, 10x magnification) at 14 DPI. Reference values are shown for ColIV total intensity. (Right) ColIV signal (colored) in CD31^+^ endothelial cells, as analyzed in CellProfiler (see the entire [image set](https://osf.io/8av6g)). **J)** Contrast between ColIV intensity in CD31^+^ endothelial cells in the ipsilateral hemisphere of PDGFRβ^KLF4-KO^ mice and corresponding controls for the linear [model](https://osf.io/mw6vt) Intensity ∼ Genotype family = student. Posterior draws (spread_draws) are shown with stat_halfeye() and ROPE (see Suppl. Table 34; source [data](https://osf.io/qnj4m), [code](https://github.com/elalilab/Stroke_PDGFR-B_Reactivity/blob/main/Widefield_10x_Ipsilateral_KO_CD31-ColIV_Inten.qmd), and [plot](https://osf.io/64scq). **K)** Representative images (10x magnification) of ColIV^+^/CD31^+^ immunolabeling in brain sections at 14 DPI from PDGFRβ^KLF4-KO^ mice and corresponding controls (see the entire [image set](https://osf.io/r83ea)). **L)** Contrast between ColIV^+^ area of CD31^+^ vasculature in cortical ROIs of the ipsilateral hemisphere of PDGFRβ^KLF4-KO^ mice and corresponding controls for the linear [model](https://osf.io/q5p93) Area ∼ Genotype family = student. Posterior draws (spread_draws) are shown with stat_halfeye() and ROPE (see Suppl. Table 36; source [data](https://osf.io/ec6wf), [code](https://github.com/elalilab/Stroke_PDGFR-B_Reactivity/blob/main/Widefield_10x_ROIs_KO_CD31-ColIV_Coloc.qmd), and [plot](https://osf.io/kycd3). High-resolution figure: https://osf.io/jxb4z/files/xv4bt.

### 3.9. Specific attenuation of KLF4 expression in PDGFRβ^+^ cells does not affect scar markers

Next, we analyzed structural changes, specifically hemispheric shrinkage, as well as glial and fibrotic scar formation in PDGFRβ^KLF4-KO^ mice and corresponding controls **(Suppl. Method 17; Figure 7D-G)**, as previously described **(see Suppl. Method 1-3)**. Our results showed no differences in brain shrinkage between genotypes [β_1_ = 0.02. 95% CI = -0.04 – 0.09] **(Figure 7E-left; Suppl. Table 25)** but a subtle increase in GFAP IntDen in PDGFRβ^KLF4-KO^ mice [β_1_ = 122. 95% CI = 12 – 233] **(Figure 7E-center; Suppl. Table 26)**. Furthermore, we observed a decrease in PDGFRβ expression in these mice [β_1_ = -148, 95% CI = -239 – -57] **(Figure 7E-right; Suppl. Table 27)**. We further analyzed PDGFRβ IntDen conditioning on shrinkage ratio and ipsilateral area **(Suppl. Figure 9A; Suppl. Table 28)**. These results suggest that PDGFRβ^KLF4-KO^ mice may experience reduced PDGFRβ expression with increasing hemispheric atrophy [β_1_ = -145. 95% CI = -239 – 52]. However, the considerable uncertainty reflected in the credible intervals does not allow us to draw firm conclusions from this association. We further explored the regulation of PDGFRβ in PDGFRβ^KLF4-KO^ mice at the mRNA level using fluorescence in situ hybridization (FISH) **(See Suppl. Method 18)**. Employing unbiased, automated detection methods, we identified individual PDGFRβ^+^ cell nuclei and analyzed their number and fluorescence intensity **(see** [QN](https://github.com/elalilab/Stroke_PDGFR-B_Reactivity/blob/main/Widefield_10x_Ipsilateral_KO_Fish_Pdgfrb.qmd) **and Figure 7F)**. Our analysis indicated no appreciable changes in the number of PDGFRβ^+^ nuclei in PDGFRβ^KLF4-KO^ mice compared to controls [β_1_ = -0.10, 95% CI = -0.38 – 0.16] **(Suppl. Figure 9D-E; Suppl. Table 30)**. However, we observed that the labeling intensity of PDGFRβ^+^ nuclei was reduced in PDGFRβ^KLF4-KO^ mice [β_Area x KO_ = -0.32, 95% CI = -0.34 – -0.29] **(Figure 7G; Suppl. Figure 8E; Suppl. Table 31)**. Furthermore, we found no evidence of altered topological arrangements at the examined scales (ROIs, 10x magnification) resulting from KLF4 attenuated expression in brain PDGFRβ^+^ cells **(see** [QN](https://github.com/elalilab/Stroke_PDGFR-B_Reactivity/blob/main/Widefield_10x_ROIs_KO_Pdgfrb_TDA.ipynb)**; and Suppl. Figure 9F)**. To verify whether the changes in PDGFRβ mRNA intensity were not an artifact of the transgene constructions, we evaluated the same marker in healthy KLF4^Flox^ and PDGFRβ^Cre^ mice **(see** [QN](https://github.com/elalilab/Stroke_PDGFR-B_Reactivity/blob/main/Widefield_10x_ROIs_Flox-Cre_Fish_Pdgfrb.qmd)**; and Suppl Method 19; Suppl. Figure 9G)**. We included PDGFRβ^tdTomato^ mice as controls. Surprisingly, we found that PDGFRβ^tdTomato^ mice exhibited a higher number of nuclei compared to PDGFRβ^Cre^ [β_Cre_ = -51, 95% CI = -78 – -24] but not compared to KLF4^Flox^ [β_Flox_ = -2.5, 95% CI = -33 – 27] **(Suppl. Figure 9H; See Suppl. Table 32)**. Conversely, an analogous analysis of PDGFRβ labeling intensity revealed that both PDGFRβ^Cre^ [β_Area x Cre_ = -0.34, 95% CI = -0.37 – -0.31] and KLF4^Flox^ [β_Area x Flox_ = -0.14, 95% CI = -0.16 – -0.12] mice exhibited lower PDGFRβ mRNA intensity compared to control animals **(Figure 7H; Suppl. Table 33)**. These results imply that the genetic constructs used in our PDGFRβ^KLF4-KO^ mice may alter the endogenous labeling intensity of PDGFRβ in an undetermined manner. This aspect warrants further investigation. We conducted next an additional set of experiments in which we have initiated KLF4 depletion in PDGFRβ^+^ cells prior to experimental stroke onset **(Suppl. Figure 9B-C; Suppl. Table 29A-B)**. In this paradigm, we found no evidence of alterations in PDGFRβ [β_1_ = 18.3, 95% CI = -41 – 74.4] or GFAP expression [β_1_ = -54, 95% CI = -162.4 – 58.1] in PDGFRβ^KLF4-KO^ mice compared to controls. This further confirms that KLF4 does not play major role in regulating the response of PDGFRβ^+^ cells to brain injury after ischemic stroke.

### 3.10. Attenuation of KLF4 post-stroke expression in PDGFRβ^+^ cells does not influence vascular responses

Given the close interaction of a subpopulation of reactive PDGFRβ^+^ cells and brain vasculature, we extended our exploration to CD31^+^ brain endothelial cells. We analyzed vascular reactivity by measuring the intensity of collagen-IV (ColIV) in CD31^+^ cells in the ipsilateral hemisphere after ischemic stroke ([QN](https://github.com/elalilab/Stroke_PDGFR-B_Reactivity/blob/main/Widefield_10x_Ipsilateral_KO_CD31-ColIV_Inten.qmd), **Figure 7I; Suppl. Method 20)**. We observed that attenuation of KLF4 expression in PDGFRβ^+^ cells does not affect ColIV expression in the ipsilateral brain vasculature [β1 = 9.9, 95% CI = - 111 – 136] **(Figure 7J; Suppl. Table 34)**. Furthermore, we performed an alternative analysis of CD31 and ColIV in regions of interest (ROIs) within the injured cortex **(QN, Figure 7K; Suppl. Figure 9A; Suppl. Method 20)**.

We found that the area occupied by CD31^+^ vasculature was similar between PDGFRβ^KLF4-KO^ and controls [β1 = 272, 95% CI = -472 – 999] **(Suppl. Figure 9B-C; Suppl. Table 35)**. However, it is noteworthy that PDGFRβ^KLF4-KO^ mice exhibited greater variability in CD31 reactivity [σ1 = 1.05, 95% CI = 0.23 –1.86] **(Suppl. Figure 9B-right; Suppl. Table 35)**, outlining a potentially more heterogeneous response to ischemic stroke. Similarly, we did not observe differences in the area occupied by ColIV^+^ structures associated with CD31^+^ endothelial cells [β_1_ = 699, 95% CI = -466 – 1825] (**Figure 7L; Suppl. Figure 10A-C; Suppl. Table 35)**. This finding was consistent with the analysis of total collagen in the injured cortex and striatum using Picosirius staining ([QN](https://github.com/elalilab/Stroke_PDGFR-B_Reactivity/blob/main/Widefield_20x_ROIs_KO_Picosirius.qmd), **Suppl. Figure 9D-F)**. Although attenuation of KLF4 expression in PDGFRβ^+^ cells appeared to affect the mean of ColIV^+^ in the injured cortex [β_Ctx-KO_ = 511, 95% CI = 14 – 1573], the considerable uncertainty in this model [σ_1_ = 3974, 95% CI = 3062 – 5150] precludes definitive conclusions **(Suppl. Figure 10F; Suppl. Table 36; Suppl. Table 37)**. Our overall results suggest that although specific attenuation of KLF4 expression in PDGFRβ^+^ cells may augment the variability of injury responses, this does not noticeably affect the mean vascular reactivity.

## 4. Discussion

In this study, we aimed to analyze the reactivity of brain PDGFRβ^+^ cells and its contribution to scar formation in ischemic stroke with emphasis on morphological changes and topological rearrangements over the course of injury. Furthermore, we aimed to elucidate whether KLF4 contributes to the regulation of the reactivity of brain PDGFRβ^+^ cells, as reported for PDGFRβ^+^ peripheral cells. This was achieved using PDGFRβ^tdTomato^ reporter mice and PDGFRβ^KLF4-KO^ transgenic mice subjected to experimental ischemic stroke, implicating the generation of a defined cortico-striatal injury associated with spatiotemporally well-organized progressive scar formation.

First, we analyzed the reactivity patterns of PDGFRβ^+^ cells in the ipsilateral hemisphere using IntDen measurements and PPA. Our findings indicate that the reactivity of PDGFRβ^+^ cells is integral to glial scar formation and is closely associated with brain shrinkage and atrophy in the chronic phase. This observation is consistent with previous studies, which identified, using qualitative approaches, the compartmentalization of PDGFRβ^+^ cells within a GFAP^+^ cell convex hull (Reeves et al., 2019). Herein, we advanced the understanding of this compartmentalization process by providing a quantitative model based on PPA to characterize the distribution of PDGFRβ^+^ cells within the injured tissue in relation to scar-forming GFAP^+^ astrocytes, as previously validated (Manrique-Castano et al., 2024). Our open data and model outputs can be used to validate and strengthen a model-based exploration of scar formation in CNS injuries. Furthermore, our characterization of the reactive PDGFRβ^+^ cells populating the tissue located inside the GFAP^+^ cell convex hull, an area destined for degeneration and liquefaction, warrants deeper exploration. We speculate that at the boundary of GFAP^+^ cell convex hull, there may exist molecular signals demarcating a “no-entry” point for reactive GFAP^+^ astrocytes and limiting extension of the outer astroglial layer (Shibahara et al., 2020). It has been proposed that reactive PDGFRβ^+^ cells could comprise resident perivascular cells, including pericytes and fibroblasts (Dias et al., 2021; Shibahara et al., 2020), as well as infiltrating PDGFRβ^+^ cells (Jung et al., 2011; Kokovay et al., 2006). Further research is warranted to elucidate the specific functions of reactive PDGFRβ^+^ cells and contribution to orchestrating overall scar formation after stroke.

Several transcriptomic studies have outlined the presence of a PDGFRβ^+^ astrocyte subpopulation (Sadick et al., 2022). Some investigations using anti-PDGFRβ antibody suggested that PDGFRβ^+^ astrocytes may contribute to glial scar compartmentalization (Dias et al., 2021; Kyyriäinen et al., 2017). However, it is important to consider that in 2021 the manufacturer indicated that the PDGFRβ antibody used in those studies (Abcam, ab32570, RRID:AB_777165) reacts as well as PDGFRα that is expressed in a subpopulation of perivascular astrocytes as well as fibroblasts (Su et al., 2008). This reclassification promoted an inquiry by our group concerning the labeling specificity of this antibody using the PDGFRβ^tdTomato^ reporter mouse. Therefore, studies utilizing ab32570 antibody may require reevaluation considering anti-PDGFRβ antibody’s updated specificity by the manufacturer. The potential for non-specific labeling of different cell populations underscores the need for careful interpretation of results. However, we acknowledge that the exploration of the role of PDGFRβ^+^ astrocytes using adequate tools and approaches is worth pursuing. For instance, there is evidence suggesting that depletion of PDGFRβ in astrocytes increases vascular leakage and increases infarction after ischemic insults (Shen et al., 2012) suggesting that the subpopulation of PDGFRβ^+^ astrocytes may play important role in maintaining homeostasis of neurovascular interaction in health and disease.

Next, we analyzed the proportion of perivascular and parenchymal reactive PDGFRβ^+^ cells in the injured brain and explored their proliferative potential using Ki67. Our results indicate that brain injury after ischemic stroke prompts the appearance of parenchymal PDGFRβ^+^ cells, which was prominent in the injured cortex. Various brain injury models have reported the detachment of PDGFRβ^+^ pericytes from the vasculature (Dore-Duffy et al., 2000). This process is thought to be controlled by the regulator of G-protein signaling (RGS)5 that is induced by hypoxia to suppress PDGFB-mediated chemotaxis (Enström et al., 2022). Indeed, RGS5^KO^ mice exhibit a reduced density of parenchymal PDGFRβ^+^ cells during the 1^st^ week after ischemic stroke (Roth et al., 2019), which are associated with a “less reactive” phenotype based on cell projections. It has been reported that reactive PDGFRβ^+^ cells detach from the vasculature and functionally reprogram by adopting fibroblast-like characteristics (Dias et al., 2021; Fernandez-Klett et al., 2013; Shibahara et al., 2020). Other reports have indicated that PDGFRα^+^ perivascular fibroblasts, which express as well PDGFRβ, are the major cell source contributing to the fibrotic reaction in various brain injuries, including stroke (Dorrier et al., 2022). In our PDGFRβ^tdTomato^ reporter mice, we did not observe tdTomato expression with PDGFRα^+^ cells detected using the Abcam antibody (ab203491, RRID:AB 2892065) in the healthy brain. The PDGFRα antibody labeled ramified astrocyte-like cells in the parenchyma but did not label PDGFRβ^tdTomato^ perivascular cells. Based on our observations, it would be important to consider the specificity of antibodies used in identifying these populations in future studies.

Detachment of reactive PDGFRβ^+^ cells is often reported to be accompanied by a remarkable proliferative activity (Özen et al., 2014). Herein, we aimed to further elucidate this aspect using PDGFRβ^tdTomato^ reporter mice. We observed an increased proportion of reactive PDGFRβ^+^ parenchymal cells at the injured tissue, which cannot be completely linked to a detachment from the vasculature and subsequent migration into the parenchyma. In PDGFRβ^tdTomato^ reporter mice, in which tdTomato expression and relative intensity depends upon PDGFRβ basal expression and activity, we reported an increased intensity at time points at which detachment and parenchymal recruitment were suggested to peak (Fernandez-Klett et al., 2013). Furthermore, upon prolonged Cre-recombination, we observed the appearance of a population of ramified cells that express low levels of tdTomato throughout the brain parenchyma of PDGFRβ^tdTomato^ reporter mice, indicative for low PDGFRβ basal expression and activity, which could represent the subpopulation of PDGFRβ^+^ astrocytes. It is plausible that the emergence of PDGFRβ^+^ parenchymal cells might be due to an increased marker expression in already existing non-perivascular populations, as reported (Bernard et al., 2024; Renner et al., 2003), such as PDGFRβ^+^ astrocytes, rather than vascular detachment *per se*.

Moreover, the contribution of PDGFRβ^+^ immune-like cells derived from the periphery that infiltrate the injured tissue via the affected vasculature cannot be excluded (Jung et al., 2011; Kokovay et al., 2006). In this regard, some reports have indicated that perivascular cells co-expressing PDGFRβ and CD13, identified as pericytes, leave the vasculature, proliferate and express microglial markers the 1^st^ week after ischemic stroke (Özen et al., 2014). It is highly probable that PDGFRβ^+^/CD13^+^ cells might be infiltrating PDGFRβ^+^ immune cells that are transmigrating through the destabilized vasculature, which is not adequately covered by resident perivascular cells, including pericytes. Although CD13 is expressed in brain pericytes under normal conditions, it is expressed in myeloid cells that are recruited to the angiogenic vessels after ischemic stroke (Nguyen et al., 2023b). Thereafter, it would be important to take into consideration these aspects in data interpretation, and additional investigations are required to better characterize the different reactive PDGFRβ^+^ cell populations after ischemic stroke and other brain injuries. Detachment of PDGFRβ^+^ cells have been reported to coincide with a considerable number of reactive cells exhibiting proliferation markers, such as BrdU or Ki67, in ischemic stroke (Fernandez-Klett et al., 2013; Özen et al., 2014) as well as in other brain injuries (Zehendner et al., 2015). However, utilizing an unbiased, automated, and reproducible workflows, we found that the proportion of reactive PDGFRβ^+^ cells expressing Ki67 is extremely low and cannot account for the increased overall density of cells observed at the injured tissue over time. This holds true for both vascular-associated and parenchymal PDGFRβ^+^ cells. This discrepancy could be due to the antibody-based approaches used in previous studies to label PDGFRβ^+^ cells. To promote transparency and facilitate further research, we provide the raw data and analysis code for validation by the scientific community. Unfortunately, we were unable to obtain comparable evidence using BrdU, as its incorporation requires a denaturation process that significantly damages tdTomtao signal in PDGFRβ^+^ cells, rendering the analysis unreliable.

Next, we characterized the morphological changes of PDGFRβ^+^ cells in both the injured and intact regions after stroke. Consistent with previous reports (Menezes et al., 2020; Özen et al., 2014), our PDGFRβ^tdTomato^ reporter mice label perivascular cells co-expressing CD13 in the healthy regions. However, we found no convincing evidence that these cells co-express GFAP, IBA1, or PDGFRα in the injured tissue, in contrast to findings reported by other studies. We argue that the close interaction between neighboring cells at the CD31^+^ vasculature, such as PDGFRα^+^ cells (Vanlandewijck et al., 2018), or the extensive superposition of reactive PDGFRβ^+^ cells and IBA1^+^ microglia in the injured regions (Riew et al., 2018), may lead to misinterpretations, especially in the absence of unbiased and reproducible workflows. We acknowledge that we did not perform a quantitative, unbiased analysis of co-localization with these markers, as the increased cell density and staining artifacts in the injured tissue rendered the analysis unreliable. This limitation highlights the need to develop reproducible workflows for segmentation, reconstruction, and object-based co-localization studies. Furthermore, we postulate that our morphological analysis has quantitatively characterized the amoeboid PDGFRβ^+^ cells described by Zehendner et al. (2015), fibroblast-like cells (reticulite) reported by Dias et al. (2021) and Göritz et al. (2011), and reactive RGS5^+^ cells (reticulite) identified by Özen et al. (2014). Our raw data (images) and statistical models provide a robust foundation for future investigations into the morphological changes of the diverse population of PDGFRβ^+^ cells particularly the ambiguous cell types such as the reticuloparenchymal cells. A significant strength of our analysis is that it offers reproducible and reusable workflows for cell morphological analysis based on open-source tools (Python). We encourage fellow scientists to leverage our workflows to improve cell segmentation models. Additionally, this material can serve as input or priors to validate and train machine learning-based cell classifiers.

Subsequently, we analyzed the redistribution and topological arrangements of reactive PDGFRβ^+^ cells over injury progression. We found that in addition to the changes of PDGFRβ expression and activity, PDGFRβ^+^ cells undergo a predictable rearrangement pattern. Traditionally, analyses of cell reactivity and glial scar formation have relied on qualitative assessments that disregard spatial insights. Herein, we implemented PPA and TDA, as insightful frameworks for quantifying the distribution of point patterns or point clouds derived from individually detected cells (Manrique-Castano et al., 2024). TDA and PPA have broad applications in both biological (Ben-Said, 2021; Bhaskar et al., 2023; McGuirl et al., 2020) and non-biological sciences (Baddeley et al., 2021; Bilotti et al., 2024). Our approach utilizing these tools offers a viable alternative to surpass raw cell counts or density estimations that lack comprehensive spatiotemporal information. In this article, we extended our previous analyses of NeuN^+^, GFAP^+^, and IBA1^+^ cells (Manrique-Castano et al., 2024) to include PDGFRβ^+^ cells and provided expanded, reproducible workflows. These incorporate the analysis of Haralick features (Löfstedt et al., 2019), the implementation of bootstrapping (Chernik, 2007b) to analyze Wasserstein and bottleneck distances (Hajij et al., 2018), and the generation of Vietoris-Rips complexes at different filtration values (Chambers et al., 2010). We provide complete and reusable workflows using Python and related libraries, along with entire sets of point patterns (.R objects) and point clouds (.npy) for further analysis of the spatial features of cells. An important limitation of our study is the identification of PDGFRβ^+^ cells using CellProfiler and QuPath, particularly in regions with high cell density. This inaccuracy may be influenced by multiple factors, including labeling patterns, cell morphology, and significant cell overlaps. These constraints are especially pronounced when dealing with ramified and variable-intensity cells, as current algorithms are primarily optimized for detecting round-shaped cells (Al-Kofahi et al., 2018, 2018; Stringer et al., 2021). Consequently, our approach estimates local cell density but is limited in its ability to precisely quantify the absolute number of cells within ROIs or time windows. We acknowledge that the segmentation of ramified cells remains a significant challenge that requires further advancement, particularly through machine learning-based methods that leverage morphological features for cell segmentation. Addressing this limitation represents an important avenue for future research.

We next aimed to investigate the mechanisms underlying the transition of quiescent PDGFRβ^+^ cells towards a reactive state upon injury. We were particularly interested in KLF4, a transcription factor that has been reported to be critically involved in regulating the activation of peripheral PDGFRβ^+^ cells to de-differentiate and reprogram to adopt fibrotic-like properties in response to hypoxic or ischemic insults (Chandran et al., 2021; Shankman et al., 2015b; Sheikh et al., 2015). The Allen Brain Atlas protein map (https://mouse.brain-map.org/gene/show/16373) shows that KLF4 is expressed mainly in neurogenic zones such as the olfactory bulb in the healthy adult brain. Murgai et al., (2017) demonstrated that KLF4 expression maintains a pluripotent state in pericytes, leading to enhanced ECM production. Bulut et al., (2021) showed that KLF4 depletion in smooth muscle cells (SMCs) reduces infiltration of pro-inflammatory macrophages in the inflamed adipose tissue. KLF4 expression in SMC was reported to modulate cell transition to a macrophage-like state upon heart ischemia-reperfusion injury (IRI) (Haskins et al., 2018). In the context of ischemic stroke, KLF4 in brain endothelial cells alleviates vascular inflammation (X. Zhang et al., 2020). Furthermore, KLF4 expression in S100A astrocytes attenuates an A1 neurotoxic polarization to promote an A2 neuroprotective phenotype after ischemic stroke (C. Wang & Li, 2023). While some reports have indicated that KLF4 expression could be detected in PDGFRβ^+^ pericytes after ischemic stroke and contribute to cell reprogramming (Nakagomi et al., 2015), its role in regulating the reactivity of PDGFRβ^+^ cells remains completely unknown.Using the anti-KLF4 antibody from Abcam (ab214666, RRID:AB_2943042) on PFA-fixed healthy brain tissue, we observed a stable abundant nuclear KLF4 signal in CD31^+^ brain endothelial cells. This observation aligns with data from protein atlases, which indicate that KLF4 is mainly expressed in brain vascular structures (see <u>The Human Protein Atlas</u>). We obtained similar results in tests performed on glyoxal-fixed brains (data not shown). Zhang et al. (2020) have reported that KLF4 expression is scarce in the healthy brain, using the anti-KLF4 antibody ab129473 from Abcam, and that it is only upregulated upon injury. We suggest that this discrepancy may be due to differences in antibody efficiency to label KLF4 in brain endothelial cells. Additionally, we did not observe an increased KLF4 expression in the ischemic core the first 4 days post-stroke. Instead, our analysis indicate that KLF4 immunoreactivity is slightly reduced in the ischemic core 3 days after stroke, which is consistent with the loss of endothelial cells (Chen et al., 2019) and pericytes (Fernandez-Klett et al., 2013), and is restored during the chronic phase to reach similar levels as in the healthy brain. While we did observe a redistribution of KLF4 towards the injured tissue, there was no overall meaningful increase in KLF4 expression post-stroke. Thus, we highlight the importance of antibody selection and methodological considerations in detecting KLF4 expression and call for the availability of raw data to contrast or validate previous results. Nonetheless, we acknowledge that regulation of KLF4 specific expression in brain endothelial cells after ischemic stroke warrants further investigations using co-localization analysis. Furthermore, we did not observe KLF4 expression pattern in S100^+^ astrocytes reported by Wang & Li (2023), in which anti-KLF4 antibody ab129473 was used. Although we did not specifically investigate S100^+^ astrocytes, our approach did not reveal any staining patterns beyond nuclear localization, which is typical for transcription factors like KLF4 in CD31+ brain endothelial cells. Furthermore, we argue that co-localization between KLF4 (which exhibits nuclear staining) and GFAP (labeling major cytoskeletal branches) is not feasible using unbiased and reproducible methods. We consider it crucial to re-examine this aspect when investigating KLF4 and have shared all our raw data and protocols to facilitate future research leveraging our approach. Importantly, our findings indicate that KLF4 does not play a major role in regulating the reactivity of brain PDGFRβ^+^ cells upon ischemic stroke, as reported for peripheral PDGFRβ^+^ cells under hypoxic or ischemic conditions (Chandran et al., 2021; Shankman et al., 2015b; Sheikh et al., 2015). We did not observe meaningful changes in the progression of the injury or in the overall structure of glial scar upon KLF4 conditional specific depletion in PDGFRβ^+^ cells either prior to ischemic stroke onset or at its expression peak at 14 DPI. Likewise, our results reveal that vascular ColIV and total collagen expression remain unchanged in PDGFRβ^KLF4-KO^ mice. In contrast to the findings that have outlined KLF4’s role regulating the reactivity PDGFRβ^+^ cells in peripheral organs, which was supposed to be conserved in all PDGFRβ^+^ cells, our observations indicate that KLF4 does not fulfill the same functions in brain PDGFRβ^+^ cells. This difference could be due to the distinct origin of brain PDGFRβ^+^ cells that are mainly derived from the neuroectoderm, whereas peripheral PDGFRβ^+^ cells originate from the mesoderm (Korn et al., 2002b). A limitation of our study is the inability to perform an unbiased, single-cell analysis of KLF4 due to the close interaction between CD31^+^ and PDGFRβ^+^ cells, and the high cell density in injured tissue. Future studies employing single-cell RNA sequencing may provide deeper insights into the potential regulatory roles that KLF4 might exert on PDGFRβ^+^ cell responses. Additionally, our findings underscore the need to reconsider the approaches that investigate KLF4 and PDGFRβ using immunohistochemistry, especially following CNS injuries.

Finally, one scope of our study is the use of model-based Bayesian inference with brms and MCMC, which allows us to obtain precise parameters/coefficients with their respective uncertainties. We are convinced that our scientific inference based on effect sizes and full posterior distributions (Kruschke & Liddell, 2018), which draws on a coherent family of distributions and statistical (linear and nonlinear) models, provides a more informative approach than threshold-based analysis based on the uninformative notion of significance. We make our statistical models (as .R objects) and raw data available as open source to promote their reproducibility and validation in further research. We are convinced that our models can be used as priors or quantitative information for research in the field of brain injuries.

## 5. Conclusion

Herein, our study shows that brain PDGFRβ^+^ cells actively respond to injury after ischemic stroke to form an inner fibrotic core at the injured tissue surrounded by GFAP^+^ reactive astrocytes in a predictable (model-based) manner. We postulate that reactive PDGFRβ^+^ cells are mainly early allocated to regions designated to irreversible damage. Brain injury triggers profound morphological changes in PDGFRβ^+^ cells and the emergence of a parenchymal PDGFRβ^+^ cell population, which remains to be characterized. Our findings suggest that the increased density of reactive PDGFRβ^+^ cells might not be the simple result of vascular detachment, proliferation, and migration, of resident cells, thus providing a new framework to further investigate the specific functions of the diverse subpopulations of reactive PDGFRβ^+^ cells. We show that PPA or TDA, with point clouds or point patterns as input, are suitable strategies to model the redistribution patterns of reactive PDGFRβ^+^ cells and to analyze their location with respect to covariates such as brain coordinates or GFAP^+^ astrocytes. Finally, our results indicate that in KLF4 does contribute to the regulation of brain PDGFRβ^+^ cell response to ischemic stroke. Finally, we are making the raw images/data, analysis code and statistical models available for reproducibility and validation of our findings.

