## Supplementary figures for "Brain PDGFRβ^+^ cells exhibit diverse reactive phenotypes after stroke without requiring KLF4"

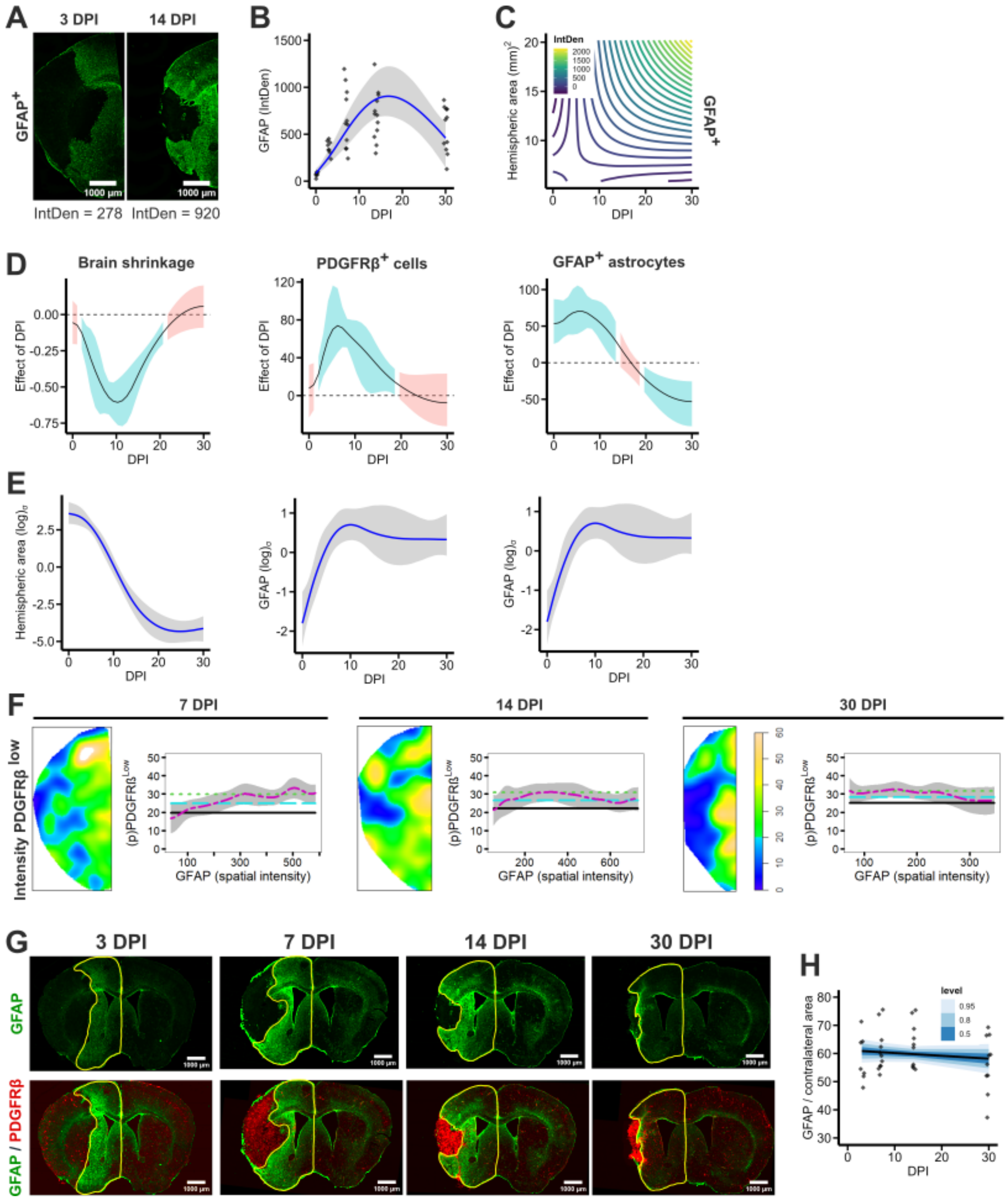

**Supplementary Figure 1. Reactivity of PDGFRβ$^+$ and GFAP$^+$ cells after ischemic stroke. A)** Representative brain section (stitched, 5x magnification) at 3 and 14 DPI showing the astroglial scar formed by GFAP$^+$ astrocytes. Integrated density (IntDen) values are shown at the button as a reference. **B)** Fitted splines model for $GFAP_{IntDen}$ ~ s(DPI). The conditional_effects are shown with the mean (blue) and 95% CI (gray-shadowed region) and observations as diamonds. **C)** Fitted splines for $GFAP_{IntDen}$ ~ t2(DPI, Area). The conditional_effects are displayed with a Viridis scale (IntDen). **D)** Model derivatives calculated with modelbased for the non-linear models fitted for brain shrinkage (left), PDGFRβ (middle), and GFAP (right). The curves show the effect of each DPI (positive or negative) in the response variable. **E)** Display the sigma ($\sigma$) coefficients and their respective uncertainty for the fitted models for brain shrinkage (left), PDGFRβ (middle), and GFAP (right). The results are shown in the log scale. **F)** PDGFRβ$^{low}$ density kernels (left) at 7, 14, and 30 DPI. The density kernels are represented using the topo.colors(256) scale. The relative distribution diagrams (right) depict the probability of the allocation of PDGFRβ$^{low}$ cells (y-axis) relative to GFAP spatial intensity (x-axis). The pooled mean intensity (cyan-dashed line) is shown with their corresponding lower (black-solid line) and upper (green-dotted line) limits of the two-sigma confidence intervals. Note the dissimilar *xy* axes limited by the mapping of the rhohat function for each DPI. Diagrams for PDGFRβ$^{high}$ are displayed in Figure 2 H-I. **G)** Representative GFAP$^{+}$/PDGFRβ$^{tdTomato}$ immunolabeling in brain sections (stitched, 5x magnification) showing the GFAP$^{+}$ cell convex hull (yellow contour). Reactive PDGFRβ$^{+}$ cells allocate primarily outside these boundaries. **H)** Fitted linear model for GFAP / contralateral area ~ DPI. We display stat_lineribbon() with the mean hemispheric area (black) and different credible intervals: 0.05, 0.8, 0.95 (blue brewer scale). The observations are displayed as black diamonds. High-resolution figure: https://osf.io/jxb4z/files/ztfb6.

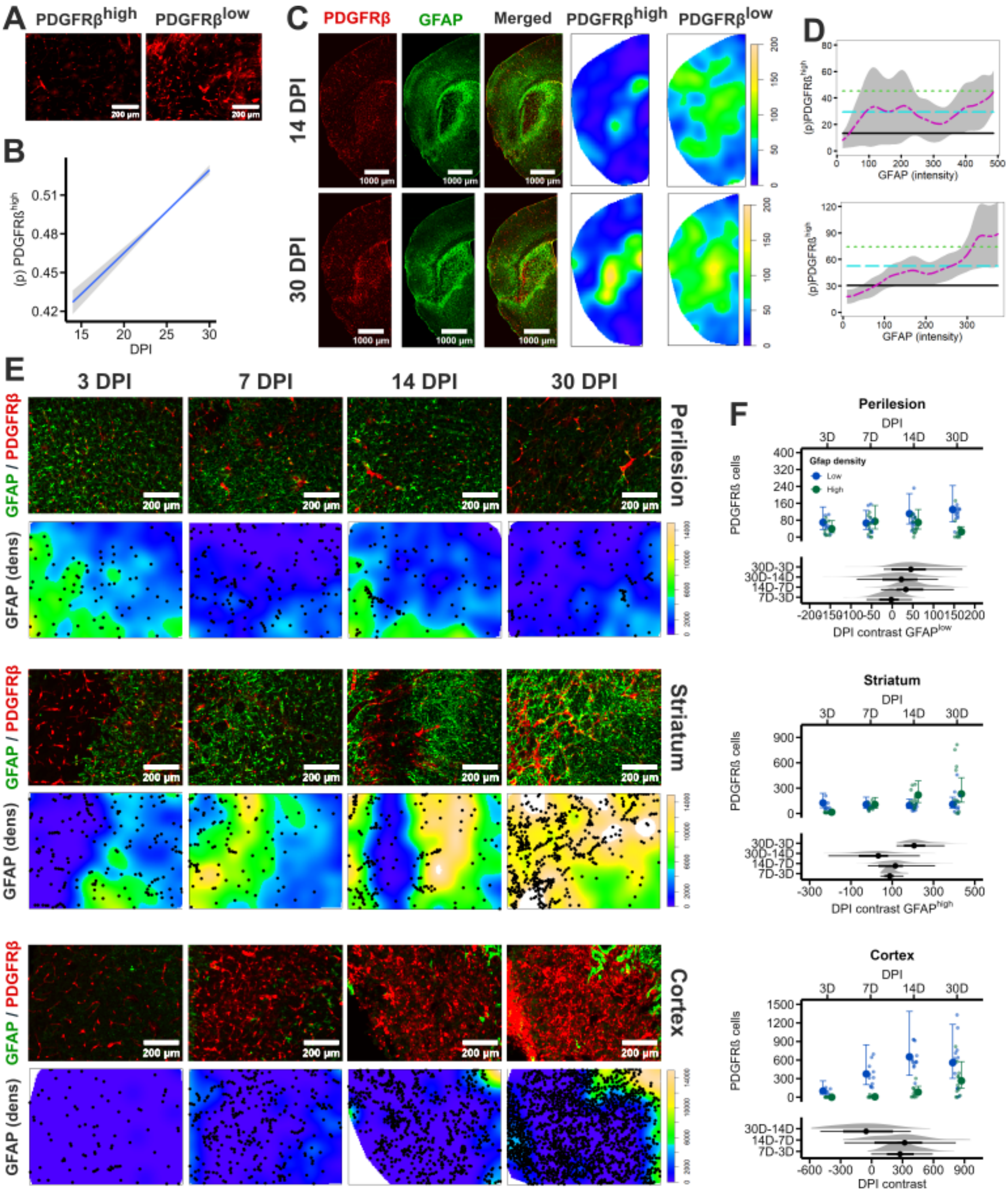

**Supplementary Figure 2. Distribution of PDGFRβ$^+$ cells upon striatal-specific injury**. **A)** Sample crops of PDGFRβ$^{low}$ (non-reactive) and PDGFRβ$^{high}$ (reactive) cells in the ischemic hemisphere of animals with ischemic stroke-mediated striatal-specific injury. **B)** Fitted binomial model for PDGFRβ$^{high}$ | PDGFRβ$^{total}$ ~ DPI in mice with striatal-specific injury. The conditional_effects are displayed with the mean (blue) and 95% CI (gray-shadowed region). **C)** Representative PDGFRβ$^{tdTomato}$/GFAP$^+$ immunolabeling in brain sections (left) (stitched, 5x magnification) with corresponding PDGFRβ$^+$ cell density kernels (right) at 14, and 30 DPI in animals with striatal-specific injury. The density kernels are represented using the *topo.colors(256)* scale. **D)** Relative distribution diagrams (*rhohat*) depicting the allocation probability of PDGFRβ$^{high}$ cells (y-axis) relative to GFAP spatial intensity (x-axis) in animals with striatal-specific injury. The pooled mean intensity (*cyan-dashed line*) is shown with their corresponding lower (*black*-solid line) and upper (*green-dotted line*) limits of the two-sigma confidence intervals. Note the dissimilar *xy* axes limited by the mapping of the rhohat function for each DPI. **E)** Representative GFAP$^{+}$/PDGFRβ$^{tdTomato}$ immunolabeling in brain sections (upper rows) (10x magnification) with corresponding GFAP density kernels (bottom rows) at 3, 7, 14, and 30 DPI in the healthy peri-lesional region, the injured striatum, and the injured cortex. The density kernels are represented using the *topo.colors(256)* scale and PDGFRβ$^{tdTomato}$ cells as black dots. **F)** Fitted models for PDGFRβ ~ Region. The conditional_effects() with mean cell counts and 95% CI are displayed as whiskers. At the bottom, entire posterior densities for the contrast between DPI are shown with stat_pointinterval. High-resolution figure: https://osf.io/jxb4z/files/3q8pm.

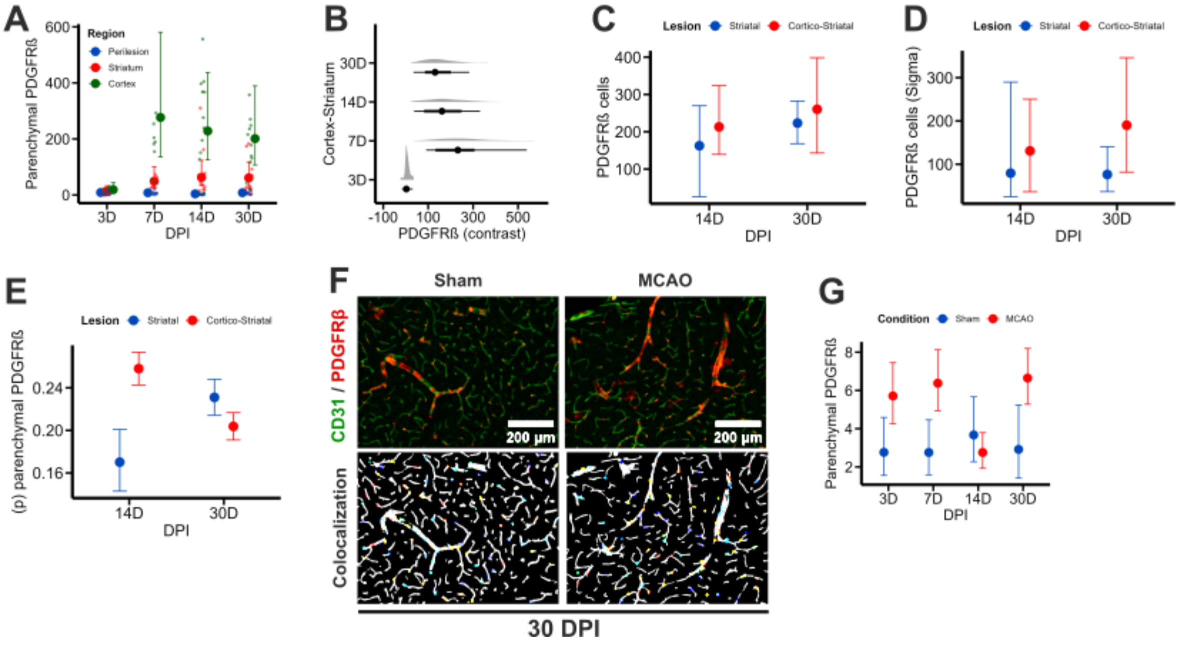

**Supplementary Figure 3. Emergence of a subpopulation of parenchymal PDGFRβ$^+$ cells after ischemic stroke. A)** Fitted model (Pdgfrb_Total ~ DPI * Region, family = hurdle_lognormal) for the total number of parenchymal PDGFRβ$^+$ cells in the peri-lesional region, the injured striatum, and the injured cortex. Colored dots are observations. **B)** Contrast between the number of PDGFRβ$^+$ cells in the injured striatum and the injured cortex. After vanishing at 3 DPI in the cortex, the number of PDGFRβ$^+$ increases subsequently. **C-D)** Fitted model (Pdgfrb_Total ~ DPI * Lesion, sigma ~ DPI * Lesion, family = student) for the comparison between striatal-specific (striatal) and cortico-striatal injuries at 14 and 30 DPI. **E)** Fitted model (Pdgfrb_Parenchymal | trials(Pdgfrb_Total) ~ DPI * Lesion, family = binomial) for the proportion of parenchymal PDGFRβ$^+$ cells in striatal-specific (striatal) and cortico-striatal injuries at 14 and 30 DPI. **F)** Representative images (10x magnification) of PDGFRβ$^{tdTomato}$/CD31$^+$ immunolabeling in brain sections (upper row), and co-localization analysis (CellProfiler) of nuclei in PDGFRβ$^{tdTomato}$ cells (colored) and CD31$^+$ vasculature (white) (bottom row) for sham and ischemic stroke animals (MCAo). **G)** Fitted model (Pdgfrb_Parenchymal ~ DPI * Condition, family = student) to study the impact of Cre-recombination on the number of parenchymal PDGFRβ$^+$ cells in the healthy cortex. Prolonged Cre-recombination does not affect the emergence of parenchymal PDGFRβ$^+$ cells. **A,C-E,G)**. The posterior estimates are displayed with conditional_effects (mean + 95% CI). High-resolution figure: https://osf.io/jxb4z/files/ywnez.

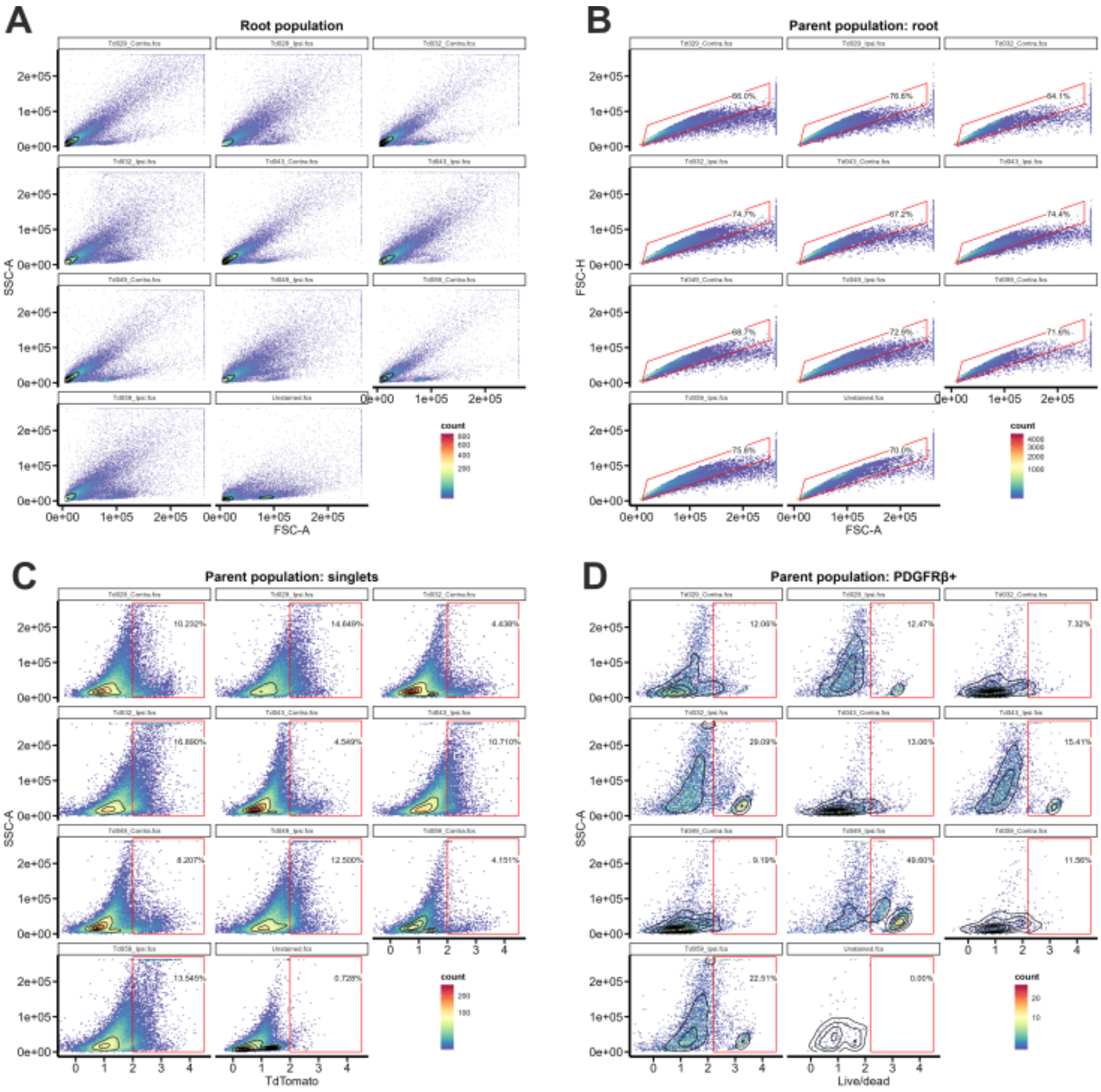

**Supplementary Figure 4. Sorting of brain PDGFRβ⁺ cells after stroke. A)** FSC-A (forward scatter area) and SSC-A (side scatter area) from ipsilateral (ipsi) and contralateral (contra) hemispheres at 14 DPI. **B)** Singlet gating, **C)** tdTomato (PDGFRβ$^+$ cells) gated from singlets and, D) dead cells (Live/dead$^+$ cells) from PDGFRβ$^+$ cells. **A-D)**. The graphs are plotted using geom_hex (bins), geom_density2d (black contour), geom_gate (red gated area), geom_stats (percentage of cells in gated regions). High-resolution figure: https://osf.io/jxb4z/files/zxku4.

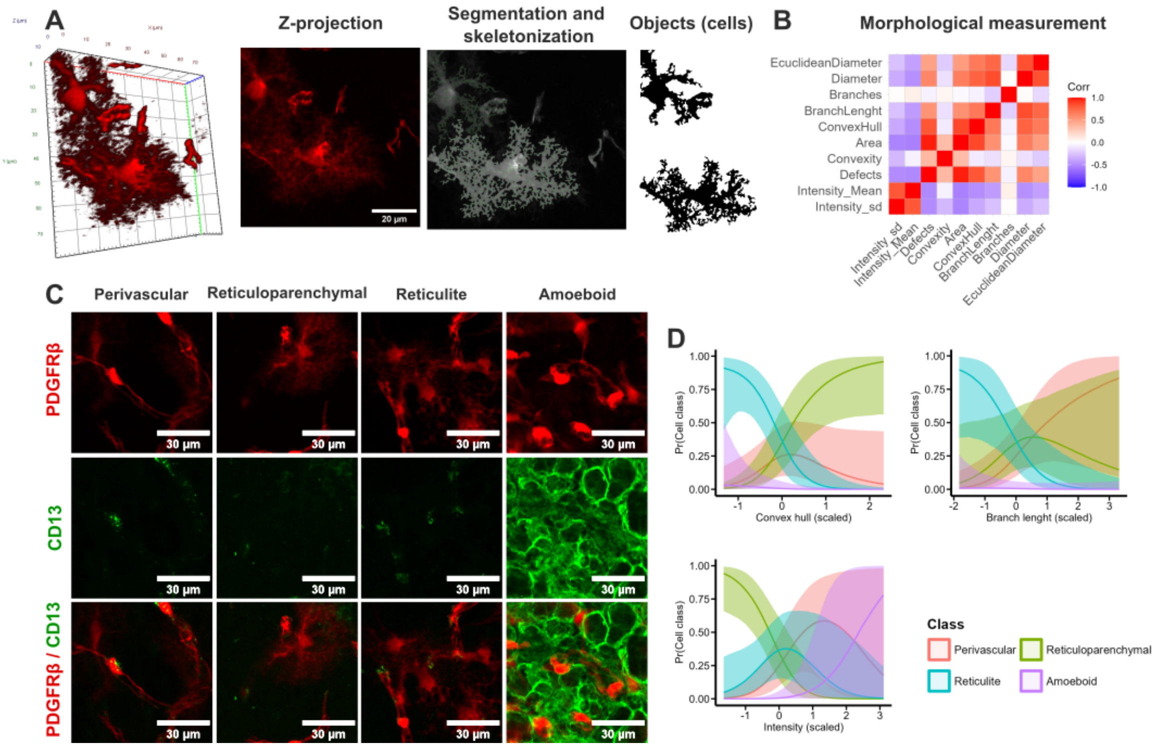

**Supplementary Figure 5. Processing of brain PDGFRβ⁺ cells for morphological analysis. A)** Laser scan confocal microscope was used to acquire z-stacks at 40x magnification of PDGFRβ$^+$ cells in the cortical peri-lesional region, the injured striatum and the injured cortex. The z-stacks were projected in a single plane (2D cells), and images were processed using Python libraries skimage and scipy to enhance, threshold, and generate masks of single cells. Then, the morphological characteristics were measured using regionprops and regionprops_table functions from skimage.measure and data exported to R for principal component analysis (PCA). **B)** Correlation plot of the morphological features used for PCA in the ggcorrplot package. The features are defined in the respective QN notebook. **C)** Perivascular PDGFRβ$^+$ (tdTomato) cells exhibit changes in the immunoreactivity to CD13 upon injury. CD13 immunoreactivity was not detected in reticuloparenchymal PDGFRβ$^+$ cells in the healthy tissue. The meaningful induction of CD13 immunoreactivity in ischemic regions (amoeboid, right column) hinders the quantitative, unbiased analysis of PDGFRβ$^{tdTomato}$/CD13$^+$ immunolabeling co-localization. **D)** Logistic regression plots for the model *Class ~ ConvexHull + BranchLenght + Intensity_Mean, family = categorical*. The conditional_effects are displayed with means and 95% CI for the probability of perivascular (red), reticuloparenchymal (green), reticulate (cyan), and amoeboid (magenta) cells conditioning on defined morphological features (Convex hull, branch length, and intensity). Please note that all morphological traits have been scaled during PCA analysis. High-resolution figure: https://osf.io/jxb4z/files/famn2.

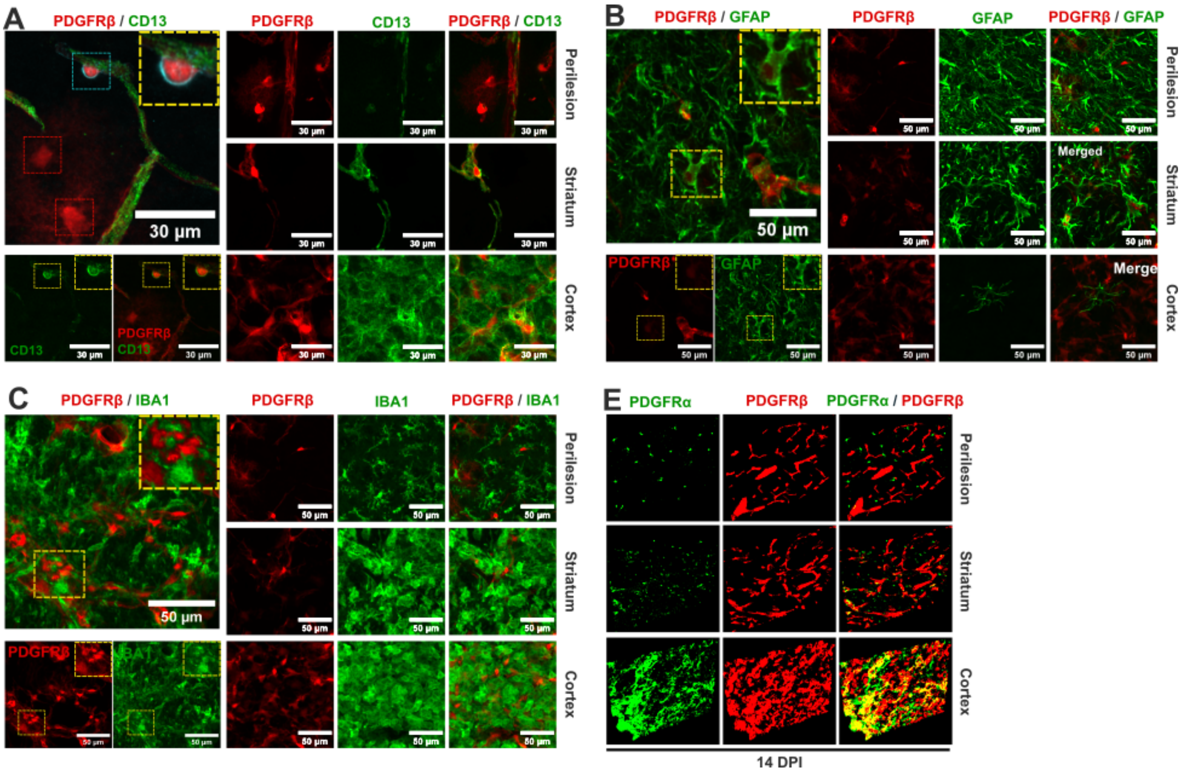

**Supplementary Figure 6. Profiling expression of CD13, GFAP, IBA1, and PDGFRα in brain PDGFRβ$^{tdTomato}$ cells.** **A)** Representative laser scan confocal images of PDGFRβ$^{tdTomato}$/CD13$^{+}$ immunolabeling in brain sections at 40x magnification in the perilesional cortex (perilesion), the injured striatum and the injured cortex. The upper panel on the left shows perivascular PDGFRβ$^{tdTomato}$/CD13$^{+}$ cells (zoomed in the yellow squared section). In contrast, reticuloparenchymal PDGFRβ$^{+}$ cells do not co-express CD13. The patterns of CD13 immunoreactive change in the striatum and cortex following injury. In particular, the ischemic cortex shows increased immunoreactivity to CD13. **B)** Confocal images of PDGFRβ$^{tdTomato}$/GFAP$^{+}$ immunolabeling in brain sections at 20x magnification in the same ROIs, as previously described. GFAP$^{+}$ cell branches appear to wrap some PDGFRβ$^{tdTomato}$ nuclei in regions of strong astrogliosis. However, the origin of GFAP$^{+}$ branches is ambiguous. **C)** Confocal images of PDGFRβ$^{tdTomato}$/IBA1$^{+}$ immunolabeling in brain sections at 20x magnification in the same ROIs, as previously described. PDGFRβ$^{tdTomato}$/IBA1$^{+}$ cells were not observed in the healthy brain and IBA1 increased immunoreactivity in the injured tissue made an unbiased and reproducible analysis impracticable. **D)** Widefield images (with apotome optical sectioning) of PDGFRβ$^{tdTomato}$/PDGFRα$^{+}$ immunolabeling in brain sections at 20x magnification. PDGFRβ$^{+}$ and PDGFRα$^{+}$ cells constitute two distinct populations in the healthy brain. Following injury, both cell types react by increasing the marker intensity and covered area. Although both markers partially co-localize in some regions, undefined cellular morphology and non-specific nuclear staining make co-localization analysis challenging with the tools at our disposal. **A-D)** In all cases, unbiased and automated colocalization with PDGFRβ$^{tdTomato}$ cells is ambiguous given the high cell density and uncertain nuclei labeling. High-resolution figure: https://osf.io/jxb4z/files/ebfdw.

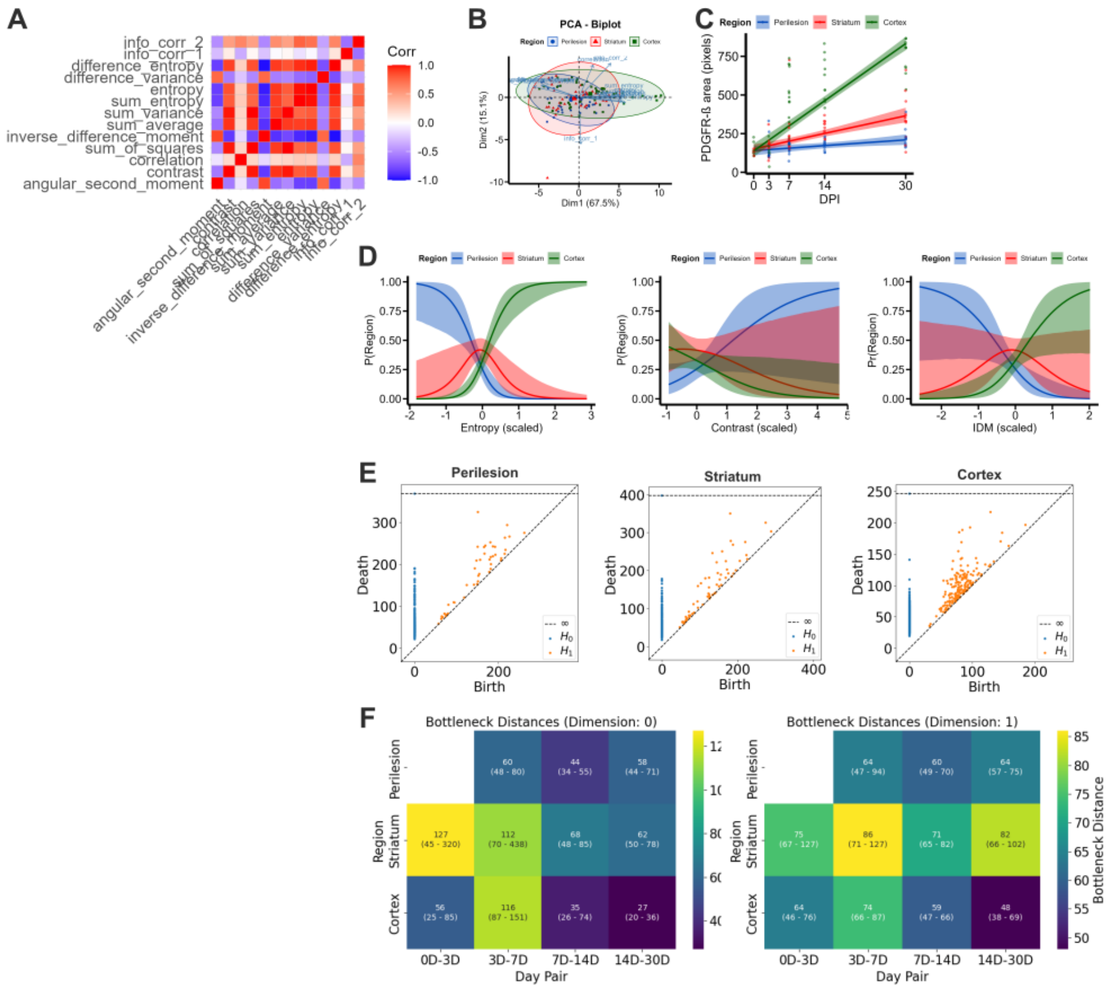

**Supplementary Figure 7. Haralick features and topological data analysis (TDA) of reactive PDGFRβ⁺ cells. A)** Correlation plot of the Haraclick features used for PCA using the ggcorrplot package. See the definition of all the features in Supplementary Method 12. **B)** The Bioplot of Haralick features show the similarities between the variables and its impact on each component. The analysis suggests that all the variables have a similar loading for the components. **C)** Fitted model for the area covered by reactive PDGFRβ⁺ cells (Area ~ DPI * Region, family = student). The posterior estimates are displayed using conditional_effects with lines as mean and 95% CI. Dots are observations (see Suppl. Table 18). **D)** Fitted model for the probability of cell type given specific morphological features (Region ~ entropy + contrast + IDM, family = categorical). The posterior estimates are displayed using conditional_effects with lines as mean and 95% CI. **E)** Representative persistent diagrams per brain region showing the birth (x-axis) and death (y-axis) of topological features. Zero- and 1-dimensional persistent homology from point clouds were calculated using the ripser Python package. Betti curves per brain region are displayed in Figure 5. **F)** Bottleneck distances between pairs of time points calculated from 0 and 1-dimensional homology persistent diagrams. The uncertainty in the mean distances (confidence intervals) was calculated using bootstrapping with 1000 iterations (see Suppl. Table 19c-d). matplotlib was used to produce heatmaps. High-resolution figure: https://osf.io/jxb4z/files/5kumt.

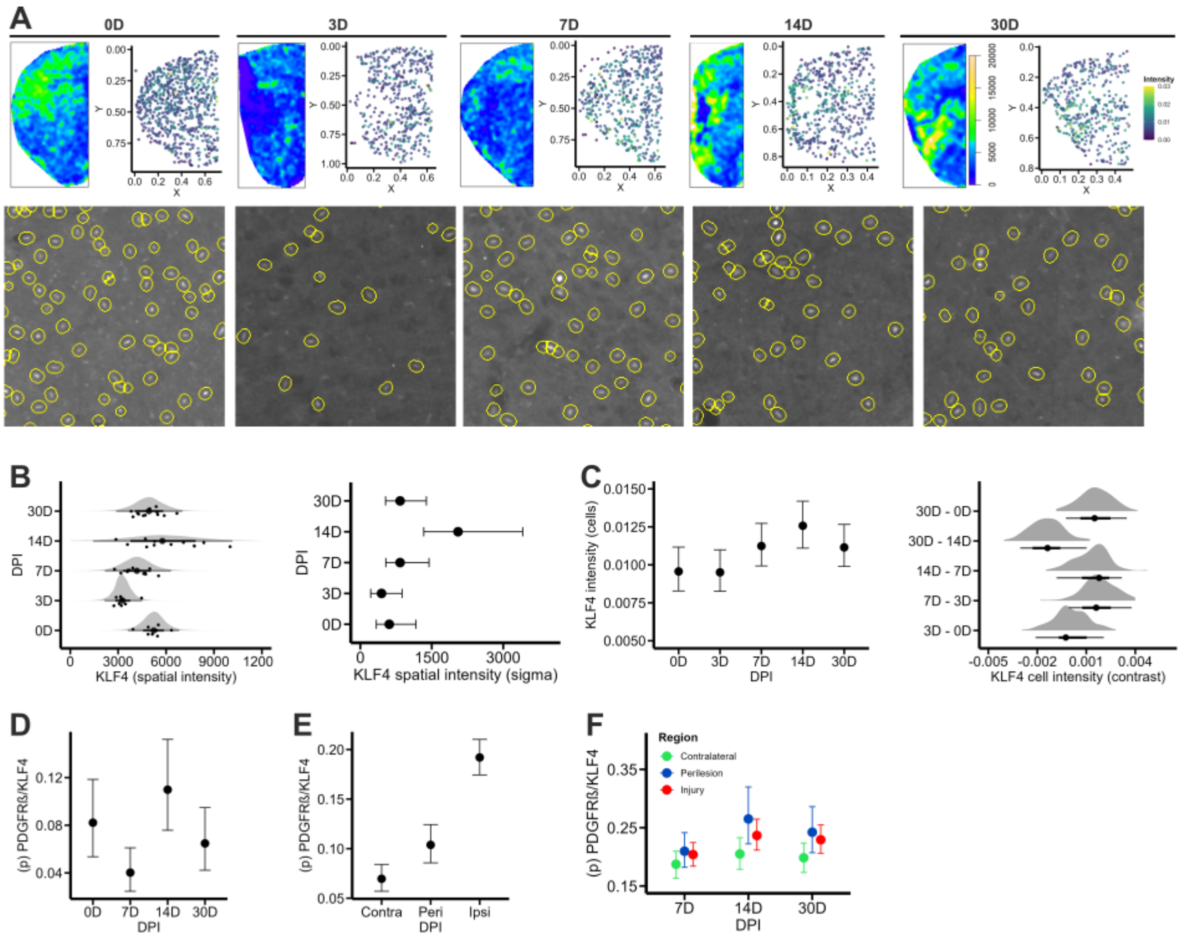

**Supplementary Figure 8. Temporal regulation of KLF4 expression upon ischemic stroke. A)** (upper row) Representative smoothed intensity kernels obtained with *density* (sigma = 0.02) from spatstat at each DPI. Individual cells with KLF4 signal intensity are shown as scatter plots on the right of each density kernel. The dorsolateral brain boundary is located on the left side of the images. (bottom row) Examples of detected individual cells in CellProfiler. The complete image set is available in the OSF repository. **B)** Fitted model for KLF4 intensity ~ 0 + DPI, sigma ~ 0 + DPI, family = student. The posterior probability per DPI (left) is displayed using add_predicted_draws with stat_halfeye and the sigma (heteroskedasticity) estimates with conditional_effects (mean + 95% CI). **C)** Left, fitted model for single cell KLF4 signal intensity (Intensity ~ 0 + DPI * Scaled_CenterX + (1 | MouseID), family = lognormal). The posterior probability with conditional_effects showing mean estimates with 95% CI is displayed as whiskers. Right, the posterior probability of contrasts (emmeans) between time points of interest is displayed using add_predicted_draws with stat_halfeye. **D)** Fitted model for the proportion of PDGFRβ$^+$/KLF4$^+$ cells over total KLF4$^+$ cells in the contralateral hemisphere (Colocalized | trials(Klf4) ~ DPI, family = binomial). This information was used to estimate a ground truth for false-positive colocalization rates given the proximity of non-KLF4-expressing perivascular PDGFRβ$^+$ cells with CD31$^+$ / KLF4$^+$ vasculature. **E)** Fitted model for the proportion of PDGFRβ$^+$ / KLF4$^+$ cells over the total of KLF4$^+$ cells in distinct brain regions (Colocalized | trials(Klf4) ~ Region, family = binomial). **F)** Fitted model for the proportion of PDGFRβ$^+$/KLF4$^+$ cells over the total of PDGFRβ$^+$ cells (Colocalized | trials(Pdgfrb) ~ DPI * Region, family = binomial). **D-F)** The posterior probability with conditional_effects showing mean estimates with 95% CI is displayed as whiskers. High-resolution figure: https://osf.io/jxb4z/files/qgs42.

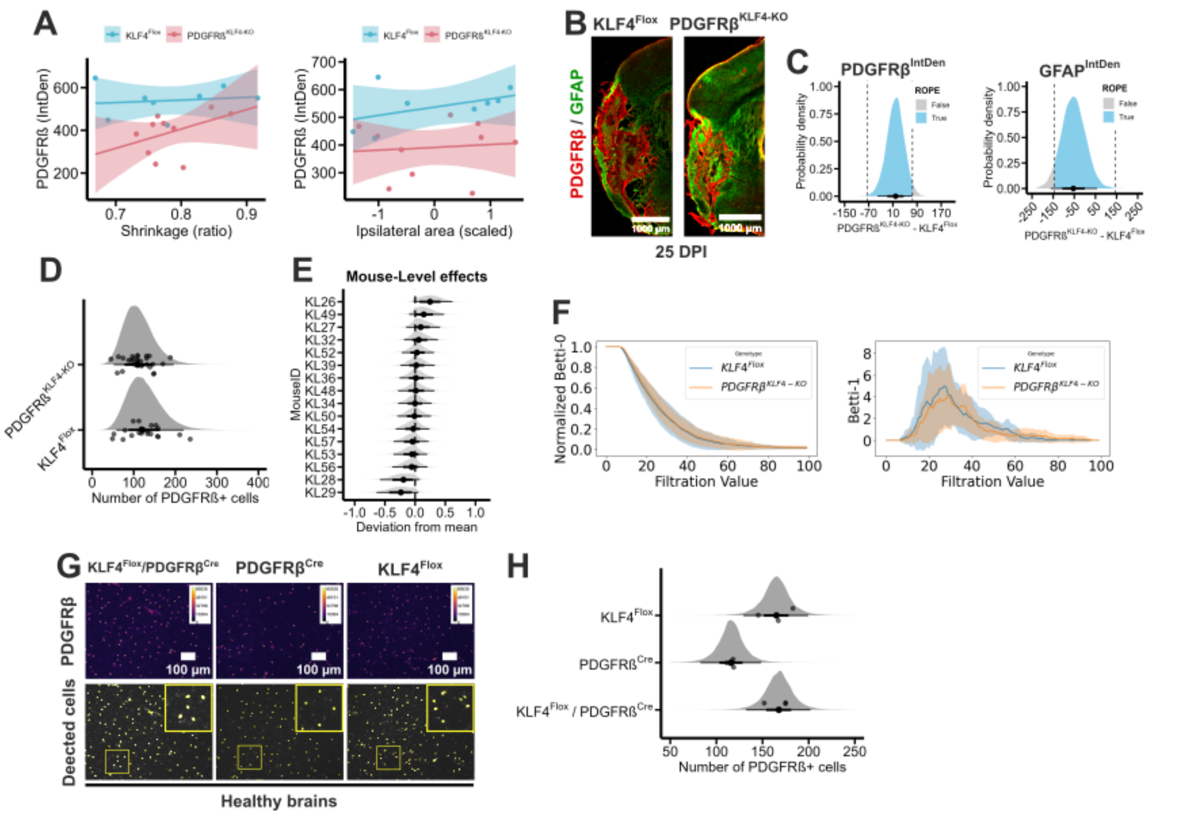

**Supplementary Figure 9. Scar tissue formation in the injured brain of PDGFRβ$^{KLF4-KO}$ mice**. **A)** Fitted models for the PDGFRβ IntDen conditioning on the ipsilateral/contralateral hemispheric ratio (left) (IntDen ~ Genotype * Shrinkage, family = student), and the ipsilateral (scaled) area (IntDen ~ Genotype * Ipsilateral, family = student). PDGFRβ$^{KLF4-KO}$ mice exhibit reduced PDGFRβ expression with increasing hemispheric atrophy (with considerable uncertainty). **B)** Representative brain sections (stitched, 5x magnification) at 25 DPI from PDGFRβ$^{KLF4-KO}$ mice and corresponding controls in which KLF4 depletion was initiated prior to ischemic stroke onset. PDGFRβ (red) and GFAP (green) immunolabeling show a mature glial scar at the chronic phase. **C)** Contrast for PDGFRβ IntDen (left) and GFAP IntDen (right) in the ipsilateral hemisphere of PDGFRβ$^{KLF4-KO}$ mice and corresponding controls for the linear models IntDen ~ Genotype, family = student. The posterior draws (*spread_draws*) are shown with *stat_halfeye()* and ROPE. **D-E)** Fitted multilevel model (Counts ~ Genotype + (1 | MouseID), family = negbinomial) for the number of PDGFRβ$^{+}$ cells conditioning on the genotype (PDGFRβ$^{KLF4-KO}$ and KLF4$^{Flox}$). Three images (ROIs) per mouse brain were analyzed. The posterior estimates are displayed using *add_predicted_draws* with *stat_halfeye()*, showing point estimates and 95% CI. Dots in (D) are observations. Note that E represents the variation within animals. **F)** Mean Betti curves calculated from 0 (left) and 1-dimensional homology (right). Betti curves for 0-dimensional homology are normalized to 1 (1-dimensional graphs are not normalized). PDGFRβ$^{+}$ cells from both genotypes share similar topological features. Line plots are shown using matplotlib. **G)** (upper row) Representative images (10x magnification, 'fire' LUT) of ROIs in the ipsilateral cortex labeled for PDGFRβ mRNA (FISH) in the healthy brain of KLF4$^{Flox}$ / PDGFRβ$^{Cre}$, PDGFRβ$^{Cre}$, and KLF4$^{Flox}$. Yellow represents higher PDGFRβ expression. (bottom row) Corresponding cell detections in CellProfiler showing individual nuclei of PDGFRβ$^{+}$ cells. **H)** Fitted multilevel model (Counts ~ Genotype + (1 | MouseID), family = student) for the number of PDGFRβ$^{+}$ cells conditioning on the genotype (KLF4$^{Flox}$ / PDGFRβ$^{Cre}$, PDGFRβ$^{Cre}$, and KLF4$^{Flox}$). Three images (ROIs) per mouse brain were analyzed. The posterior estimates are displayed using *add_predicted_draws* with *stat_halfeye()*, showing point estimates and 95% CI. Dots are observations. High-resolution figure: https://osf.io/jxb4z/files/875wx.

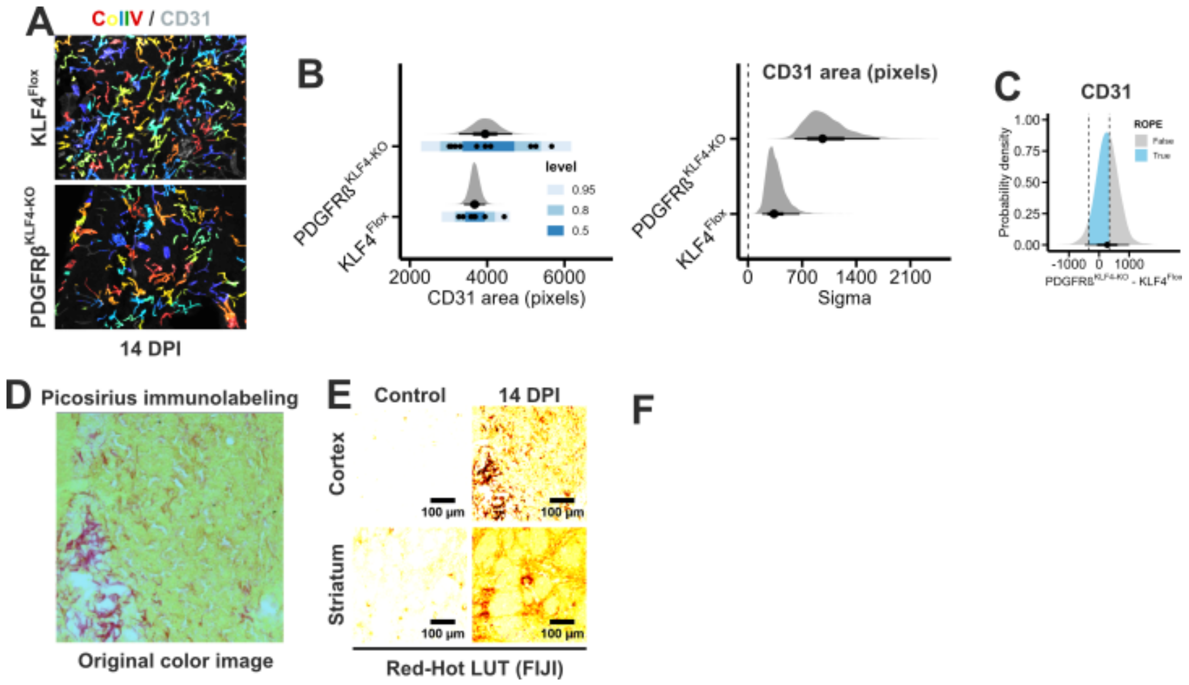

**Supplementary Figure 10. Brain vascular reactivity in PDGFRβ$^{KLF4\text{-}KO}$ mice. A)** Representative images (10x magnification) of Collagen-IV (ColIV) / CD31$^{+}$colocalization analysis in PDGFRβ$^{KLF4\text{-}KO}$ and KLF4$^{Flox}$ mice. Colored regions are ColIV and CD31 is CD31$^{+}$ vasculature. **B)** Fitted model for the labeled CD31 area (Sum_CD31_Area ~ Genotype, sigma ~ Genotype, family = student). We present mean point estimates and their uncertainty using half-eye and stat_interval. Black dots are observations accompanied by prediction intervals (Brewer scale). **C)** Contrast for CD31 labeled area in defined ROIs of PDGFRβ$^{KLF4\text{-}KO}$ mice and corresponding controls for the linear model described in (B). We show posterior draws (*spread_draws*) with *stat_halfeye()* and ROPE. **D)** Representative image from Picosirious staining in the ischemic cortex. Total collagen is visible as a red staining over a yellowish background. **E)** Picosirious-stained images in the striatum and cortex from healthy and ischemic stroke animals (14 DPI). Images are shown showing the Red-Hot LUT in FIJI. **F)** Fitted model for the Picosirious-labeled area (Area ~ Region * Genotype, family = student) for the comparison between PDGFRβ$^{KLF4\text{-}KO}$ and KLF4$^{Flox}$ mice at 14 DPI. We display posterior estimates with conditional_effects (mean + 95% CI). High-resolution figure: https://osf.io/jxb4z/files/cxw6b.
