## Supplementary tables for "Brain PDGFRβ^+^ cells exhibit diverse reactive phenotypes after stroke without requiring KLF4"

**Supplementary Table 1: Hemispheric area following MCAo. A)** Posterior summary with parameter estimates (Est.) and 95% credible intervals (2.5% - 97.5%) for the non-linear (splines) model *Area ~ s(DPI, K=5), sigma ~ DPI, family = student*. Parameters for sigma are in the scale. **B)** Model derivatives (estimate_smooth) showing linear changes from 0-23 DPI and 23-30 DPI.

**(A)** Splines model

| **Parameter** | **Est.** | **2.5 %** | **97.5 %** |
|---|---|---|---|
| Intercept | 11.9 | 11.5 | 12.45 |
| sigma_Intercept | 0.2 | -0.25 | 0.63 |
| sigma_DPI | 0.005 | 0.000 | 0.01 |
| sDPI_1 | 0.079 | -1.91 | 1.99 |
| sds_sDPI_1 | 14.7 | 7.66 | 28.7 |
| Num.Obs. | 49 | | |
| R2 | 0.76 | | |

**(B)** Model derivatives

| **Start** | **End** | **Length** | **Change** | **Slope** | **R2** |
|---|---|---|---|---|---|
| 0.00 | 23.33 | 0.7 | -7.91 | -0.34 | 0.9 |
| 23.33 | 30.00 | 0.2 | 0.17 | 0.03 | 0.9 |

**Supplementary Table 2: PDGFRβ integrated density following MCAo. A)** Posterior summary with parameter estimates (Est.) and 95% credible intervals (2.5% - 97.5%) for the non-linear (splines) model *IntDen ~ s(DPI, K=5), sigma ~ DPI, family = student*. Parameters for sigma are in the scale. **B)** Model derivatives (estimate_smooth) showing linear changes from 0-23 DPI and 23-30 DPI.

**(A)** Splines model

| **Parameter** | **Est.** | **2.5 %** | **97.5 %** |
|---|---|---|---|
| Intercept | 620 | 533 | 702 |
| sigma_Intercept | 5.4 | 5.2 | 5.6 |
| sDPI_1 | 0.003 | -1.9 | 1.9 |
| sigma_sDPI_1 | 11.06 | 1.52 | 24.6 |
| sds_sDPI_1 | 2084 | 920 | 4734 |
| sds_sigma_sDPI_1 | 10.9 | 1.1 | 46.5 |
| s_sigma_sDPI_1[1] | -5.5 | -14.8 | 1.1 |
| s_sigma_sDPI_1[2] | -1.2 | -19.7 | 14.3 |
| s_sigma_sDPI_1[3] | -2.1 | -5.1 | 0.6 |
| Num.Obs. | 49 | | |
| R2 | 0.59 | | |
| R2 Adj. | 0.57 | | |

**(B)** Model derivatives

| **Start** | **End** | **Length** | **Change** | **Slope** | **R2** |
|---|---|---|---|---|---|
| 0 | 23 | 0.66 | 841 | 36.07 | 0.84 |
| 23 | 30 | 0.24 | -29.4 | -4.42 | 0.84 |

**Supplementary Table 3: PDGFRβ integrated density following MCAo (conditioning on hemispheric area). A)** Posterior summary with parameter estimates (Est.) and 95% credible intervals (2.5% - 97.5%) for the non-linear (splines) model *Pdgfrb_IntDen ~ t2(DPI,Tissue_Area), family = Gaussian*.

| Parameter | Est. | 2.5 % | 97.5 % |
|---|---|---|---|
| b_Intercept | 620 | 512 | 745 |
| bs_t2DPITissue_Area_1 | -206 | -471 | 118 |
| bs_t2DPITissue_Area_2 | -119 | -460 | 133 |
| bs_t2DPITissue_Area_3 | 85 | -391 | 463 |
| sds_t2DPITissue_Area_1 | 563 | 18 | 1763 |
| sds_t2DPITissue_Area_2 | 621 | 17 | 2382 |
| sds_t2DPITissue_Area_3 | 876 | 34 | 2685 |
| sigma | 268 | 215 | 336 |
| Num.Obs. | 49 | | |
| R2 | 0.63 | | |
| R2 Adj. | 0.56 | | |

**Supplementary Table 4: GFAP integrated density following MCAo. A)** Posterior summary with parameter estimates (Est.) and 95% credible intervals (2.5% - 97.5%) for the non-linear (splines) model *IntDen ~ s(DPI, K=5), sigma ~ DPI, family = student*. Parameters for sigma are shown in the scale. **B)** Model derivatives (estimate_smooth) showing linear changes from 0-16 DPI and 16-30 DPI.

**(A)** Splines model

| Parameter | Est. | 2.5 % | 97.5 % |
|---|---|---|---|
| b_Intercept | 490 | 381 | 615 |
| b_sigma_Intercept | 5.2 | 5.08 | 5.5 |
| bs_sDPI_1 | 0.003 | -1.9 | 1.9 |
| bs_sigma_sDPI_1 | 26.4 | 6.1 | 49.6 |
| sds_sDPI_1 | 1698 | 817 | 3576 |
| sds_sigma_sDPI_1 | 22.7 | 4.9 | 85.8 |
| s_sigma_sDPI_1[1] | -14.7 | -25.9 | -4.2 |
| s_sigma_sDPI_1[2] | 5.1 | -19.2 | 33.3 |
| s_sigma_sDPI_1[3] | -2.9 | -8.1 | 1.7 |
| Num.Obs. | 49 | | |
| R2 | 0.51 | | |
| R2 Adj. | 0.18 | | |

**(B)** Model derivatives

| Start | End | Length | Change | Slope | R2 |
|---|---|---|---|---|---|
| 0 | 16 | 0.46 | 835.13 | 50.11 | 0.24 |
| 16 | 30 | 0.44 | -480.65 | -36.05 | 0.24 |

**Supplementary Table 5: GFAP integrated density following MCAo (conditioning on hemispheric area).** Posterior summary with parameter estimates (Est.) and 95% credible intervals (2.5% - 97.5%) for the non-linear (splines) model *Gfap_IntDen ~ t2(DPI,Tissue_Area), family = Gaussian*.

| Parameters | Est. | 2.5 % | 97.5 % |
|---|---|---|---|
| b_Intercept | 509 | 425 | 605 |
| bs_t2DPITissue_Area_1 | 229 | 17 | 451 |
| bs_t2DPITissue_Area_2 | -325 | -532 | -126 |
| bs_t2DPITissue_Area_3 | -369 | -645 | -108 |
| sds_t2DPITissue_Area_1 | 360 | 17 | 1078 |
| sds_t2DPITissue_Area_2 | 350 | 11 | 1288 |
| sds_t2DPITissue_Area_3 | 411 | 14 | 1405 |
| sigma | 242 | 196 | 301 |
| Num.Obs. | 49 | | |
| R2 | 0.45 | | |
| R2 Adj. | 0.37 | | |

**Supplementary Table 6: PDGFRβ$^{high}$ likelihood following MCAo.** Posterior summary with parameter estimates (Est.) and 95% credible intervals (2.5% - 97.5%) for the binomial model *Pdgfrb_high | trials (Pdgfrb_Total) ~ s(DPI, k =5), family = binomial*.

| Parameters | Est. | 2.5 % | 97.5 % |
|---|---|---|---|
| b_Intercept | 1.130 | 1.114 | 1.146 |
| bs_sDPI_1 | 25.6 | 23.9 | 27.3 |
| sds_sDPI_1 | 9.5 | 5.1 | 18.4 |
| Num.Obs. | 49 | | |
| R2 | 0.97 | | |

**Supplementary Table 7: GFAP convex hull following MCAo.** Posterior summary with parameter estimates (Est.) and 95% credible intervals (2.5% - 97.5%) for the linear model *Healthy_Ratio ~ DPI, family = student*.

| Parameter | Est. | 2.5 % | 97.5 % |
|---|---|---|---|
| b_Intercept | 61.1 | 56.4 | 65.9 |
| b_DPI | -0.100 | -0.38 | 0.17 |
| sigma | 8.20 | 6.33 | 10.47 |
| Num.Obs. | 38 | | |
| R2 | 0.02 | | |

**Supplementary Table 8: PDGFRβ counts in tesselated GFAP regions. A)** Posterior summary with parameter estimates (Est.) and 95% credible intervals (2.5% - 97.5%) for the model *Pdgfrb ~ Region x Gfap, family = hurdle_lognormal()*. **B-D)** Posterior summary for models by region *Pdgfrb ~ DPI x Gfap, family = hurdle_lognormal()*.

**(A)** Model for all regions

| **Paremeters** | **Est.** | **2.5 %** | **97.5 %** |
|---|---|---|---|
| b_Intercept | 3.7 | 3.5 | 4.0 |
| b_Striatum | 0.4 | 0.1 | 0.8 |
| b_Cortex | 1.06 | 0.66 | 1.45 |
| b_Gfap_High | -0.46 | -0.83 | -0.09 |
| b_Striatum × Gfap_High | 0.45 | -0.06 | 0.97 |
| b_RegionCortex × Gfap_High | -0.64 | -1.19 | -0.06 |
| sigma | 1.20 | 1.10 | 1.31 |
| Num.Obs. | 262 | | |
| R2 | 0.43 | | |

**(B)** Model for perilesion

| **Parameters** | **Est.** | **2.5 %** | **97.5 %** |
|---|---|---|---|
| b_Intercept | 3.53 | 3.08 | 3.99 |
| b_DPI7D | 0.29 | -0.35 | 0.93 |
| b_DPI14D | 0.51 | -0.09 | 1.12 |
| b_DPI30D | 0.09 | -0.53 | 0.73 |
| b_Gfap_High | -0.40 | -1.05 | 0.25 |
| b_DPI7D × Gfap_High | 0.48 | -0.41 | 1.37 |
| b_DPI14D × Gfap_High | 0.07 | -0.79 | 0.93 |
| b_DPI30D × Gfap_High | -0.75 | -1.62 | 0.13 |
| sigma | 0.99 | 0.84 | 1.17 |
| Num.Obs. | 88 | | |
| R2 | 0.391 | | |

**(C)** Model for striatum

| Parameters | Est. | 2.5 % | 97.5 % |
|---|---|---|---|
| b_Intercept | 3.38 | 2.95 | 3.81 |
| b_DPI7D | 0.88 | 0.30 | 1.44 |
| b_DPI14D | 1.19 | 0.62 | 1.75 |
| b_DPI30D | 1.27 | 0.70 | 1.85 |
| b_Gfap_High | -1.47 | -2.08 | -0.87 |
| b_DPI7D × Gfap_High | 1.47 | 0.67 | 2.29 |
| b_DPI14D × Gfap_High | 2.05 | 1.25 | 2.86 |
| b_DPI30D × Gfap_High | 2.01 | 1.21 | 2.82 |
| sigma | 0.92 | 0.79 | 1.09 |
| Num.Obs. | 88 | | |
| R2 | 0.241 | | |

**(D)** Model for cortex

| Parameters | Est. | 2.5 % | 97.5 % |
|---|---|---|---|
| b_Intercept | -12.51 | -49.35 | 0.74 |
| b_DPI7D | 16.45 | 3.20 | 53.36 |
| b_DPI14D | 17.98 | 4.72 | 54.96 |
| b_DPI30D | 18.48 | 5.21 | 55.41 |
| b_Gfap_High | -24.25 | -76.36 | -5.43 |
| b_DPI7D × Gfap_High | 21.41 | 2.51 | 73.70 |
| b_DPI14D × Gfap_High | 22.79 | 3.95 | 74.85 |
| b_DPI30D × Gfap_High | 23.72 | 4.92 | 75.88 |
| shape | 0.76 | 0.54 | 1.04 |
| Num.Obs. | 86 | | |
| R2 | 0.575 | | |

**Supplementary Table 9: PDGFRβ$^{high}$ likelihood in striatal-only lesions**. Posterior summary with parameter estimates (Est.) and 95% credible intervals (2.5% - 97.5%) for the binomial model *Pdgfrb_high | trials(Pdgfrb_Total) ~ DPI, family = binomial*.

| **Parameters** | **Est.** | **2.5 %** | **97.5 %** |
|---|---|---|---|
| b_Intercept | -0.65 | -0.73 | -0.57 |
| b_DPI | 0.02 | 0.02 | 0.02 |
| Num.Obs. | 14 | | |
| R2 | 0.640 | | |

**Supplementary Table 10: Parenchymal PDGFRβ likelihood following MCAo. A-C)** Posterior summary with parameter estimates (Est.) and 95% credible intervals (2.5% - 97.5%) for the binomial models *Pdgfrb_Parenchymal | trials (Pdgfrb_Total) ~ DPI, family = binomial* for perilesion, sriatum and cortex, respectively.

**(A)** Model for perilesion

| **Parameters** | **Est.** | **2.5 %** | **97.5 %** |
|---|---|---|---|
| b_Intercept | -2.84 | -2.98 | -2.70 |
| bs_sDPI2_1 | 0.059 | -3.93 | 4.71 |
| sds_sDPI2_1 | 3.22 | 1.13 | 7.70 |
| Num.Obs. | 43 | | |
| R2 | 0.207 | | |

**(B)** Model for striatum

| **Parameters** | **Est.** | **2.5 %** | **97.5 %** |
|---|---|---|---|
| b_Intercept | -1.37 | -1.44 | -1.30 |
| bs_sDPI2_1 | 4.19 | 1.08 | 7.64 |
| sds_sDPI2_1 | 2.92 | 0.96 | 7.07 |
| Num.Obs. | 43 | | |
| R2 | 0.69 | | |

**(C)** Model for cortex

| **Parameters** | **Est.** | **2.5 %** | **97.5 %** |
|---|---|---|---|
| b_Intercept | -0.93 | -0.97 | -0.89 |
| bs_sDPI2_1 | 17.37 | 15.07 | 19.75 |
| sds_sDPI2_1 | 8.71 | 4.42 | 18.52 |
| Num.Obs. | 40 | | |
| R2 | 0.83 | | |

**Supplementary Table 11: FACS-sorted PDGFRβ⁺ cells derived from contralateral and ipsilateral hemispheres**. **A-B)** Posterior summary with parameter estimates (Est.) and 95% credible intervals (2.5% - 97.5%) for the linear models *Cells ~ Hemisphere, family = student* estimating the percentage of total PDGFRβ+ cells **(A)** and dead PDGFRβ+ cells **(B)**.

**(A)** Model for the number of PDGFR$\beta^+$ cells

| **Parameter** | **Est.** | **2.5 %** | **97.5 %** |
|---|---|---|---|
| b_Intercept | 0.062 | 0.034 | 0.091 |
| b_HemisphereIpsi | 0.074 | 0.033 | 0.114 |
| sigma | 0.030 | 0.017 | 0.054 |
| Num.Obs. | 10 | | |
| R2 | 0.72 | | |

**(B)** Model for the number of PDGFR$\beta^+$ dead cells

| **Parameter** | **Est.** | **2.5 %** | **97.5 %** |
|---|---|---|---|
| b_Intercept | 0.106 | 0.004 | 0.209 |
| b_HemisphereIpsi | 0.137 | -0.006 | 0.299 |
| sigma | 0.108 | 0.044 | 0.208 |
| Num.Obs. | 10 | | |
| R2 | 0.33 | | |

**Supplementary Table 12: Parenchymal PDGFRβ likelihood in striatal-restricted lesions**. Posterior summary with parameter estimates (Est.) and 95% credible intervals (2.5% - 97.5%) for the binomial model *Pdgfrb_Parenchymal | trials (Pdgfrb_Total) ~ DPI, family = binomial* in striatal-only lesions.

| Parameters | Est. | 2.5 % | 97.5 % |
|---|---|---|---|
| b_Intercept | -1.58 | -1.79 | -1.38 |
| b_DPI30D | 0.38 | 0.15 | 0.60 |
| b_Cortex | 0.52 | 0.30 | 0.74 |
| b_DPI30D × Cortex | -0.68 | -0.93 | -0.43 |
| Num.Obs. | 38 | | |
| R2 | 0.653 | | |

**Supplementary Table 13: Number of PDGFRβ$^+$ cells in striatal-only and cortico-striatal lesions**. Posterior summary with parameter estimates (Est.) and 95% credible intervals (2.5% - 97.5%) for the model *Pdgfrb_Total ~ DPI x Lesion, sigma ~ DPI x Lesion, family* = *student* in striatal-only lesions.

| Parameters | Est. | 2.5 % | 97.5 % |
|---|---|---|---|
| b_Intercept | 159 | 25 | 269 |
| b_sigma_Intercept | 4.40 | 3.21 | 5.66 |
| b_DPI30D | 64.4 | -61.2 | 211.6 |
| b_Cortex | 59.7 | -82.4 | 229.5 |
| b_DPI30D × Cortex | -20.3 | -228.2 | 180.2 |
| b_sigma_DPI30D | -0.07 | -1.46 | 1.11 |
| b_sigma_Cortex | 0.37 | -1.16 | 1.51 |
| b_sigma_DPI30D × Cortex | 0.50 | -0.90 | 2.17 |
| Num.Obs. | 38 | | |
| R2 | 0.065 | | |

**Supplementary Table 14: Number of parenchymal PDGFRβ⁺ cells in sham and MCAo mice**. Posterior summary with parameter estimates (Est.) and 95% credible intervals (2.5% - 97.5%) for the model *Parenchymal ~ DPI x Condition, family = student* in sham and MCAo mice.

| Parameters | Est. | 2.5 % | 97.5 % |
|---|---|---|---|
| b_Intercept | 1.006 | 0.44 | 1.52 |
| b_DPI7D | -0.001 | -0.75 | 0.73 |
| b_DPI14D | 0.29 | -0.40 | 0.98 |
| b_DPI30D | 0.04 | -0.81 | 0.85 |
| b_MCAo | 0.73 | 0.16 | 1.34 |
| b_DPI7D × MCAo | 0.11 | -0.68 | 0.91 |
| b_DPI14D × MCAo | -1.02 | -1.83 | -0.20 |
| b_DPI30D × MCAo | 0.10 | -0.74 | 1.02 |
| Num.Obs. | 60 | | |
| R2 | 0.154 | | |

**Supplementary Table 15: PDGFRβ+ cells proliferation in the ischemic hemispheric following MCAo. A)** Posterior summary with parameter estimates (Est.) and 95% credible intervals (2.5% - 97.5%) for the non-linear (splines) model *Percentage_Pdgfrb ~ s(DPI_Cont, k = 5), family = lognormal*. **B)** Model derivatives (estimate_smooth) showing linear changes from 0-3 DPI and 3-30 DPI.

**(A)** Splines model

| **Parameters** | **Est.** | **2.5 %** | **97.5 %** |
|---|---|---|---|
| b_Intercept | 0.74 | 0.59 | 0.88 |
| bs_sDPI_Cont_1 | 65.7 | 48.6 | 80.9 |
| sds_sDPI_Cont_1 | 33.7 | 16.6 | 67.4 |
| sigma | 0.52 | 0.42 | 0.66 |
| Num.Obs. | 50 | | |
| R2 | 0.528 | | |

**(B)** Model derivatives

| **Start** | **End** | **Length** | **Change** | **Slope** | **R2** |
|---|---|---|---|---|---|
| 0 | 3 | 0.08 | 6.87 | 2.06 | 0.24 |
| 3 | 30 | 0.82 | -6.10 | -0.23 | 0.24 |

**Supplementary Table 16: PDGFRβ⁺ cells proliferation in defined ROIs following MCAo. A)** Posterior summary with parameter estimates (Est.) and 95% credible intervals (2.5% - 97.5%) for the model *Cells ~ DPI x Region, family = negbinomial* showing the total amount of colocalized cells. **B)** The same estimates for the model *Vascular | trials (Total) ~ DPI x Region, family = binomial* showing the proportion of vascular PDGFRβ$^+$ cells relative to the total number of PDGFRβ$^+$ cells.

**(A)** Model for total PDGFRβ/Ki67$^+$ cells

| **Parameters** | **Est.** | **2.5 %** | **97.5 %** |
|---|---|---|---|
| b_Intercept | -1.55 | -3.46 | -0.01 |
| b_DPI7D | 0.37 | -1.56 | 2.51 |
| b_RegionStr | 2.22 | 0.43 | 4.31 |
| b_RegionCtx | 2.75 | 0.99 | 4.82 |
| b_DPI7D × RegionStr | -0.95 | -3.45 | 1.41 |
| b_DPI7D × RegionCtx | 0.20 | -2.19 | 2.48 |
| shape | 1.01 | 0.45 | 2.04 |
| Num.Obs. | 59 | | |
| R2 | 0.391 | | |

**(B)** Model for vascular PDGFRβ/Ki67$^+$ cells

| **Parameters** | **Est.** | **2.5 %** | **97.5 %** |
|---|---|---|---|
| b_Intercept | 95.8 | 3.3 | 437.3 |
| b_DPI7D | -57.3 | -346.1 | 72.3 |
| b_RegionStr | -92.8 | -434.7 | -0.14 |
| b_RegionCtx | -94.4 | -436.3 | -1.7 |
| b_DPI7D × RegionStr | 56.2 | -73.8 | 343.3 |
| b_DPI7D × RegionCtx | 56.06 | -73.68 | 345 |
| Num.Obs. | 31 | | |
| R2 | 0.813 | | |

**Supplementary Table 17: Haralick features of PDGFRβ$^{+}$ cells**. Posterior summary with parameter estimates (Est.) and 95% credible intervals (2.5% - 97.5%) for the logistic model *Region ~ entropy + contrast + IDM, family = categorical*.

| **Parameters** | **Est.** | **2.5 %** | **97.5 %** |
|---|---|---|---|
| b_muStriatum_Intercept | 0.49 | -0.011 | 1.04 |
| b_muCortex_Intercept | 0.25 | -0.29 | 0.82 |
| b_muStriatum_entropy | 2.54 | 0.54 | 4.68 |
| b_muStriatum_contrast | -0.79 | -1.92 | 0.27 |
| b_muStriatum_IDM | 1.43 | -0.13 | 3.09 |
| b_muCortex_entropy | 5.26 | 3.21 | 7.55 |
| b_muCortex_contrast | -1.12 | -2.22 | -0.03 |
| b_muCortex_IDM | 2.92 | 1.13 | 4.83 |
| Num.Obs. | 146 | | |

**Supplementary Table 18: PDGFRβ stained area in defined ROIs**. Posterior summary with parameter estimates (Est.) and 95% credible intervals (2.5% - 97.5%) for the binomial model *Area ~ DPI_cont x Region, family = student*.

| Parameters | Est. | 2.5 % | 97.5 % |
|---|---|---|---|
| b_Intercept | 143 | 123 | 164 |
| b_DPI | 10.8 | 9.6 | 12.0 |
| b_Striatum | -0.3 | -36.6 | 38.9 |
| b_Cortex | -9.3 | -40.3 | 22.4 |
| b_DPI × Striatum | 14.6 | 12.6 | 16.6 |
| b_DPI × Cortex | 4.5 | 2.4 | 6.4 |
| sigma | 58.1 | 43.4 | 76.8 |
| Num.Obs. | 143 | | |
| R2 | 0.698 | | |

**Supplementary Table 19: Topological assessment of PDGFRβ⁺ cells following injury:** Wasserstein (**A-B)** and Bottleneck **(C-D)** distances for 0- and 1-dimension homology. We boostraped the estimates (1000 replications) to show the median Betti curve distance with 95% credible intervals (2.5% - 97.5%).

**(A)** Wasserstein distance - 0D homology

| Day1 | Day2 | Region | Median | 2.5 % | 97.5 % |
|---|---|---|---|---|---|
| 0D | 3D | Ctx | 211 | 150 | 323 |
| 3D | 7D | Ctx | 603 | 507 | 669 |
| 7D | 14D | Ctx | 208 | 146 | 280 |
| 14D | 30D | Ctx | 156 | 119 | 225 |
| 0D | 3D | Str | 385 | 168 | 548 |
| 3D | 7D | Str | 348 | 222 | 784 |
| 7D | 14D | Str | 227 | 184 | 315 |
| 14D | 30D | Str | 243 | 177 | 318 |
| 3D | 7D | Peri | 230 | 172 | 308 |
| 7D | 14D | Peri | 182 | 150 | 229 |
| 14D | 30D | Peri | 239 | 189 | 304 |

**(B)** Wasserstein distance - 1D homology

| Day1 | Day2 | Region | Median | 2.5 % | 97.5 % |
|---|---|---|---|---|---|
| 0D | 3D | Ctx | 126 | 111 | 168 |
| 3D | 7D | Ctx | 216 | 194 | 232 |
| 7D | 14D | Ctx | 135 | 111 | 166 |
| 14D | 30D | Ctx | 122 | 94 | 153 |
| 0D | 3D | Str | 134 | 115 | 162 |
| 3D | 7D | Str | 158 | 129 | 194 |
| 7D | 14D | Str | 146 | 137 | 171 |
| 14D | 30D | Str | 166 | 143 | 183 |
| 3D | 7D | Peri | 127 | 111 | 159 |

| | | | | | |
|---|---|---|---|---|---|
| 7D | 14D | Peri | 121 | 106 | 132 |
| 14D | 30D | Peri | 138 | 122 | 158 |

**(C)** Bottleneck distance - 0D homology

| **Day1** | **Day2** | **Region** | **Median** | **2.5 %** | **97.5 %** |
|---|---|---|---|---|---|
| 0D | 3D | Str | 127 | 45 | 320 |
| 3D | 7D | Str | 112 | 70 | 438 |
| 7D | 14D | Str | 68 | 48 | 85 |
| 14D | 30D | Str | 62 | 50 | 78 |
| 0D | 3D | Ctx | 56 | 25 | 85 |
| 3D | 7D | Ctx | 116 | 87 | 151 |
| 7D | 14D | Ctx | 35 | 26 | 74 |
| 14D | 30D | Ctx | 27 | 20 | 36 |
| 3D | 7D | Peri | 60 | 48 | 80 |
| 7D | 14D | Peri | 44 | 34 | 55 |
| 14D | 30D | Peri | 58 | 44 | 71 |

**(D)** Bottleneck distance - 1D homology

| **Day1** | **Day2** | **Region** | **Median** | **2.5 %** | **97.5 %** |
|---|---|---|---|---|---|
| 0D | 3D | Str | 75 | 67 | 127 |
| 3D | 7D | Str | 86 | 71 | 127 |
| 7D | 14D | Str | 71 | 65 | 82 |
| 14D | 30D | Str | 82 | 66 | 102 |
| 0D | 3D | Ctx | 64 | 46 | 76 |
| 3D | 7D | Ctx | 74 | 66 | 87 |
| 7D | 14D | Ctx | 59 | 47 | 66 |
| 14D | 30D | Ctx | 48 | 38 | 69 |
| 3D | 7D | Peri | 64 | 47 | 94 |
| 7D | 14D | Peri | 60 | 49 | 70 |
| 14D | 30D | Peri | 64 | 57 | 75 |

**Supplementary Table 20: Spatial intensity of KLF4+ nuclei following injury.** Posterior summary with parameter estimates (Est.) and 95% credible intervals (2.5% - 97.5%) for the model *Intensity ~ 0 + DPI, sigma ~ 0 + DPI, family = student*. Note that the sigma coefficients are in the log scale.

| Parameters | Est. | 2.5 % | 97.5 % |
|---|---|---|---|
| b_DPI_0D | 5214 | 4756 | 5667 |
| b_DPI_3D | 3281 | 2966 | 3658 |
| b_DPI_7D | 4177 | 3652 | 4729 |
| b_DPI_14D | 5787 | 4652 | 6835 |
| b_DPI_30D | 4928 | 4427 | 5439 |
| b_sigma_DPI_0D | 6.41 | 5.81 | 7.06 |
| b_sigma_DPI_3D | 6.09 | 5.38 | 6.77 |
| b_sigma_DPI_7D | 6.74 | 6.28 | 7.27 |
| b_sigma_DPI_14D | 7.63 | 7.19 | 8.13 |
| b_sigma_DPI_30D | 6.73 | 6.27 | 7.23 |
| Num.Obs. | 53 | | |
| R2 | 0.35 | | |

**Supplementary Table 21: Staining intensity of KLF4$^+$ nuclei following injury.** Population-level posterior summary with parameter estimates (Est.) and 95% credible intervals (2.5% - 97.5%) for the multilevel model *Intensity ~ 0 + DPI + (1 | MouseID), family = hurdle_lognormal*. Note that the group-level (mouse-level) coefficients are shown in the respective QN.

| Parameter | Est. | 2.5 % | 97.5% |
|---|---|---|---|
| 0 DPI | -4.79 | -4.94 | -4.64 |
| 3 DPI | -4.8 | -4.94 | -4.66 |
| 7 DPI | -4.63 | -4.76 | -4.51 |
| 14 DPI | -4.52 | -4.65 | -4.4 |
| 30 DPI | -4.64 | -4.76 | -4.51 |

**Supplementary Table 22: Staining intensity of KLF4⁺ nuclei relative to the x-coordinates.** Population-level posterior summary with parameter estimates (Est.) and 95% credible intervals (2.5% - 97.5%) for the multilevel model *Intensity ~ 0 + DPI x Scaled_CenterX* + *(1 | MouseID), family* = *lognormal*. Note that the group-level (mouse-level) coefficients are shown in the respective QN.

| Parameter | Est. | 2.5 % | 97.5 % |
|---|---|---|---|
| 0 DPI | -4.8 | -4.95 | -4.65 |
| 3 DPI | -4.84 | -4.99 | -4.7 |
| 7 DPI | -4.63 | -4.75 | -4.5 |
| 14 DPI | -4.42 | -4.54 | -4.3 |
| 30 DPI | -4.59 | -4.71 | -4.46 |
| Scaled_x | 0.06 | -0.01 | 0.14 |
| 3 DPI:Scaled_x | 0.06 | -0.05 | 0.16 |
| 7 DPI:Scaled_x | -0.1 | -0.21 | 0.01 |
| 14 DPI:Scaled_x | -0.54 | -0.67 | -0.41 |
| 30 DPI:Scaled_x | -0.31 | -0.47 | -0.15 |

**Supplementary Table 23: KLF4/PDGFRβ co-localization in defined ROIs.** Posterior summary with parameter estimates (Est.) and 95% credible intervals (2.5% - 97.5%) for the model *Coloc | trials(Klf4) ~ DPI x Region, family = binomial*.

| Parameter | Est. | 2.5 % | 97.5 % |
|---|---|---|---|
| b_Intercept | -2.27 | -2.41 | -2.13 |
| b_14D | 0.06 | 0.01 | 0.16 |
| b_30D | 0.12 | 0.02 | 0.28 |
| b_Perilesion | 0.07 | 0.01 | 0.17 |
| b_Injury | 0.52 | 0.37 | 0.66 |
| b_14D × Perilesion | 0.11 | 0.01 | 0.27 |
| b_30D × Perilesion | 0.07 | 0.01 | 0.20 |
| b_14D × Injury | 0.05 | 0.008 | 0.14 |
| b_30D × Injury | 0.34 | 0.15 | 0.53 |
| Num.Obs. | 64 | | |
| R2 | 0.595 | | |

**Supplementary Table 24: Spatial intensity of KLF4$^{+}$ nuclei in PDGFRβ$^{KLF4\text{-}KO}$ mice following injury.** Posterior summary with parameter estimates (Est.) and 95% credible intervals (2.5% - 97.5%) for the model *Intensity ~ Genotype, sigma ~ Genotype, family = student*.

| Parameter | Est. | 2.5 % | 97.5 % |
|---|---|---|---|
| b_Intercept | 9478 | 8435 | 10475 |
| b_sigma_Intercept | 7.2 | 6.7 | 7.9 |
| b_GenotypeKO | 258 | -805 | 1392 |
| b_sigma_GenotypeKO | -0.5 | -1.3 | 0.2 |
| Num.Obs. | 19 | | |
| R2 | 0.03 | | |

**Supplementary Table 25: Hemispheric ratio in PDGFRβ$^{KLF4\text{-}KO}$ mice.** Posterior summary with parameter estimates (Est.) and 95% credible intervals (2.5% - 97.5%) for the model *Shrinkage ~ Genotype, family = student*.

| Parameters | Est. | 2.5 % | 97.5 % |
|---|---|---|---|
| b_Intercept | 0.76 | 0.71 | 0.81 |
| b_GenotypeKO | 0.02 | -0.04 | 0.09 |
| sigma | 0.06 | 0.04 | 0.10 |
| Num.Obs. | 18 | | |
| R2 | 0.04 | | |

**Supplementary Table 26: GFAP integrated density in PDGFRβ$^{KLF4\text{-}KO}$ mice.** Posterior summary with parameter estimates (Est.) and 95% credible intervals (2.5% - 97.5%) for the model *Gfap ~ Genotype, family = student*.

| **Parameters** | **Est.** | **2.5 %** | **97.5 %** |
|---|---|---|---|
| b_Intercept | 763 | 681 | 844 |
| b_GenotypeKO | 122 | 12 | 233 |
| sigma | 107 | 73 | 158 |
| Num.Obs. | 18 | | |
| R2 | 0.26 | | |

**Supplementary Table 27: PDGFRβ integrated density in PDGFRβ$^{KLF4\text{-}KO}$ mice.** Posterior summary with parameter estimates (Est.) and 95% credible intervals (2.5% - 97.5%) for the model *Pdgfrb ~ Genotype, family = student*.

| Parameters | Est. | 2.5 % | 97.5 % |
|---|---|---|---|
| b_Intercept | 539 | 473 | 601 |
| b_GenotypeKO | -148 | -239 | -57 |
| sigma | 89 | 60 | 130 |
| Num.Obs. | 18 | | |
| R2 | 0.43 | | |

**Supplementary Table 28: PDGFRβ integrated density in PDGFRβ$^{KLF4\text{-}KO}$ mice with additional predictors.** Posterior summary with parameter estimates (Est.) and 95% credible intervals (2.5% - 97.5%) for the models a) *Pdgfrb ~ Genotype x Shrinkage, family = student* and b) *Pdgfrb ~ Genotype x Ipsilateral_Scaled, family student*.

**(A)** Model conditioning on brain shrinkage

| **Parameters** | **Est.** | **2.5 %** | **97.5 %** |
|---|---|---|---|
| b_Intercept | 447 | -250 | 1134 |
| b_GenotypeKO | -766 | -2060 | 567 |
| b_Shrinkage | 117 | -767 | 998 |
| b_GenotypeKO × Shrinkage | 787 | -912 | 2418 |
| sigma | 89 | 58 | 135 |
| Num.Obs. | 18 | | |
| R2 | 0.49 | | |

**(B)** Model conditioning on ipsilateral area

| **Parameter** | **Est.** | **2.5 %** | **97.5 %** |
|---|---|---|---|
| b_Intercept | 537 | 470 | 604 |
| b_GenotypeKO | -145 | -239 | -52 |
| b_Ipsilateral_Scaled | 29 | -35 | 92 |
| b_GenotypeKO × Ipsilateral_Scaled | -19 | -113 | 77 |
| sigma | 92 | 60 | 137 |
| Num.Obs. | 18 | | |
| R2 | 0.47 | | |

**Supplementary Table 29: PDGFRβ and GFAP integrated density in PDGFRβ$^{KLF4-KO}$ mice (intraperitoneal injection prior to ischemia).** Posterior summary with parameter estimates (Est.) and 95% credible intervals (2.5% - 97.5%) for the models a) *Pdgfrb ~ Genotype, family = student* and b) *Gfap ~ Genotype, family student*.

**(A)** Model for PDGFRβ IntDen

| **Parameters** | **Est.** | **2.5 %** | **97.5 %** |
|---|---|---|---|
| b_Intercept | 72.3 | 27.8 | 122.6 |
| b_GenotypeKO | 18.3 | -41.0 | 74.4 |
| sigma | 75.1 | 48.0 | 103.9 |
| Num.Obs. | 34 | | |
| R2 | 0.019 | | |

**(B)** Model for GFAP IntDen

| **Parameters** | **Est.** | **2.5 %** | **97.5 %** |
|---|---|---|---|
| b_Intercept | 446.6 | 366.9 | 526.0 |
| b_GenotypeKO | -54.0 | -162.4 | 58.1 |
| sigma | 144.3 | 94.1 | 196.9 |
| Num.Obs. | 34 | | |
| R2 | 0.031 | | |

**Supplementary Table 30: Number of PDGFRβ$^+$ nuclei in PDGFRβ$^{KLF4\text{-}KO}$ mice.** Posterior summary with parameter estimates (Est.) and 95% credible intervals (2.5% - 97.5%) for the multilevel model a) *Counts ~ Genotype* + *(1* | *MouseID), family* = *negbinomial*. The values are presented in the log scale.

| Parameters | Est. | 2.5 % | 97.5 % |
|---|---|---|---|
| b_Intercept | 4.82 | 4.61 | 5.04 |
| b_GenotypeKO | -0.10 | -0.38 | 0.16 |
| sd_MouseID_Intercept | 0.19 | 0.02 | 0.37 |
| shape | 12.2 | 6.5 | 20.5 |
| Num.Obs. | 47 | | |
| R2 | 0.30 | | |

**Supplementary Table 31: PDGFRβ$^{+}$ labeling intensity in PDGFRβ$^{KLF4\text{-}KO}$ mice.** Posterior summary with parameter estimates (Est.) and 95% credible intervals (2.5% - 97.5%) for the multilevel model a) *Intensity ~ Area x Genotype + (1 | MouseID), family = student*.

| Parameters | Est. | 2.5 % | 97.5 % |
|---|---|---|---|
| b_Intercept | 0.29 | 0.06 | 0.54 |
| b_Area | 0.85 | 0.83 | 0.87 |
| b_GenotypeKO | -0.44 | -0.83 | -0.04 |
| b_Area:GenotypeKO | -0.32 | -0.34 | -0.29 |
| sd_MouseID_Intercept | 0.38 | 0.260 | 0.57 |
| sigma | 0.19 | 0.18 | 0.20 |
| Num.Obs. | 5588 | | |
| R2 | 0.81 | | |

**Supplementary Table 32: Number of PDGFRβ$^+$ nuclei in PDGFRβ$^{Cre}$ and KLF4$^{Flox}$ mice.** Posterior summary with parameter estimates (Est.) and 95% credible intervals (2.5% - 97.5%) for the multilevel model *Cells ~ Genotype + (1 | MouseID), family = student*.

| Parameters | Est. | 2.5 % | 97.5 % |
|---|---|---|---|
| b_Intercept | 167 | 146 | 188 |
| b_GenotypeCre | -51 | -78 | -24 |
| b_GenotypeFlox | -2.5 | -33 | 27 |
| sd_MouseID_Intercept | 10 | 0.4 | 25 |
| sigma | 10 | 1.1 | 26 |
| Num.Obs. | 10 | | |
| R2 | 0.883 | | |
| R2 Marg. | 0.820 | | |

**Supplementary Table 33: PDGFRβ$^+$ labeling intensity in PDGFRβ$^{Cre}$ and KLF4$^{Flox}$ mice.** Posterior summary with parameter estimates (Est.) and 95% credible intervals (2.5% - 97.5%) for the multilevel model *Intensity ~ Area x Genotype + (1 | MouseID), family = student*.

| Parameters | Est. | 2.5 % | 97.5 % |
|---|---|---|---|
| b_Intercept | 0.2 | 0.09 | 0.4 |
| b_Area | 1.00 | 0.98 | 1.01 |
| b_GenotypeCre | -0.61 | -0.91 | -0.32 |
| b_GenotypeFlox | -0.20 | -0.54 | 0.11 |
| b_Area_GenotypeCre | -0.34 | -0.37 | -0.31 |
| b_Area_GenotypeFlox | -0.14 | -0.16 | -0.12 |
| sd_MouseID_Intercept | 0.19 | 0.10 | 0.36 |
| sigma | 0.12 | 0.11 | 0.13 |
| Num.Obs. | 1460 | | |
| R2 | 0.97 | | |
| R2 Marg. | 0.952 | | |
| ICC | 0.2 | | |

**Supplementary Table 34: Collagen-IV (ColIV) labeling intensity in PDGFRβ$^{KLF4\text{-}KO}$ mice.** Posterior summary with parameter estimates (Est.) and 95% credible intervals (2.5% - 97.5%) for the model *Intensity ~ Genotype, family = student*.

| Parameters | Est. | 2.5 % | 97.5 % |
|---|---|---|---|
| b_Intercept | 283 | 193 | 374 |
| b_GenotypeKO | 9.9 | -111 | 136 |
| sigma | 120 | 79 | 176 |
| Num.Obs. | 18 | | |
| R2 | 0.025 | | |

**Supplementary Table 35: CD31 area in PDGFRβ$^{KLF4\text{-}KO}$ mice.** Posterior summary with parameter estimates (Est.) and 95% credible intervals (2.5% - 97.5%) for the model *Area ~ Genotype, sigma ~ Genotype, family = student*.

| Parameters | Est. | 2.5 % | 97.5 % |
|---|---|---|---|
| b_Intercept | 3677 | 3399 | 3954 |
| b_sigma_Intercept | 5.8 | 5.2 | 6.5 |
| b_GenotypeKO | 272 | -472 | 999 |
| b_sigma_GenotypeKO | 1.05 | 0.23 | 1.86 |
| Num.Obs. | 18 | | |
| R2 | 0.044 | | |

**Supplementary Table 36: ColIV area in PDGFRβ$^{KLF4\text{-}KO}$ mice.** Posterior summary with parameter estimates (Est.) and 95% credible intervals (2.5% - 97.5%) for the model *Area ~ Genotype, family = student*.

| Parameters | Est. | 2.5 % | 97.5 % |
|---|---|---|---|
| b_Intercept | 2843 | 2010 | 3688 |
| b_GenotypeKO | 699 | -466 | 1825 |
| sigma | 1155 | 788 | 1687 |
| Num.Obs. | 18 | | |
| R2 | 0.093 | | |

**Supplementary Table 37: CollV area in PDGFRβ$^{KLF4\text{-}KO}$ mice using Picrosirius staining.** Posterior summary with parameter estimates (Est.) and 95% credible intervals (2.5% - 97.5%) for the model *Area ~ Genotype, family = student*.

| Parameters | Est. | 2.5 % | 97.5 % |
|---|---|---|---|
| b_Intercept | 3757 | 2168 | 5352 |
| b_RegionCtx | 670 | 24 | 1904 |
| b_GenotypeKO | 442 | 12 | 1409 |
| b_RegionCtx_GenotypeKO | 511 | 14 | 1573 |
| sigma | 3974 | 3062 | 5150 |
| Num.Obs. | 36 | | |
| R2 | 0.024 | | |
