## Supplementary methods for "Brain PDGFRβ^+^ cells exhibit diverse reactive phenotypes after stroke without requiring KLF4"

Herein, we provide further details on the statistical modeling and analysis methods for the article **“Brain PDGFRβ$^+$ cells exhibit diverse reactive phenotypes after stroke without requiring KLF4”**. The raw images and data are available in Zenodo (10.5281/zenodo.10553084) and Open Science Framework (OSF) (10.17605/OSF.IO/74MQN) repositories. Note that some resources are not rendered in the OSF repository and users should download them to access the content. Likewise, the complete analysis pipelines in R (Quarto notebooks) or Python (Jupyter notebooks) are available in the GitHub repository (https://github.com/elalilab/Stroke_PDGFR-B_Reactivity).

Otherwise stated, we analyzed data by fitting case-specific Bayesian linear and non-linear regression models using the brms R package (Bürkner, 2017). For point pattern analysis (PPA), we employed the spatstat package (Baddeley et al., 2005) and associated multiple point process models (*ppm*) and relative distribution (*rhohat*) functions. Given the heterogeneity of our approaches, we thoroughly describe data handling and statistical modeling (including priors, model validation, and model comparison) in fully annotated Quarto notebooks (QNs). Moreover, we used the modelsummary package to write latex (.tex) and .html summary tables displaying model results. This material is available in the “Tables/tex” and “Tables/html” components of the OSF repository. In the following sections, we provide a tailored description for the analysis of each dataset. We remark that this content contains direct links to single files (i.e. Widefield_5x_Ipsilateral_Gfap-Pdgfrb.zip) hosted in the Zenodo or OSF repositories, or routes to the OSF components (Datasets, Images_Processing) and specific folders when referring to several files (i.e. StatisticalModels_brms/Widefield_5x_Ipsilateral_Gfap-Pdgfrb_Shrinkage). We note that we have had problems running brms in the last version of R and are unaware if these issues still exist. Be aware that we employed R 4.1.2 and the last available versions of all R-packages. Please check the sessionInfo() in each QN for further details. We also expect warnings opening QuPath projects, Ilastik, and CellProfiller pipelines because of file paths. We provide all the required files in our repositories to reproduce our results, but the user must download the material and change the file path accordingly. This does not apply to QN, JN, and .csv files if the user establishes an R-project hosting the files. Do not hesitate to raise an issue on GitHub or contact the research team if some issues arise.

**Supplementary Method 1: Analysis of the hemispheric shrinkage.**

| Component | Link |
|---|---|
| Figures | Figure 2A-B, Suppl. Figure 1D-E |
| Raw Images | Widefield_5x_Ipsilateral_Gfap-Pdgfrb.zip |
| Low-resolution images | Widefield_5x_Ipsilateral_Gfap-Pdgfrb.zip |
| Image Processing | Images_Processing/Widefield_5x_Ipsilateral_Gfap-Pdgfrb |
| Raw Data | Raw_Widefield_5x_Ipsilateral_Gfap_Str_QuPathAnnotations.csv |
| Processed data | Widefield_5x_Ipsilateral_Gfap-Pdgfrb_Inten.csv |
| Statistical models | StatisticalModels_brms/Widefield_5x_Ipsilateral_Gfap-Pdgfrb_Shrinkage |
| Analysis notebook | Widefield_5x_Ipsilateral_Gfap-Pdgfrb_Handling.qmd<br>Widefield_5x_Ipsilateral_Pdgfrb_Shrinkage.qmd |

We acquired widefield-stitched images at 5x magnification of the ipsilateral hemisphere of PDGFRβ$^{tdTomato}$ mice to analyze hemispheric shrinkage (atrophy). Using a CellProfiler script, we quantified the hemispheric area in an unbiased manner. We cleaned and processed the raw data as specified in QN. The resulting dataset includes the variables analyzed in the following sections: Analysis of cell reactivity (Integrated density) for PDGFRβ and GFAP$^{+}$ and Analysis of PDGFRβhigh and PDGFRβlow populations. For the analysis of brain shrinkage, we fitted various statistical models to the data, and we made model selection aided by the Watanabe-Akaike Information Criterion (WAIC) or Leave-one-out (Loo) cross-validation (Vehtari et al., 2016). We performed statistical inference based on a splines model (with heteroskedasticity):

$$Shrinkage \sim s(DPI, k = 5), sigma \sim DPI, family\ =\ student$$

We calculated the corresponding linear derivatives to interpret the effects of DPI using modelbased package (Makowski et al., 2020) as specified in the QN. We share the raw model summary or derivatives as .tex tables in “Tables/tex” component of the OSF repository. For the visualization of the results, we obtained posterior draws using the *fitted* function from stats package and generated a line graph using *stat_lineribbon()* with 0.5, 0.8, and 0.95 posterior credible intervals (CI) (blue Brewer scale).

**Supplementary Method 2: Analysis of the reactivity (Integrated density) of PDGFRβ$^{tdTomato}$ and GFAP$^{+}$ cells in the ipsilateral hemisphere.**

| **Component** | **Link** |
|---|---|
| Figures | Figure 2C-E and Suppl. Figure 1A-E |
| Raw Images | Widefield_5x_Ipsilateral_Gfap-Pdgfrb.zip |
| Low-resolution images | Widefield_5x_Ipsilateral_Gfap-Pdgfrb.zip |
| Image Processing | Images_Processing/Widefield_5x_Ipsilateral_Gfap-Pdgfrb |
| Raw Data | Raw_Widefield_5x_Ipsilateral_Gfap-Pdgfrb_AreaIntensity.csv |
| Processed data | Widefield_5x_Ipsilateral_Gfap-Pdgfrb_Inten.csv |
| Statistical models | StatisticalModels_brms/Widefield_5x_Ipilateral_Gfap-Pdgfrb_Inten |
| Analysis notebook | Widefield_5x_Ipsilateral_Pdgfrb_IntDen.qmd<br>Widefield_5x_Ipsilateral_Gfap_IntDen.qmd |

We analyzed widefield-stitched images at 5x magnification of PDGFRβ$^{tdTomato}$/GFAP$^{+}$ immunolabeling in brain sections (see the protocol) to quantify the integrated density (IntDen) (mean intensity x labeled area) of PDGFRβ and GFAP. We made the measurements in the ipsilateral hemisphere using a FIJI script and cleaned the data for analysis as specified in the QN. Based on WAIC, we made statistical inferences with a spline model that included heteroskedasticity (the estimation of sigma given different group variance):

$$IntDen \sim s(DPI, k = 5), sigma \sim DPI, family\ =\ student$$

We calculated the corresponding linear derivatives as described previously. We share the raw model summary or derivatives as .tex tables in the "Tables/tex" component of the OSF repository. We visualized the results using the *conditional_effects* function from brms with additional ggplot aesthetics. Additionally, we investigated the relationship between integrated density (IntDen) and hemispheric area (Tissue_Area) using the spline model:

$$IntDen \sim t2(DPI, TisseArea), family\ =\ guassian$$

The results are shown using the *conditional_smooths* function from brms with a Viridis scale. Please note that links in this section are for PDGFRβ. Analogous resources for GFAP are available in the OSF repository.

**Supplementary Method 3: Analysis of PDGFRβ$^{high}$ and PDGFRβ$^{low}$ subpopulations in the ipsilateral hemisphere.**

| **Component** | **Link** |
|---|---|
| Figures | Figure 2F-G, Suppl. Figure 1F. |
| Raw Images | Widefield_5x_Ipsilateral_Gfap-Pdgfrb.zip |
| Low-resolution images | Widefield_5x_Ipsilateral_Gfap-Pdgfrb.zip |
| Image Processing | Images_Processing/Widefield_5x_Ipsilateral_Gfap-Pdgfrb |
| Raw Data | Raw_Widefield_5x_Ipsilateral_Pdgfrb_QuPathAnnotations.csv<br>Raw_Widefield_5x_Ipsilateral_Pdgfrb_Str_QuPathAnnotations.csv |
| Processed data | Widefield_5x_Ipsilateral_Gfap-Pdgfrb_Inten.csv<br>Widefield_5x_Ipsilateral_Gfap-Pdgfrb_Inten_Str.csv |
| Statistical models | StatisticalModels_brms/Widefield_5x_Ipsilateral_Pdgfrb_LowHigh |
| Analysis notebook | Widefield_5x_Ipsilateral_Pdgfrb_LowHigh.qmd<br>Widefield_5x_Ipsilateral_Pdgfrb_LowHigh_Str.qmd |

We acquired widefield-stitched images at 5x magnification of sections in the ipsilateral hemisphere of PDGFRβ$^{tdTomato}$ mice to analyze the populations of PDGFRβ$^{high}$ and PDGFRβ$^{low}$ cells. We unbiasedly detected and quantified cells using QuPath (Bankhead et al., 2017). We loaded the images into a QuPath project and processed the files using a .groovy batch processing script. This pipeline makes use of machine learning-based classifiers (available in the QuPath project folder in the OSF repository) that label typical PDGFRβ cells as non-reactive (PDGFRβ$^{low}$) or increased PDGFRβ expression as reactive cells (PDGFRβ$^{high}$). We handled the raw data using the same QN employed for brain shrinkage and exported the results to the same dataset. To analyze cell proportions, we fitted Bayesian models using a binomial family distribution, where the response variable represents a series of Bernoulli trials (PDGFRβ$^{high}$ or PDGFRβ$^{low}$ cells) in a fixed number of independent trials (PDGFRβ$^{total}$). For additional details concerning this modeling strategy, validation, and model comparison, please refer to the associated QN. We performed statistical inference using a spline model:

$$PDGFR\beta_{High} \mid trials(PDGFR\beta_{Total} \sim s(DPI, k = 5), family \;=\; binomial$$

We visualized the results using the *conditional_effects* function from brms with additional ggplot aesthetics and shared the model summary as a .tex table in the OSF repository. Please note that the modelbased package does not have functions to generate linear derivatives from binomial models.

We performed the same procedure for an image set derived from mice with only striatal lesions. We generated a different Qupath project because the staining and imaging for these animals were done on a different day. The analysis pipeline is detailed in the QN and all the associated resources are analogously present in the OSF repository.

**Supplementary Method 4: Point pattern analysis (PPA) for PDGFRβ$^+$ cells and GFAP$^+$ cells in the ipsilateral hemisphere.**

| **Component** | **Link** |
|---|---|
| Figures | Figure 1H-I, Suppl. Figure 1F, Suppl. Figure 2A,F |
| Raw Images | Widefield_5x_Ipsilateral_Gfap-Pdgfrb.zip |
| Low-resolution images | Widefield_5x_Ipsilateral_Gfap-Pdgfrb.zip |
| Image Processing | Images_Processing/Widefield_5x_Ipsilateral_Gfap-Pdgfrb |
| Raw Data | Widefield_5x_Ipsilateral_Gfap-Pdgfrb_QuPath.zip<br>Widefield_5x_Ipsilateral_Gfap-Pdgfrb_QuPath_Str.zip |
| Processed data | Widefield_5x_Ipsilateral_Gfap-Pdgfrb-Dapi_Coordinates.zip<br>Widefield_5x_Ipsilateral_Gfap-Pdgfrb-Dapi_Str_Coordinates.zip |
| Point patterns | Widefield_5x_Ipsilateral_Pdgfrb-Gfap_PPP.rds<br>Widefield_5x_Ipsilateral_Pdgfrb-Gfap_Str_PPP.rds |
| Statistical models | N/A - mppm and rhohat models |
| Analysis notebook | Widefield_5x_Ipsilateral_Gfap-Pdgfrb_Covariance.qmd<br>Widefield_5x_Ipsilateral_Gfap-Pdgfrb_Covariance_Str.qmd |

We acquired widefield-stitched images at 5x magnification of PDGFRβ$^{tdTomato}$/GFAP$^+$ immunolabeling in brain sections (see the protocol) to analyze the distribution and covariance of PDGFRβ$^+$ and GFAP$^+$ cells in the ischemic hemisphere. We used QuPath (including machine-learning classifiers) to detect PDGFRβ$^{tdTomato}$, GFAP$^+$, and DAPI$^+$ cells and obtain single-cell *xy* coordinates. DAPI was used only to define the observation window of PDGFRβ and GFAP point patterns. We handled the raw data to save cell coordinates (per mouse) in individual .csv files for further processing (see QN). Subsequently, we created point patterns using the *ppp* function from spatstat as specified in the QN, and stored them as .rds (R-software) hyper-frames with their respective grouping variables. These objects are available in the OSF repository to encourage their reuse for validation, research, or educational purposes.

We used spatstat-associated functions to generate density kernels (0.2 sigma) and conduct PPA (see QN). Then, we used *rhohat* function to calculate the relative spatial intensity of PDGFRβ$^+$ cells with respect to the density kernels of GFAP$^+$ cells. In this context, spatial intensity refers to the number of objects per unit area, while maintaining all the spatial information available. Importantly, we passed the *rhohat* function with the argument *do.CI = FALSE* to allow further pooling per DPI with the *pool* function. The source code for generating density kernel plots and graphs of relative distribution **(as shown in Figure 1H-I and Suppl. Figure 1F)** are available in the QN.

Next, we fitted a spatial model for replicated point patterns using the *mppm* function from spatstat. This function assumes that the points follow an inhomogeneous Poisson process with an intensity function λ that varies over space μ. Please see the QN for further details. We fitted nested models for PDGFRβ$^{high}$ (Figure 1I-J) and PDGFRβ$^{low}$ **(Suppl. Figure 1H)**:

$$PDGFR\beta \sim DPI/GFAP\ Kernel, family\ =\ Poisson$$

The estimates with 95% confidence intervals (output on the log scale) are available in Supplementary Tables but not as a .tex table in the OSF repository. We performed an analogous procedure for mice with only striatal lesions (as explained previously).

**Supplementary Method 5: Analysis of GFAP$^{+}$ cell convex hull in the ipsilateral hemisphere.**

| **Component** | **Link** |
|---|---|
| Figures | Suppl. Figure 1F-G |
| Raw Images | Widefield_5x_Whole_Gfap-Pdgfrb.zip |
| Low-resolution images | Widefield_5x_Whole_Gfap-Pdgfrb.zip |
| Image Processing | Images_Processing/Widefield_5x_Whole_Gfap-Pdgfrb |
| Raw Data | N/A - Manual measurements |
| Processed data | Widefield_5x_Whole_Gfap_Coverage.csv |
| Statistical models | StatisticalModels_brms/Widefield_5x_Whole_Gfap_Convex |
| Analysis notebook | Widefield_5x_Ipsilateral_Gfap_Convex.qmd |

We analyzed widefield-stitched images at 5x magnification of PDGFRβ$^{tdTomato}$/GFAP$^{+}$ immunolabeling in brain sections to quantify the convex hull area (area covered by GFAP$^{+}$ cells) in the ipsilateral hemisphere. We performed manual measurements of the are covered by GFAP$^{+}$ cells using the standard *freehand section* and *measure* functions in FIJI **(as shown in Suppl. Figure 1G, yellowed encircled area)**. Previous research has shown that this area is fully populated by NeuN$^{+}$ cells and therefore corresponds to viable tissue (Manrique-Castano et al., 2024). Dataset handling and model diagnostics/comparison are specified in the QN. We performed statistical inference with a linear model:

$$Healthy\_Ratio \sim DPI, family\ =\ student$$

We share the summary table and show the results as a line plot using the same strategy as for brain shrinkage.

**Supplementary Method 6: Analysis of PDGFRβ⁺ cell count in tessellated GFAP⁺ areas in the ipsilateral hemisphere.**

| Component | Link |
|---|---|
| Figures | Figure 1K, L, Suppl. Figure 2A-B |
| Raw Images | Widefield_10x_ROIs_Gfap-Pdgfrb.zip |
| Low-resolution images | Widefield_10x_ROIs_Gfap-Pdgfrb.zip |
| Image Processing | Images_Processing/Widefield_10x_ROIs_Gfap-Pdgfrb |
| Raw Data | Raw_Widefield_10x_ROIs_Gfap-Pdgfrb_Tessellations.csv<br>Widefield_10x_ROIs_Gfap-Pdgfrb_QuPath.zip<br>Widefield_10x_ROIs_Gfap-Pdgfrb_QuPath_Str.zip |
| Processed data | Widefield_10x_ROIs_Gfap-Pdgfrb-Dapi_Coordinates.zip<br>Widefield_10x_ROIs_Gfap-Pdgfrb-Dapi_Coordinates_Excluded.zip |
| Point patterns | Widefield_10x_ROIs_Gfap-Pdgfrb_PPP.rds |
| Statistical models | StatisticalModels_brms/Widefield_10x_ROIs_Gfap-Pdgfrb_Covariance |
| Analysis notebook | Widefield_10x_ROIs_Gfap-Pdgfrb_Handling.qmd<br>Widefield_10x_ROIs_Gfap-Pdgfrb_Covariance.qmd |

We acquired widefield images at 10x magnification in the injured cortex (Ctx, cortex), injured striatum (Str, striatum), and the healthy perilesional cortex (Peri, perilesion) of PDGFRβ$^{tdTomato}$/GFAP$^{+}$ immunolabeling in brain sections (see the protocol) to analyze the relative distribution of PDGFRβ$^{+}$ conditional on the spatial intensity of reactive astrocytes. We detected PDGFRβ$^{tdTomato}$, GFAP$^{+}$, and DAPI$^{+}$ cells using batch processing scripts for QuPath and created point patterns using DAPI$^{+}$ cells to establish the observation window. We note that for this dataset we excluded mouse 80 (sham) and mouse 74 (3D), given that no detection of GFAP$^{+}$ cells (*NULL* data) prevented the successful construction of the hyper-frame. We analyzed the covariance of PDGFRβ$^{+}$ and GFAP$^{+}$ cells using GFAP-based tessellations. These are non-overlapping regions segregating areas of low and high GFAP spatial intensity. We built tessellations using two quantiles for GFAP$^{low}$ and GFAP$^{high}$ spatial intensity and the *tess* function from spatstat. Then, we obtained the number of PDGFRβ$^{+}$ in each region using the *quadratcount* function as shown in QN. We employed two modeling strategies for this dataset. First, we fitted a linear model:

$$PDGFR\beta \sim Region * GFAP, family\ =\ hurdle\_lognormal$$

to estimate the number of PDGFRβ$^{+}$ cells conditioning on the interaction between region (perilesion, striatum, or cortex) and GFAP spatial intensity (low, high). We used the hurdle_lognormal family distribution to account for the overdispersion of positive data values containing zeros. Model diagnostics are available in the QN. We show the results using conditional_effects for point estimates and their uncertainty and comparisons (*emmeans*) between regions using stat_slab + stat_pointinterval. The summary table is available in OSF.

In the second place, we could not fit linear models conditioning on the three grouping variables (region, GFAP tessellation, DPI), given that MCMC chains did not converge on a stable posterior distribution. Therefore, we fitted separate models for the perilesion, striatum, and, cortex with the same model formula and distribution family. We show the results as previously described. The source code for the representative images displayed in the research article is available in the QN.

**Supplementary Method 7: Analysis of perivascular and parenchymal PDGFRβ⁺ cells in defined ROIS of the ipsilateral hemisphere.**

| **Component** | **Link** |
|---|---|
| Figures | Figure 3B-C, L, Suppl. Figure 3A-G |
| Raw Images | Widefield_10x_ROIs_CD31-Pdgfrb.zip |
| Low-resolution images | Widefield_10x_ROIs_CD31-Pdgfrb.zip |
| Image Processing | Images_Processing/Widefield_10x_ROIs_CD31-Pdgfrb |
| Raw Data | Raw_Widefield_10x_ROIs_CD31-Pdgfrb_Coloc.csv |
| Processed data | Widefield_10x_ROIs_CD31-Pdgfrb_Coloc.csv |
| Statistical models | StatisticalModels_brms/Widefield_10x_ROIs_CD31-Pdgfrb_Coloc |
| Analysis notebook | Widefield_10x_ROIs_CD31-Pdgfrb_Coloc.qmd |

We acquired widefield images at 10x magnification in the injured cortex (Ctx, cortex), injured striatum (Str, striatum), and the healthy perilesional cortex (Peri, perilesion) of $PDGFR\beta^{tdTomato}$ /CD31 imunolabeling in brain sections (see the protocol) to analyze the proportions of perivascular (vascular-associated) and parenchymal (no vascular-associated) $PDGFR\beta^{+}$ cells after ischemic stroke.

We used the pixel classification tool from Ilastik (Berg et al., 2019) to enhance the detection of $PDGFR\beta^{tdTomato}$ cells. Subsequently, we imported .png segmentation masks into CellProfiler (Stirling et al., 2021) to perform unbiased/automated cell detection using the *MaskImage* and *IdentifyPrimaryObjects* modules. On the other hand, we detected $CD31^{+}$ cells using the *IdentifyPrimaryObjects* module and dilated (*DilateObjects*) the objects with disk = 2 as the structuring element. With this procedure, we aimed to capture the $PDGFR\beta^{+}$ cells interacting with $CD31^{+}$ vasculature from a side plane (see graph below). Finally, we related the $PDGFR\beta^{+}$ and $CD31^{+}$ objects using the RelateObjects module and exported the data as a .csv file. The complete CellProfiler pipeline is available in the OSF repository.

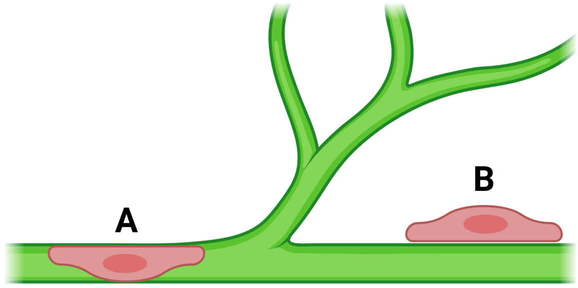

PDGFRβ+ perivascular pericytes can be detected directly over the vasculature (A) or from a side plane (B). We dilated PDGFRβ+ objects to capture cells from both imaging planes.

We analyzed the changes in the proportion of parenchymal PDGFRβ+ cells by fitting independent binomial Bayesian models per region (perilesion, striatum, and cortex), as described previously:

$$PDGFR\beta_{parenchymal} \mid trials(PDGFR\beta_{Total} \sim s(DPI, k = 4), family = binomial$$

The modeling strategy, validation, and model comparison are available in QN. We visualized the results using the *conditional_effects* function from brms with additional ggplot aesthetics and shared the posterior estimates as .tex tables in the OSF repository. Please note that the modelbased package does not have functions to generate linear derivatives from binomial models.

Complementarily, we modeled the number (not the proportion) of PDGFRβ+ cells with the interaction between DPI and Regions (cortex, striatum, perilesion) as predictors:

$$PDGFR\beta_{parenchymal} \sim DPI * Region, family = hurdle\_lognormal$$

We used a hurdle_lognormal model to account for overdispersion and a high proportion of zeros in perilesional areas. We show these posterior estimates using *conditional_effects* and provide the .tex table in the OSF repository. From this model, we also calculated the contrast between the number of PDGFRβ+ cells in the cortex and striatum, and plot them using stat_slab() + stat_pointinterval().

Next, we compared mice with cortico-striatal and striatal-only lesions at 14 and 30 DPI to evaluate the influence of cortical-striatal injuries on the appearance of PDGFRβ+ parenchymal cells. For this purpose, we used two modeling strategies (see QN). First, given the exploratory data analysis, we fitted a linear model with heteroscedasticity (predicting sigma) to estimate the total number of PDGFRβ+ cells having the interaction of DPI and type of lesion as predictors:

$PDGFR\beta$[redacted]$\sim DPI * Lesion, sigma \sim DPI * Lesion, family = student$

We show the posterior estimates for the number of cells and sigma with their respective posterior summary as described previously in this section.

Secondly, we modeled the proportion of PDGFRβ+ cells with the interaction of DPI and lesion type as predictors:

$$PDGFR\beta_{parenchymal} \mid trials(PDGFR\beta_{Total} \sim DPI, family \ = \ binomial$$

See the posterior summary in the OSF repository and the QN for further details on the modeling strategy, diagnostics, and source code for data visualization.

Finally, we investigated the effects of 4-OHT-mediated Cre-recombination on the number of parenchymal PDGFRβ+ cells. We performed an analogous tamoxifen injection in sham animals and harvested the brains at the same time points as done for MCAO mice. We analyzed the cortex of Sham and MCAO mice using a linear model:

$$PDGFR\beta_{parenchymal} \sim DPI \ * \ Condition, family \ = \ student$$

We show the results and posterior summary as previously described in this section.

**Supplementary Method 8: Fluorescence-activated cell sorting (FACS) of brain PDGFRβ⁺ cells.**

| Component | Link |
|---|---|
| Figures | Figure 3D-G; Suppl. Figure 4. |
| Raw Images | N/A |
| Low-resolution images | N/A |
| Image Processing | N/A |
| Raw Data | FACS |
| Processed data | Data_Processed/Facs_Pdgfrb_Cells-Dead.csv |
| Point patterns | N/A |
| Statistical models | StatisticalModels_brms/Facs_Psgfrb-Dapi |
| Analysis notebook | FlowCytometry_Pdgfrb_Processing.qmd<br>FlowCytometry_Pdgfrb_Analysis.qmd |

We use the adult brain dissociation kit (Miltenyi Biotec, cat # 130-107-677) to sort PDGFRβ$^{+}$ cells from PDGFRβ$^{tdTomato}$ mice at 14 DPI, as detailed in the protocol. To compare the proportion of dead cells between the ipsilateral and contralateral hemispheres, we stained the cells with Fixable Viability Stain 780 (Live/Dead; BD Biosciences, cat #3565388). We analyzed the resulting .fcs files—available in "Datasets/FACS" component of the OSF repositories—using open-source packages from Bioconductor.

We loaded the files using the *read.flowset* function from the flowCore package (Hahne et al., 2009). Next, we compensated the samples using the *compensate* function from flowWorkspace and performed quality checks with the *flow_auto_qc* function from the flowAI package (Monaco et al., 2016). These procedures ensured the removal of anomalies related to the flow rate, signal acquisition, and dynamic range. We have made the quality check files available in the OSF repository. Subsequently, we defined the gating strategy and visualized the clee proportions using functions from the ggcyto package (Van et al., 2018). **Supplementary Figure 4** presents plots of the complete sample set. The complete handling and processing pipeline of the raw .fcs files is provided in the QN.

We modeled the number of PDGFRβ$^{+}$ cells per hemisphere using a linear model (single 14 DPI):

$$PDGFR\beta_{Cells_{\blacksquare}} \sim Hemisphere, family \ = student$$

We show the results and posterior summary in the OSF repository.

Second, we modeled the number of PDGFRβ$^{+}$ dead cells distributed by hemisphere at the same DPI. We fitted two models:

Model 1 = $PDGFR\beta_{Dead_{\blacksquare}} \sim Hemisphere, family \ = student$

Model 2 = $PDGFR\beta_{Dead_{\blacksquare}} \sim Hemisphere, sigma \sim Hemisphere, family \ = student$

The posterior predictive checks showed that the model with heteroskedasticity (Model 2) deviates less from the observed data and is less penalized by out-of-sample prediction (see QN). We show the results and posterior summary for this model in the OSF repository.

**Supplementary Method 9: PPA and colocalization analysis of Ki67⁺ / PDGFRβ⁺ cells in the ipsilateral hemisphere.**

| **Component** | **Link** |
|---|---|
| Figures | Figure 3H-K |
| Raw Images | Widefield_10x_Ipsilateral_Ki67-Pdgfrb(a).zip<br>Widefield_10x_Ipsilateral_Ki67-Pdgfrb(b).zip |
| Low-resolution images | Widefield_10x_Ipsilateral_Ki67-Pdgfrb.zip |
| Image Processing | Images_Processing/Widefield_10x_Ipsilateral_Ki67-Pdgfrb |
| Raw Data | Data_Raw/Widefield_10x_Ipsilateral_Ki67-Pdgfrb |
| Processed data | Data_Processed/Widefield_10x_Ipsilateral_Ki97-Pdgfrb |
| Point patterns | Widefield_10x_Ipsilateral_Ki67-Pdgfrb_PPP.rds |
| Statistical models | StatisticalModels_brms/Widefield_10x_Ipsilateral_Ki67-Pdgfrb |
| Analysis notebook | Widefield_10x_Ipsilateral_Ki67_PPA.qmd<br>Widefield_10x_Ipsilateral_Ki67-Pdgfrb_Coloc.qmd |

We acquired widefield images at 10x magnification of PDGFRβ$^{tdTomato}$ / Ki67 imunolabeling in brain sections (see the protocol) in the ipsilateral hemisphere to analyze the proportion and distribution of Ki67$^{+}$/PDGFRβ$^{+}$ cells following injury. First, we performed pixel classification for Ki67$^{+}$ and PDGFRβ$^{tdTomato}$ using Ilastik to enhance cell detection in CellProfiler. We then imported the segmentation masks into cellProfiler and conducted unbiased, automated object detection using the IdentifyPrimeryObjects module. Next, we filtered Ki67 and PDGFRβ objects to eliminate small particles and abnormally large objects. Finally, we related the objects using the RelateObjects module and exported the data into .csv files. We share the complete pipeline in the OSF repository, which includes the generation of outlines and crops for visualization, validation, and reuse.

Secondly, we created point patterns using *xy* coordinates for individual Ki67$^{+}$ and PDGFRβ$^{+}$ cells obtained from CellProfiler, as previously specified. We used the

PDGFRβ$^{tdtomato}$ coordinates to define the observation window. We then analyzed the distribution of Ki67$^{+}$ cells conditioned on the spatial intensity of PDGFRβ by calculating the relative distribution (rhohat) and fitting a point process model (mppm) as described earlier. For additional details, please refer to the QN. Afterward, we evaluated the PDGFRβ$^{+}$ / Ki67 colocalization in the ipsilateral hemisphere using the summary .csv file from CellProfiler containing cell counts per brain. We estimated the percentage of colocalized cells using a spline model:

$$PDGFR\beta_{\blacksquare} \sim s(DPI, k = 5), family = lognormal$$

and calculated the corresponding linear derivatives, as described previously. We share the raw model summary or derivatives as .tex tables in the "Tables/tex" component of the OSF repository. We visualized the results using the *conditional_effects* with additional ggplot aesthetics.

**Supplementary Method 10: Analysis of Ki67$^+$ / PDGFRβ$^{tdTomato}$ cells in defined ROIs of the ipsilateral hemisphere.**

| **Component** | **Link** |
|---|---|
| Figures | Figure 3H-I |
| Raw Images | Confocal_20x_ROIs_Ki67-Pdgfrb-CD31.zip |
| Low-resolution images | Confocal_20x_ROIs_Ki67-Pdgfrb-CD31.zip |
| Image Processing | Images_Processing/Confolcal_20x_ROIs_Ki67-Pdgfrb-CD31 |
| Raw Data | Data_Raw/Confolcal_20x_ROIs_Ki67-Pdgfrb-CD31 |
| Processed data | Confocal_20x_ROIs_Ki67-Pdgfrb_Coloc.csv |
| Statistical models | StatisticalModels_brms/Confocal_20x_ROIsl_Ki67-Pdgfrb_Coloc |
| Analysis notebook | Confocal_20x_ROIs_Ki67-Pdgfrb_Coloc.qmd |

We acquired confocal images at 20 x magnification of PDGFRβ$^{tdTtomato}$ / Ki67 imnuolabeling in brain sections (see the protocol) in defined ROIs (healthy cortical peri-lesional region, injured striatum, and injured cortex) to analyze the proportion of vascular (attached to CD31$^+$ cells) Ki67$^+$ / PDGFRβ$^+$ cells. To enhance cell detection in CellProfiler, we used pixel classification for Ki67$^+$ and PDGFRβ$^{tdTomato}$ using Ilastik. For Ki67, we applied the Smooth and MaskImage modules (utilizing segmentation masks from Ilastik) before proceeding to detect individual cells with IndentifyPrimeryObjects. We then filtered the objects using FilterObjects (*maximum value only* = 2000) to eliminate abnormalities and shrank the remaining objects with ExpandOrShrinkObjects by *10 pixels* to reduce false-positive colocalization caused by cell proximity. We conducted an analogous procedure for PDGFRβ without object shrinkage. Finally, we detected CD31$^+$ vasculature by smoothing and merging using SplitOrMergeObjects with a maximum pixel distance of 2, followed by filtering the objects as aforementioned.

Using this analysis pipeline, we quantified (1) the number of perivascular (CD31-attached) PDGFRβ$^+$ cells; (2) the number of vascular Ki67$^+$ / PDGFRβ$^+$ cells; and (3) the total number of Ki67$^+$ / PDGFRβ$^+$ cells, including both vascular and non-vascular associated cells. The CellProfiler pipeline, along with a complete set of object images and outlines, is available in the OSF repository. To investigate the total number of Ki67$^+$ / PDGFRβ$^+$ cells, we fitted a Bayesian model incorporating the interaction of DPI and brain region (perilesion, striatum, cortex) as predictors:

$$Cells \sim DPI * Region, family = negbinomial$$

We used a negative binomial distribution (family = negbinomial) to model counts and account for overdispersion. In contrast, to model the proportion of vascular associated (CD31-attached) Ki67$^+$ / PDGFRβ$^+$ cells, we employed the same predictors but used a binomial distribution as previously described:

$$Ki67/PDGFR\beta_{vascular} \mid trials(Ki67/PDGFR\beta_{total} \sim DPI * Region, family = binomial$$

Additional modeling details are provided in the QN. We present the model results using the conditional_effects function and share the posterior summaries in the OSF repository.

**Supplementary Method 11: Morphological analysis of PDGFRβ⁺ cells in defined ROIs of the ipsilateral hemisphere.**

| Component | Link |
| --- | --- |
| Figures | Figure 4A-C; Suppl. Figure 5A-D |
| Raw Images | Confocal_40x_ROIs_CD31-Pdgfrb-CD13.zip |
| Low-resolution images | Confocal_40x_ROIs_CD31-Pdgfrb_CD13.zip<br>Confocal_40x_ROIs_Pdgfrb_Crops.zip<br>Confocal_40x_ROIs_Pdgfrb_Segmented.zip |
| Image Processing | Images_Processing/Confolcal_40x_ROIs_CD31-Pdgfrb-CD13 |
| Raw Data | Data_Raw/Confolcal_40x_ROIs_Pdgfrb_Morph |
| Processed data | Confocal_40x_ROIs_Pdgfrb_Morph.csv |
| Statistical models | Confocal_40x_ROIs_Pdgfrb_Morph_Fit1.rds |
| Analysis notebook | Confocal_40x_ROIs_Pdgfrb_Morpho_BatchScript.qmd<br>Confocal_40x_ROIs_Pdgfrb_Morpho_Analysis.qmd |

We acquired confocal z-stack images at 40x magnification of PDGFRβ$^{tdTtomato}$ / CD31 / CD13 immunolabeling in brain sections (see the protocol) in defined ROIs (healthy cortical peri-lesional region, injured striatum, and injured cortex) to analyze the morphological features of cells. Using a FIJI script, we created z-projection images and manually cropped them to segment one or more well-differentiated cells (see resulting cells). Subsequently, we employed the Python libraries skimage and scipy to enhance, threshold, and generate masks of single cells, as demonstrated in the QN. We measure the morphological features of these masks using the regionprops and regionprops_table functions from skimage.measure.

Afterward, we used the Object Classification module from CellProfiler to perform a user-defined classification into “Amoeboid”, “Perivascular”, “Reticuloparenchymal”, and “Reticulite” cells. We utilized the manually cropped and segmented cells, along with the masks generated by the Python pipeline, as input for classification. We share the resulting raw data files (.csv per brain) containing the classification results and measured morphological traits, including *Intensity_sd, Intensity_Mean, Defects,*

*Convexity, Area, ConvexHull, BranchLenght, Branches, Diameter, EcuclideanDiameter*.

After loading, cleaning, and processing the raw data files, we created a single dataset containing the relevant grouping variables and morphological traits (see QN). To analyze the data, we performed principal component analysis (PCA) using the FactoMineR (Lê et al., 2008) and factoextra-R packages. Based on the resulting correlation matrix and biplot, we selected area, convex hull, branch length, and mean intensity as modeling variables. We then fitted a multinomial logistic regression model with these morphological features as predictors of cell class (amoeboid, perivascular, reticuloparenchymal, and reticulite):

$$Class \sim Area + ConvexHull + BranchLenght + Intensity\_Mean, family = categorical$$

The modeling strategy and diagnostics are fully described in the QN. We present the results as a spyder plot showing means (without displaying uncertainty) and provide individual plots for each morphological attribute using conditional_effects. The posterior summary is also available in the OSF repository.

**Supplementary Method 12: Analysis of covered area and Haralick features (texture) of PDGFRβ⁺ cells in defined ROIs of the ipsilateral hemisphere.**

| Component | Link |
| --- | --- |
| Figures | Figure 5A; Suppl. Figure 7A-D |
| Raw Images | Widefield_20x_ROIs_Pdgfrb.zip |
| Low-resolution images | Widefield_20x-ROIs_Pdgfrb.zip |
| Image Processing | Images_Processing/Widefield_20x_ROIs_Pdgfrb |
| Raw Data | Raw_Widefield_20x_ROIs_Pdgfrb_Haralick.csv<br>Data_Raw/Widefield_20x_ROIs_Pdgfrb/CellProfiler |
| Processed data | N/A – Raw data is clean |
| Statistical models | Widefield_20x_ROIs_Pdgfrb_Area_Fit1.rds<br>Widefield_20x_ROIs_Pdgfrb_Haralick_Fit1.rds |
| Analysis notebook | Widefield_20x_ROIs_Pdgfrb_Haralick_BatchScript.qmd<br>Widefield_20x_ROIs_Pdgfrb_Haralick.qmd |

We acquired widefield z-stack images at 20x magnification of PDGFRβ$^{tdTomato}$ / PDGFRα$^{+}$ immunolabeling in brain sections (see the protocol) in defined ROIs (healthy cortical perilesion, injured striatum, and injured cortex) to analyze the spatial and topological arrangement of cells. For this analysis, we focused exclusively on PDGFRβ$^{+}$ cells. Using FIJI, we created two-dimensional z-projections and analyzed the resulting images with CellProfiler. We identified single cells using the IndentifyPrimaryObjects module and measured each object's occupied area with the MeasureImageAreaOccupied module. We analyzed the cell area data using a linear model with DPI and Region (cortex, striatum, perilesion) as predictors (see QN):

$$Area \sim DPI\_cont * Region, family = student$$

We present the results using conditional_effects function with additional ggplot aesthetics and have made the posterior summary available in the OSF repository. Subsequently, we measured Haralick features (see table below) (Löfstedt et al., 2019) using the mahotas Python package (Coelho, 2013) as detailed in the QN.

| Haralick features | Description |
|---|---|
| Angular Second Moment (ASM) / Energy | Measures the uniformity of an image. A higher value indicates that the image has more uniform textures or constant regions. |
| Contrast | Represents the difference between the highest and the lowest intensity value in the co-occurrence matrix. It measures the amount of local variations present in an image. |
| Correlation | Measures the joint probability occurrence of the specified pixel pairs. It provides information about the linear dependency of gray levels in the neighboring pixels. |
| Sum of Squares / Variance | It provides a measure of the squared differences from the mean intensity value. |
| Inverse Difference Moment (IDM) / Homogeneity | Measures the local homogeneity of an image. The values are high when the local textures are consistent or homogeneous. |
| Sum Average | Represents the average intensity value of the co-occurrence matrix. |
| Sum Variance | Measures the variance of the sum of the intensity values from the average value in the co-occurrence matrix. |
| Sum Entropy | Represents the randomness or complexity in the sum of the intensity values of the co-occurrence matrix. |
| Entropy | Provides a measure of the randomness or complexity of the image. Higher values indicate more complex textures. |
| Difference Variance | Represents the variance in the differences between the intensity values of pairs of pixels. |
| Difference Entropy | Measures the randomness or complexity in the differences between the intensity values of pairs of pixels. |
| Informational Correlation 1 (Info Corr 1) | Represents the correlation between the occurrence of the specified pixel pairs and their average intensity values. |
| Informational Correlation 2 (Info Corr 2) | Provides another measure of the correlation between the occurrence of the specified pixel pairs and their average intensity values. It’s typically more sensitive to changes than Info Corr 1. |

We analyzed the Haralick features using PCA and presented the results as previously described. Based on the PCA, we selected entropy, contrast, and inverse difference moment (IDM) as the most relevant features (see QN). We performed statistical modeling by fitting a multinomial logistic regression model in brms to predict the brain region (perilesion, striatum or cortex) based on the selected Haralick features:

$$Region \sim Entropy\ +\ Contrast\ + IDM, family\ =\ categorical$$

Model diagnostics are available in the QN. We visualize the results for each feature using the conditional_effects function and have made the posterior summary in the OSF repository.

**Supplementary Method 13: Topological data analysis (TDA) of PDGFRβ⁺ cells in defined ROIs of the ipsilateral hemisphere.**

| **Component** | **Link** |
|---|---|
| Figures | Figure 5B-E; Suppl. Figure 7E-F |
| Raw Images | Widefield_20x_ROIs_Pdgfrb.zip |
| Low-resolution images | Widefield_20x-ROIs_Pdgfrb.zip |
| Image Processing | Images_Processing/Widefield_20x_ROIs_Pdgfrb |
| Raw Data | Data_Raw/Widefield_20x_ROIs_Pdgfrb/CellProfiler |
| Processed data | Data_Processed/Widefield_20x_ROIs_Pdgfrb/ |
| Statistical models | N/A |
| Analysis notebook | Widefield_20x_ROIs_Pdgfrb_TDA.ipynb |

We performed TDA using the same images and datasets from the previous section. We obtained single *xy* cell coordinates from CellProfiler to serve as input for TDA. Utilizing the GUDHI Python library (The GUDHI Project, 2023), we calculated Vietoris-rips complex (Chambers et al., 2010) with a maximum dimension of 2 and a radius of 100 (see JN). We have made available a full set of static and animated (see example) Vietoris-rips complex visualizations for reuse in research or educational settings.

To calculate persistent homology (Aktas et al., 2019; Otter et al., 2017), we generated point clouds and saved them as .npy Numpy arrays. We used the ripser Python library (Tralie et al., 2018) to compute persistent homology for dimension-0 (0D, connected components) and dimension-1 (1D, loops). The complete set of persistent diagrams is shared as .npy files for research purposes and as .png for visualization. Subsequently, we calculated Betti curves to analyze the evolution of the topological features revealed by persistent homology. For connected components (0D homology), we calculated normalized (to 1) Betti curves (steps = 300, max filtration = 300) to eliminate the bias by the number of components (cells). For loops (1D homology), we employed 100 steps and a maximum filtration value of 700 without normalization. We plotted the Betti curves per dimension and region as line plots using matplotlib (Hunter,

2007). To quantify the differences between DPIs, we calculated Bottleneck and Wasserstein distances (Hajij et al., 2018) using the *bottleneck_distance* and *wasserstein_distance* functions from the gudhi library, respectively. We obtained median estimates with their respective uncertainty by bootstrapping (Chernik, 2007) the samples with 1000 replications using the *choices* function from the random library. We present the results for Bottleneck and Wasserstein distances as DPI contrast heatmaps, displaying median values and 95% confidence intervals. For additional details on Vietoris-Rips complexes, persistent homology, Betti curves, and Bottleneck/Wasserstein calculations, please refer to the JN.

**Supplementary Method 14: Spatial and labeling intensity analysis of KLF4 in the ipsilateral hemisphere.**

| Component | Link |
|---|---|
| Figures | Figure 6A-C; Suppl. Figure 8A-C |
| Raw Images | Widefield_10x_Ipsilateral_KLF4.zip<br>Widefield_10x_Ipsilateral_KLF4.z01<br>Widefield_10x_Ipsilateral_KLF4.z02 |
| Low-resolution images | Widefield_10x_Ipsilateral_Klf4.zip |
| Image Processing | Images_Processing/Widefield_10x_Ipsilateral_Klf4 |
| Raw Data | Data_Raw/Widefield_10x_Ipsilateral_Klf4 |
| Processed data | Data_Processed/Widefield_10x_Ipsilateral_Klf4 |
| Point patterns | Widefield_10x_Ipsilateral_Klf4_PPP.rds |
| Statistical models | StatisticalModels_brms/Widefield_10x_Ipsilateral_Klf4 |
| Analysis notebook | Widefield_10x_Ipsilateral_Klf4_Exp.qmd |

We acquired widefield images at 10 x magnification of PDGFRβ$^{tdTomato}$ / KLF4 immunolabeling in brain sections (see the protocol) in the ipsilateral hemisphere to analyze the spatial distribution and signal intensity of KLF4 following ischemia. To improve cell detection in CellProfiler, we performed pixel classification for KLF4 using Ilastik. We detected KLF4$^+$ cells by using the segmentation masks as input for the IdentifyPrimeryObjects module in CellProfiler. Additionally, we detected DAPI$^+$ cells to define the observation window for point patterns. The CellProfiler pipeline includes the generation of cropped images representing the detection of single cells. We processed the raw files from CellProfiler to build point patterns (*ppp*), calculate the spatial intensity (*$intensity*), derive density kernels (*density*), and fit multiple point process models (*mppm*). Please refer to the QN for further details and the source code. To evaluate the distribution of KLF4 following injury, we fitted a mixed-effects model using spatstat, with the x-axis (representing distance from the ischemic area in the ipsilateral hemisphere) as a predictor varying by DPI:

$$KLF4 \sim x, random = \sim x \mid DPI, family = Poisson$$

Please note that the model summary in the QN is on the log scale.

To account for variance across different DPIs, we extracted the spatial intensity of $KLF4^{+}$ cells from the point patterns and fitted a linear model with heteroskedasticity using brms. Specifically, we modeled intensity as a function of DPI without an intercept and allowed the residual variance (σ) to vary with DPI:

$$Intensity \sim 0 + DPI, sigma \sim 0 + DPI, family = student$$

Model diagnostics and source code are available in the QN. We present the results as full posterior distributions per DPI for both the parameter estimates (*add_predicted_draws* + *stat_haleye* from tidybayes) and the residual standard deviation (sigma) (conditional_effects). We calculated the contrast between time points using the emmeans package (Lenth, 2024) as detailed in the QN. The posterior summary is shared as a .tex table in the OSF repository. We performed an analogous analysis for point patterns and estimated KLF4 mean intensity for brains derived from $PDGFR\beta^{KLF4\text{-}KO}$ mice. Subsequently, we analyzed the labeling intensity of KLF4 on a per-cell basis, as obtained from the CellProfiler pipeline. Due to computational limitations with the full dataset of 278,340 observations, we randomly subsampled 10% of the data using sample_frac(0.1) and fitted a multilevel model with a hurdle_lognormal() distribution to handle zero-inflated data and capture continuous observations:

$$Intensity \sim 0 + DPI + (1 \mid MouseID), family = hurdle\_lognormal()$$

We included MouseID as a random intercept to account for the hierarchical structure of the data and to obtain more precise estimates by integrating variance within mice. Modeling details and diagnostics are available in the QN. We present the results using *conditional effects*, rendering point estimates with 95% CI, and contrasts between DPIs using stat_pointinterval.

Additionally, we fitted a second multilevel model to evaluate the influence of the x-coordinates within the ipsilateral hemisphere on KLF4 labeling intensity. We centered the x-coordinates to mitigate bias from brain shrinkage and to facilitate parameter space exploration. Employing a similar modeling strategy with random intercepts for MouseID, we specified:

$$Intensity \sim 0 + DPI * x.coor + (1 \mid MouseID), family = hurdle\_lognormal()$$

We visualized the results using conditional_effects, rendering colored line plots per DPI with 95% CI. Additionally, we generated scatter plots of cell positions with a *Viridis* color scale to depict KLF4 immunolabeling intensity per cell.

**Supplementary Method 15: Analysis of PDGFRβ⁺ / KLF4⁺ cells in defined ROIs of the ipsilateral hemisphere.**

| **Component** | **Link** |
|---|---|
| Figures | Figure 6D-E; Suppl. Figure 8D |
| Raw Images | Confocal_20x_ROIs_Klf4-Pdgfrb-CD31(a).zip<br>Confocal_20x_ROIs_Klf4-Pdgfrb-CD31(b).zip |
| Low-resolution images | Confocal_20x_ROIs_Klf4-Pdgfrb_Overlay.zip |
| Image Processing | Images_Processing/Confocal_20x_ROIs_Klf4-Pdgfrb-CD31 |
| Raw Data | Data_Raw/Confocal_20x_ROIs_Klf4-Pdgfrb-CD31 |
| Processed data | Confocal_20x_ROIs_Klf4-Pdgfrb_Coloc.csv |
| Statistical models | StatisticalModels_brms/Confocal_20x_ROIs_Klf4-Pdgfrb-CD31 |
| Analysis notebook | Confocal_20x_ROIs_Klf4-Pdgfrb_Coloc.qmd |

We acquired confocal images at 20x magnification of PDGFRβ$^{tdTomato}$ / KLF4 / CD31 immunolabeling in brain sections (see the protocol) in defined ROIs (contralateral hemisphere, healthy perilesion, and injured cortex) to analyze PDGFRβ$^{tdTtomato}$ and KLF4 co-localization. This dataset differs from others in that we included the contralateral hemisphere instead of the injured striatum. We used the contralateral hemisphere as a control (ground truth) for colocalization error, given that in healthy tissue, KLF4 is prominently expressed in the vasculature by CD31⁺ brain endothelial cells but not in PDGFRβ⁺ perivascular cells.

To enhance cell detection and co-localization analysis in CellProfiler, we performed pixel classification for KLF4 using Ilastik. For analyzing colocalization, we employed a pipeline that included filling (FiilObjects), filtration (FilterObjects), and erosion (ErodeObjects) of PDGFRβ⁺ and KLF4⁺ cells. This approach aimed to reduce the rate of false positives in PDGFRβ / KLF4 co-localization due to the proximity of these two markers in the perivascular space. We calculated co-localization by cell counts using the RelateObjects module. Note that CD31 was included in this pipeline solely to generate reference visualizations, as shown in **Figure 6D**. The pipeline also generated images with overlaid contours (found in the "ObjectsOverlay" folder) for visualization and illustration purposes **(see Figure 6D)**.

We processed the CellProfiler output to obtain a clean dataframe containing the counts of PDGFRβ$^{+}$, KLF4$^{+,}$ and co-localized PDGFRβ$^{+}$ / KLF4$^{+}$ cells. Our statistical inference was based on four different binomial models, as previously described. First, we fitted a linear model (Mdl1) to explore the distribution of PDGFRβ$^{+}$/ KLF4$^{+}$ co-localized cells across distinct DPIs:

$$PDGFR\beta - KLF4 \mid trials(Klf4) \sim DPI, family \; = \; binomial$$

Secondly, using the same approach, we modeled (Mdl2) this cell distribution by brain regions— injured cortex (Ctx), healthy cortical perilesion (Peri), and contralateral hemisphere (Ctr)s—. Next, excluding data from day 0 (sham animals), we fitted a third model (Mdl3) to evaluate PDGFRβ$^{+}$ / KLF4$^{+}$ co-localization, conditioning on the interaction effect between DPI and Regions. We excluded sham animals because this group does not have differentiated injured cortex or perilesion areas. The model takes the following notation:

$$PDGFR\beta - KLF4 \mid trials(Klf4) \sim DPI \; * \; Region, family \; = \; binomial$$

Finally, we applied the same strategy to fit a fourth model (Mdl4) estimating the proportion of PDGFRβ$^{+}$ / KLF4$^{+}$ co-localization relative to the number of PDGFRβ$^{+}$ cells. For additional details on the modeling strategy and diagnostics, please refer to the QN. Note that the summaries of binomial models (Bernoulli trials) are presented on the logit scale. We visualized the model results using conditional_effects, displaying point estimates with 95% Cis and, when applicable, color codes for different brain regions. Model summaries are also available in the `Tables` component of the OSF repository.

**Supplementary Method 16: Analysis of KLF4 expression in PDGFRβ$^{KLF4\text{-}KO}$ mice.**

| Component | Link |
|---|---|
| Figures | Figure 7A-E |
| Raw Images | Widefield_10x_Ipsilateral_KO_Klf4-Pdgfrb-CD31(a).zip<br>Widefield_10x_Ipsilateral_KO_Klf4-Pdgfrb-CD31(b).zip |
| Low-resolution images | Widefield_10x_Ipsilateral_KO_Klf4-Pdgfrb-CD31.zip |
| Image Processing | Images_ProcessingWidefield_10x_Ipsilateral_KO_Klf4-Pdgfrb-CD31 |
| Raw Data | Data_Raw/Widefield_10x_Ipsilateral_KO_Klf4 |
| Processed data | Data_Processed/Widefield_10x_Ipsilateral_KO_Klf4 |
| Statistical models | StatisticalModels_brms/Widefield_10x_Ipsilateral_KO_Klf4_Exp |
| Analysis notebook | Widefield_10x_Ipsilateral_KO_Klf4_Exp.qmd |

We acquired widefield images at 10x magnification of KLF4$^{+}$ / PDGFRβ$^{+}$ / CD31$^{+}$ immunolabeling in brain sections (see the protocol) from the ipsilateral hemisphere of PDGFRβ$^{KLF4\text{-}KO}$ mice and corresponding controls. We included four sham-operated animals as a reference group for exploratory data visualization and to estimate error ratios. We performed pixel classification for KLF4 in Ilastik to eliminate the background and improve cell detection in CellProfiler. The generated binary masks were used as input for the IndentifyPrimeryObjects module in CellProfiler. Additionally, we detected DAPI$^{+}$ cells to establish the observation window for point patterns. We exported the data sheets and cropped images ('KLF4_Crops' folder) illustrating the identified cells.

Our initial aim was to analyze KLF4 specifically in PDGFRβ$^{+}$ cells, segregated from CD31$^{+}$ vasculature. However, antibody staining made this analysis impracticable and susceptible to multiple systematic errors. In contrast to PDGFRβ$^{tdTomato}$ cells, antibody labeling does not fill the cell nuclei expected to colocalize with KLF4. Subsequently, we processed (see QN) the CellProfiler output to extract the coordinates of individual cells and constructed point patterns using spatstat. We generated and plotted density kernels (sigma = 0.02) and performed mppm to analyze the relative distribution of KLF4 along the x-coordinates:

$$KLF4 \sim x, random \ = \sim x |\ Genotype, family \ = \ Poisson$$

We extracted the mean intensity from the point pattern hyperframe and modeled this variable using brms. We fitted an intercept-only model (Intensity ~ 1) using the data from sham animals to estimate a baseline value for KLF4 expression. Then, we fitted a linear model with heteroskedasticity with ‘genotype’ as predictor:

$$Intensity \sim Genotype, sigma \sim Genotype, family \ = \ student$$

Model diagnostics are available in the QN. We calculated the contrast between PDGFRβ$^{KLF4\text{-}KO}$ and controls using the emmeans package and presented the results as half-eye densities and point intervals. The model summary is also available in the “Tables” component of the OSF repository.

**Supplementary Method 17: Analysis of brain shrinkage and PDGFRβ / GFAP expression in PDGFRβ$^{KLF4\text{-}KO}$ mice.**

| Component | Link |
|---|---|
| Figures | Figure 7F-H; Suppl. Figure 9A-C |
| Raw Images | Widefield_5x_Ipsilateral_KO_Pdgfrb-NeuN-Gfap.zip<br>Widefield_5x_Whole_KO_Pdgfrb-NeuN-Gfap.zip |
| Low-resolution images | Widefield_5x_Ipsilateral_KO_Pdgfrb-NeuN-Gfap.zip |
| Image Processing | Images_Processing/Widefield_5x_Ipsilateral_KO_Pdgfrb-NeuN-Gfap |
| Raw Data | Data_Raw/Widefield_5x_Ipsilateral_KO_Pdgfrb-NeuN-Gfap |
| Processed data | Data_Processed/Widefield_5x_Ipsilateral_KO_Structure.csv |
| Statistical models | StatisticalModels_brms/Widefield_5x_Ipsilateral_KO_Pdgfrb-NeuN-Gfap_Structure |
| Analysis notebook | Widefield_5x_Ipsilateral_KO_Structure.qmd |

We acquired widefield images at 5x magnification of PDGFRβ$^{+}$ / NeuN+ / GFAP$^{+}$ immunolabeling in brain sections from PDGFRβ$^{KLF4\text{-}KO}$ mice and corresponding controls. We included four sham-operated animals as a reference group for exploratory data visualization and to estimate error ratios. To calculate the amount of hemispheric atrophy (shrinkage), we measured the area of the ischemic and contralateral hemispheres using the standard *measure* function in FIJI.

For the analysis of brain shrinkage, we first estimated the intrinsic area ratio (ipsilateral/contralateral) in healthy brains by fitting a statistical model with no predictors (Shrinkage ~ 1, family = student). Then, we fitted a linear model to analyze the distribution of brain shrinkage by genotype:

$$Shrinkage \sim Genotype, family = student$$

We share the model summary as .tex table in the "Tables/tex" component of the OSF repository. For the visualization of the results, we obtained posterior draws using the *spread_draws* function and generated a density plot using *stat_halfeye* with the ROPE limits in cyan.

Next, we performed pixel classification for PDGFRβ and GFAP in Ilastik to analyze the integrated density of these markers in CellProfiler. We exported binary masks from Ilastik into CellProfiler with the *MaskImage* module to measure the intensity of PDGFRβ and GFAP solely in the trained regions using the *MeasureImageIntensity* module. We exported the results into a .cvs file for statistical modeling in R. We fitted analogous linear models for PDGFRβ and GFAP to estimate the marker intensity conditioning on genotype:

$$Intensity \sim Genotype, family = student$$

We present the results using the *spread_draws* function and density plots with *stat_halfeye.* In both cases, the ROPE is shown as cyan. We share the model summaries as .tex tables in the "Tables/tex" component of the OSF repository.

**Supplementary Method 18: Analysis of PDGFRβ mRNA transcript spatial expression in the brain of PDGFRβ$^{KLF4\text{-}KO}$ mice.**

| **Component** | **Link** |
|---|---|
| Figures | Suppl. Figure 9D-F |
| Raw Images | Widefield_10x_ROIs_KO_Fish_Pdgfrb.zip |
| Low-resolution images | Widefield_10x_ROIs_KO_Fish_Pdgfrb.zip |
| Image Processing | Images_Processing/Widefield_10x_ROIs_KO_Fish_Pdgfrb |
| Raw Data | Data_Raw/Widefield_10x_ROIs_KO_Fish_Pdgfrb |
| Processed data | Data_Processed/Widefield_10x_ROIs_KO_Fish_Pdgfrb |
| Statistical models | StatisticalModels_brms/Widefield_10x_ROIs_KO_Fish_Pdgfrb |
| Analysis notebook | Widefield_10x_ROIs_KO_Fish_Pdgfrb.qmd<br>Widefield_10x_ROIs_KO_Pdgfrb_TDA.ipynb |

We explored PDGFRβ at the mRNA level using fluorescence in situ hybridization (FISH) in PDGFRβ$^{KLF4\text{-}KO}$ mice brains (see the protocol). We acquired widefield images at 10x magnification in cortical ROIs. We used the *IdentifyPrimeryObjects* module for CellProfiler to identify labelled cells and measured the morphological properties and intensities of the objects using the *MeasureObjectSizeShape* and *MeasureObjectIntensity* modules, respectively. Detected cells are available in the low-resolution component of the OSF.

To analyze the number of PDGFRβ$^{+}$ cells at 14 DPI, we fitted a multilevel negative binomial model to account for population-level effects of genotype and group-level variability due to individual differences among mice:

$$Counts \sim Genotype\ +\ (1\,|\,MouseID), family\ =\ negbinomial$$

We visualized the posterior distribution for population and group-level effects using stat_halfeye combined with geom_jitter to show actual measurements.

Next, we evaluated PDGFRβ expression intensity at the cellular level by analyzing the object-level output from CellProfiler containing area and intensity data. We fitted a mixed-effects model allowing for the interaction between cell area and genotype, using scaled predictors to improve model fit:

$$Intensity \sim Area * Genotype + (1 \mid MouseID), family = student$$

We visualized the results using conditional effects plots to display the relationship between nuclei area and PDGFRβ intensity by genotype. Both model diagnostics are available in the QN. We share the model summaries as .tex tables in the "Tables/tex" component of the OSF repository.

Finally, we performed TDA to evaluate the spatial organization of PDGFRβ$^{+}$ nuclei (see QN). First, we generated point clouds from the xy-coordinates of individual nuclei (CellProfiler output) for each animal and section. Then, we computed Vietoris-Rips complexes and persistence diagrams using the GUDHI and Ripser libraries in Python. We analyzed 0-dimensional (connected components) and 1-dimensional (loops/holes) homology features by calculating Betti curves. Finally, we aggregated these curves by genotype (PDGFRβ$^{Flox}$ vs. PDGFRβ$^{KLF4\text{-}KO}$) to compare the topological features of cell distribution. We then visualized the results as mean Betti curves with standard deviation for both dimensions **(see Suppl. Figure 9F)**.

**Supplementary Method 19: Analysis of PDGFRβ mRNA transcript spatial expression in the brain of PDGFRβ$^{KLF4\text{-}KO}$, KLF4$^{Flox}$, and PDGFRβ$^{Cre}$ mice.**

| **Component** | **Link** |
|---|---|
| Figures | Suppl. Figure 9G-H |
| Raw Images | Widefield_10x_ROIs_Flox-Cre_Fish_Pdgfrb.zip |
| Low-resolution images | Widefield_10x_ROIs_Flox-Cre_Fish_Pdgfrb.zip |
| Image Processing | Images_Processing/Widefield_10x_ROIs_Flox-Cre_Fish_Pdgfrb |
| Raw Data | Data_Raw/Widefield_10x_ROIs_Flox-Cre_Fish_Pdgfrb |
| Processed data | Data_Processed/Widefield_10x_ROIs_Flox-Cre_Fish_Pdgfrb |
| Statistical models | StatisticalModels_brms/Widefield_10x_ROIs_Flox-Cre_Fish_Pdgfrb |
| Analysis notebook | Widefield_10x_ROIs_Flox-Cre_Fish_Pdgfrb.qmd |

We analysed PDGFRβ mRNA transcript spatial expression in the brain of PDGFRβ$^{KLF4\text{-}KO}$, KLF4$^{Flox}$, and PDGFRβ$^{Cre}$ mice as described in Supplementary Method 18 (See QN). We used analogous CellProfiler outputs and conducted the same statistical modeling strategy. We share the model summaries as .tex tables in the "Tables/tex" component of the OSF repository.

**Supplementary Method 20: Analysis of Collagen IV (ColIV) in PDGFRβ$^{KLF4\text{-}KO}$ mice.**

| **Component** | **Link** |
|---|---|
| Figures | Figure 7I-L; Suppl. Figure 10A-C |
| Raw Images | Widefield_5x_Ipsilateral_KO_CD31-ColIV-Iba1.zip<br>Widefield_10x_ROIs_KO_CD31-ColIV-Iba1.zip |
| Low-resolution images | Widefield_10x_Ipsilateral_KO_CD31-ColIV.zip<br>Widefield_5x_Ipsilateral_KO_CD31-ColIV.zip |
| Image Processing | Images_Processing/Widefield_5x_Ipsilateral_KO_CD31-ColIV |
| Raw Data | DataRaw/Widefield_5x_Ipsilateral_KO_CD31-ColIV<br>Data_Raw/Widefield_10x_Ipsilateral_KO_CD31-ColIV |
| Processed data | Data_Processed/Widefield_10x_Ipsilateral_KO_CD31-ColIV |
| Statistical models | StatisticalModels_brms/Widefield_10x_Ipsilateral_KO_CD31-ColIV<br>StatisticalModels_brms/Widefield_10x_Ipsilateral_KO_CD31-ColIV_Inten |
| Analysis notebook | Widefield_10x_ROIs_KO_CD31-ColIV_Coloc.qmd<br>Widefield_10x_Ipsilateral_KO_CD31-ColIV_Inten.qmd |

We acquired widefield images at 5x and 10x magnification of CD31 / ColIV immunolabeled brain sections (see the protocol) from the ipsilateral hemisphere of PDGFRβ$^{KLF4\text{-}KO}$ mice and corresponding controls. We analized the immonolabeling intensity of vascular (CD31$^+$) basement membrane Collagen-IV (ColIV) in the whole ipsilateral hemisphere using CellProfiler outputs. First, we fitted a distributional model to estimate both the mean expression and the group-level variability (sigma) by genotype, accounting for the heterogeneity observed in the data:

$$Area \sim Genotype, sigma \sim Genotype, family = student$$

Next, we fitted a robust linear model to analyze the integrated intensity conditioning on genotype:

$$Intensity \sim Genotype, family = student$$

Model diagnostics and source code are available in the QN. We present the results as full posterior distribution contrast between genotypes using spread_draws from tidybayes). The posterior summary is shared as a .tex table in the OSF repository. Secondly, we analyzed vascular-associated Collagen-IV area in the cortex using images taken at 10x. We fitted a linear model to estimate the stained area conditioning on genotype:

$$Area \sim Genotype, family = student$$

We visualized the posterior distributions using stat_halfeye and stat_interval. We share the model summaries as .tex tables in the Tables/tex component of the OSF repository.
